# APOE4 disrupts the central dogma by arresting neuronal proteome dynamics

**DOI:** 10.64898/2026.07.15.738801

**Authors:** Einar K. Krogsaeter, Justin McKetney, Lishi Li, Isabelle Liu, Alicia L. Richards, Vishvak Subramanyam, Keyi Yin, Martin Gordon, Pedro Belio Mairal, Erica J. Stevenson, Houzhen Qian, Yadong Huang, Hani Goodarzi, Nevan J. Krogan, Danielle L. Swaney

## Abstract

Apolipoprotein E4 (APOE4) is the strongest genetic risk factor for late-onset Alzheimer’s disease and promotes neuronal dysfunction through incompletely understood mechanisms. Here, we integrated transcriptomic, translatomic, and proteomic profiling of isogenic APOE3 and APOE4 human iPSC-derived neurons and found that APOE4 fundamentally impairs neuronal proteome renewal. Although transcriptional changes were modest, APOE4 disrupted ribosome occupancy, altered translational dynamics, and uncoupled protein abundance from transcript levels. Proteome-wide turnover measurements revealed a global extension of protein half-lives and widespread accumulation of long-lived proteins. Functional proteomic analyses demonstrated concurrent lysosomal and proteasomal impairments associated with reduced proteasome activity and increased association of APOE with neuronal proteasomes. Longitudinal proteomics further showed that protein accumulation emerges during neuronal maturation and precedes a senescence-like cellular stress state. Together, these findings identify impaired proteome renewal as a central mechanism underlying neuronal vulnerability to APOE4 and establish defective proteostasis as an early pathogenic event in Alzheimer’s disease.

## INTRODUCTION

Every year, 10 million people are diagnosed with dementia. Among these, Alzheimer’s Disease (AD) represents the most common form of dementia, accounting for more than 60% of all cases.^1^ AD can be further stratified into two categories: The relatively rare early-onset AD, accounting for ∼5% of all AD cases, and late-onset AD (LOAD), accounting for the remaining ∼95%.^2^ The ε4 allele of apolipoprotein (*APOE4*) is marked by a C112R polymorphism and represents the most common risk factor for AD. In contrast to the rare genetic variants underlying early-onset AD, the *APOE4* allele is carried by 60-75% of all AD patients and homozygosity of *APOE4* increases the risk of developing AD 15-fold, as well as decreases the age of AD onset by a decade.^3,4^

The molecular mechanisms underlying early-onset AD are well-studied: rare mutations within the *APP* gene and its associated secretase complexes *PSEN1* and *PSEN2* culminate in the excessive neuronal generation of amyloid-β (Aβ) peptides and subsequent deposition of extracellular amyloid plaques, which alongside intraneuronal tau tangles represent canonical pathological hallmarks of AD. This axis has received extensive pharmacological attention, culminating in the clinical approval for Aβ-targeting antibodies. While this represents a paradigm shift in AD therapy from palliative care towards disease-modifying treatments, it comes with two caveats. First, Aβ-targeting antibodies can trigger the dangerous side-effect of Amyloid-Related Imaging Abnormalities (ARIAs), particularly in APOE4 carriers, often resulting in therapeutic discontinuation. Secondly, while such therapies dampen neuroinflammation, they do not directly restore neuronal viability. It is therefore paramount to understand how APOE4 renders neurons vulnerable in AD to develop complementary therapies which boost neuronal resilience to AD.

The role of APOE4 in AD pathogenesis is widely studied in the context of astrocytes and microglia, while its pathological effects are less characterized in neurons.^5,6^ Astrocytes produce 75-80% of the APOE within the brain,^7–9^ of which the majority is secreted.^10,11^ Astrocytic APOE4 has profound effects on cellular function, favoring lysosomal lipid accumulation^12^ and pro-inflammatory chemokine secretion.^13^ Microglia and neurons express lower levels of APOE, but upregulate its expression following cellular injury or stress-^14–18^ While homeostatic microglia produce APOE facilitate amyloid plaque clearance,^19^ in AD pathogenesis APOE4 can impair microglial lipid handling to exacerbate the cellular stresses,^20,21^ resulting in autophagic^22^ and mitochondrial defects, and increased cytokine release.^23^ Resultantly, glial APOE4 is a significant contributor to AD processes, impairing lipid and debris clearance, while driving AD neuroinflammation.

Despite comparatively low APOE expression in neurons relative to glial cells, multiple studies have demonstrated that neuronal APOE4 exerts direct effects on AD pathology and cognitive decline. Seminal work by Huang *et al*. demonstrated a causal role for neuronal APOE4 in AD pathology and cognitive decline. ^5,6^ Neuron-specific *APOE4* removal ameliorates AD-associated pathological phenotypes of the PS19/E4 tauopathy mouse model, including tau pathology, gliosis, and hippocampal atrophy.^24^ Analogously, pan-neuronal and interneuronal *APOE4* removal both reverse cognitive defects in an APOE4 mouse model of aging.^7^ Consistent with these findings, transplantation of mouse wild-type interneuron progenitors rescued age-associated cognitive decline in an APOE4 mouse model,^25^ highlighting the central role of neuronal APOE4 in cognitive deterioration.

The pathogenic influence of neuronal APOE4 can be attributed to a number of mechanisms. Neuronal APOE4 undergoes proteolytic cleavage, yielding neurotoxic protein fragments. These fragments cause neurons to accumulate cellular p-Tau inclusions,^26^ laying the foundation for the accumulation of one of the two AD hallmark aggregates. Furthermore, neuronal APOE4, especially its fragments, binds mitochondria^27,28^ and suppresses the expression of electron transport chain proteins,^29^ disrupting neuronal metabolism. Simultaneously, APOE4 neurons show impaired cytoskeletal polarization, marked by excessive PKA-mediated phosphorylation of the cytoskeletal regulator VASP.^30^ While these phenotypes have been independently attributed to APOE4, it remains unclear whether they reflect distinct pathogenic mechanisms or emerge from a shared source of cellular vulnerability. Notably, many reported APOE4 phenotypes, including protein aggregation, organellar defects, and defective cellular remodeling, suggest a broader disruption of cellular proteostasis. While we recently demonstrated that neuronal APOE4 impairs lysosomal function through the aberrant lysosomal accumulation of TMED5,^31^ a systems-level framework of neuronal vulnerabilities to APOE4 is lacking. Here, we combined transcriptomic, translatomic, and proteomic profiling of isogenic APOE3 and APOE4 iPSC-derived neurons to define the alterations of neuronal proteostasis and molecular pathways underlying neuronal vulnerability to APOE4. These integrated analyses thereby provide a systems-level framework for understanding how neuronal APOE4 reshapes proteostatic pathways to promote neuronal vulnerability in AD.

## RESULTS

### Global comparative profiling of human neurons expressing APOE3 or APOE4

To address how APOE4 renders neurons vulnerable to AD pathology, we utilized previously characterized isogenic APOE3^C112/C112^ and APOE4^C112R/C112R^ iPSC lines (from here on referred to as APOE3 and APOE4, respectively).^32^ From these iPSCs, we employed a differentiation protocol that uses small molecules for dual SMAD and Wnt inhibition alongside activation of hedgehog signaling to generate mixed neuronal cultures (Figure 1A, S1A).^32,33^ Importantly, these cultures contain a majority proportion of GABAergic interneurons that are vulnerable to APOE4 neuropathology. We confirmed that at the time of analysis (day *in-vitro* 49; DIV49), the relative proportion of interneurons was comparable between APOE3 and APOE4 cultures (Figure 1B-C, S1B-C). The mixed neuronal cultures did not yield GFAP^+^ astrocytes, an important consideration as astrocytes produce substantially more APOE than healthy neurons (Figure S1D). While a minor neuronal population exhibited nuclear Ki67 immunoreactivity indicative of immaturity and potential proliferation, this proportion was comparable between APOE3 and APOE4 neuronal cultures (Figure 1D-E). Having obtained neuronal cells that are representative of those vulnerable in APOE4 AD patients, we set out to systematically profile the intrinsic neuronal function across several molecular strata.

**Figure 1.**
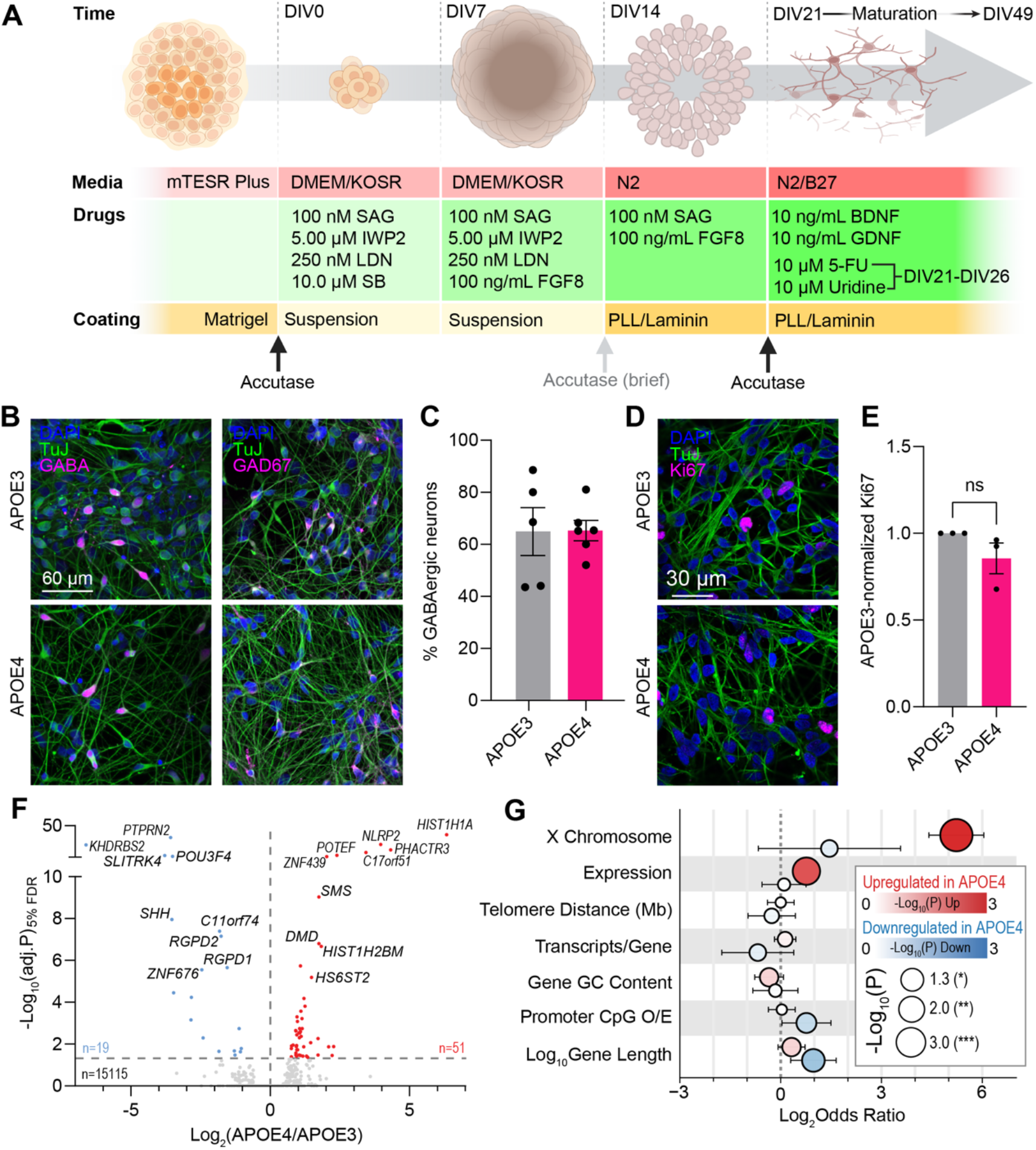
Characteristics and transcriptomics of iPSC-derived APOE4 neuronal cultures. (**A**) Schematic of how iPSCs were differentiated via embryoid body, neurosphere and neuroepithelium stages, giving rise to mixed neuronal cultures matured for 28 days before experiments were conducted. (**B** and **C**) Immunofluorescence (B) and quantification (C) of DIV35 iPSC-derived neurons, indicating total neurons (TuJ), inhibitory neurons (GABA and GAD67). n ≥ 5 biological and technical replicates from independent differentiations. (**D** and **E**) Immunofluorescence (D) and quantification (E) of DIV49 iPSC-derived neurons, indicating total neurons (TuJ) and proliferative cells (Ki67). n = 3 biological and technical replicates from independent differentiations. (F) Volcano plot comparing transcripts between APOE4 and APOE3 iPSC-derived neurons. Up- and down-regulated transcripts (adj. p<0.05) are colored red and blue, respectively. n = 3 biological and technical replicates from independent differentiations. See also Table S1. (G) Multivariable logistic regression forest plot showing effect of mRNA features on transcriptional dysregulation. X-axis position represents the odds ratio that features result in mRNA upregulation (red) or downregulation (blue), point color shades and size represent -Log_10_(P)-value of significant odds ratio enrichment, error bars represent the 95% confidence intervals.

We first compared the transcriptomes of APOE4 neurons to their healthy, isogenic, APOE3 counterparts. APOE4 neurons showed upregulation of 51 transcripts, and downregulation of 19 transcripts (Figure 1F, Table S1). We employed a multivariable logistic regression analysis to identify features associated with APOE4-associated transcriptional dysregulation, including transcriptional regulatory elements governing both general and age-associated transcription. Since females show a particularly heightened risk of developing APOE4-associated LOAD,^34,35^ we also assessed whether the APOE4 allele particularly influences transcription of X-chromosomal genes. APOE4-upregulated transcripts are enriched for both X-chromosomal genes as well as genes with high expression (Figure 1G). Inversely, APOE4-downregulated genes were enriched for longer genes and those with a higher density of CpG dinucleotides. Among the most downregulated APOE4 transcripts associated with high CpG densities included several regulators of the Sonic Hedgehog signaling pathway (*SHH*, *NKX2-4*, *LHX6*, *FOXJ1*, *RAX*, *POU3F4*) that are important for neuronal migration, maturation, and specification.

We next profiled whether dysregulated transcripts were associated with particular transcription factors (Figure S1E).^33–35^ APOE4-downregulated transcripts were associated with the SUZ12 transcription factor, which is a component of the polycomb repressive complex 2 (PRC2). PRC2 suppresses transcription by H3K27 methylation of its target genes, such as SHH signaling components.^36^ These results are consistent with a model in which APOE4-associated hypomethylation causes SUZ12 gene repression, and are aligned with previous findings of decreased DNA methylation in APOE4 AD.^37,38^ Conversely, APOE4-upregulated transcripts are associated with the transcriptional activator E2F1, targeting a number of X-chromosomal genes regulating DNA synthesis and repair (Figure S1F). Here, balancing female gene dosage with males is also strictly controlled by DNA-methylation through X-chromosome inactivation (XCi). Taken together, APOE4-associated transcriptional dysregulation is linked to methylated gene elements, including suppression of autosomal PRC2-linked genes and increased transcription of X-linked genes.

### Translational decoupling and ribosomal stalling in APOE4 neurons

While transcriptional regulation lays the foundation for differential gene expression, the translation of protein is necessary to generate active effector protein molecules from mRNA blueprints. In general, we found that translation rates were comparable across APOE genotypes via puromycin incorporation assays (Figures S2A-B). However, we observed that a subset of APOE4 neurons (17%) were refractory to puromycin incorporation as compared to APOE3 neurons (9%; Figure S2C), suggesting that APOE4 neurons may harbor a greater fraction of translationally inactive cells. For a more targeted analysis of APOE4-associated translational changes, we performed ribosome footprinting (Ribo-seq). In general, higher mRNA levels were associated with more ribosome-protected fragments (RPF) in both APOE3 and APOE4 neurons (Figures 2A-B, Table S2).

**Figure 2.**
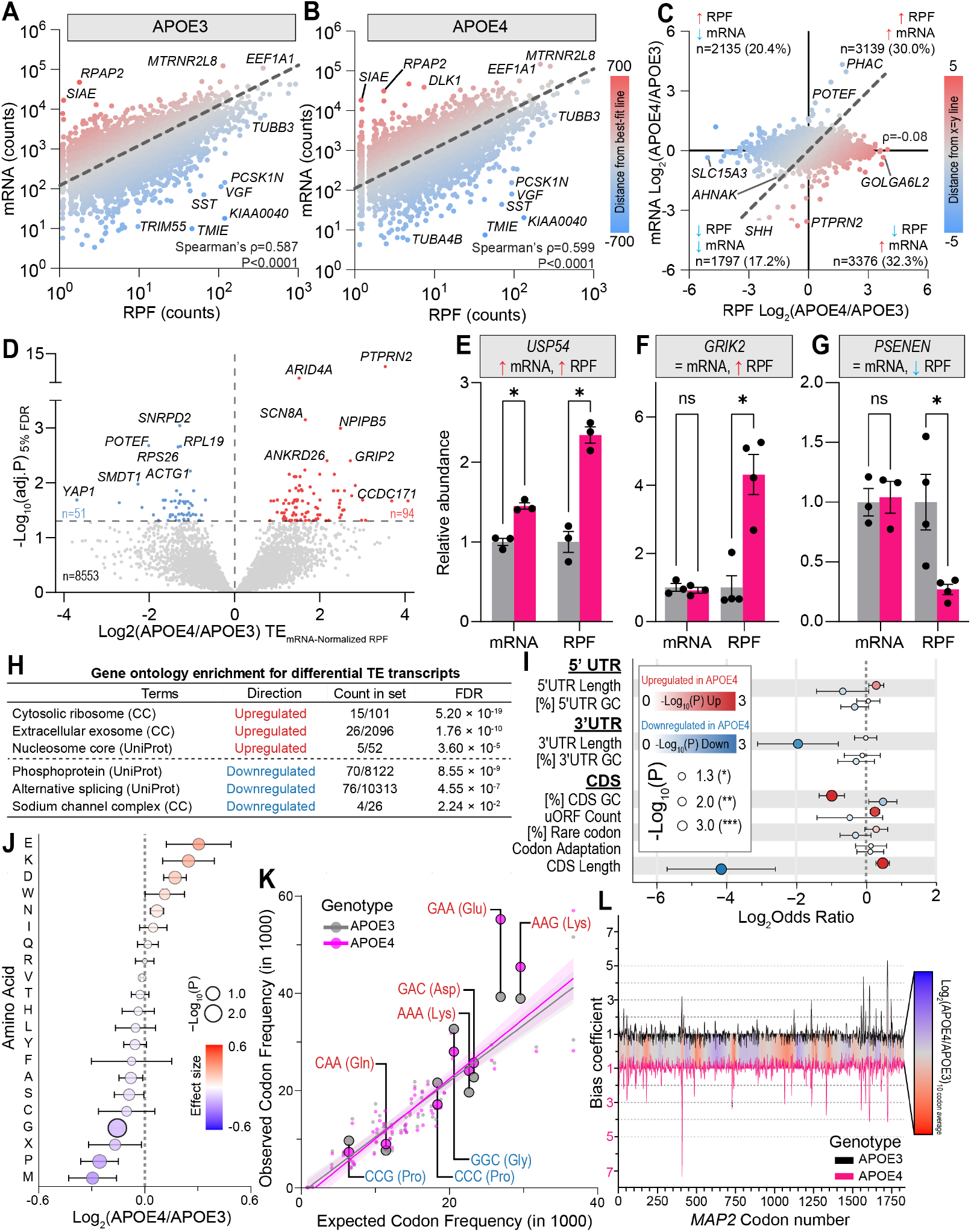
Ribosomal translation is decoupled from transcript abundance in APOE4 neurons. (**A**-**B**) Correlation of mRNA abundance and RPF in APOE3 (A) and APOE4 (B) neurons. Dashed line at x=y shows the best-fit line for each genotype. Color represents transcript-translation decoupling score, where red indicates more mRNA relative to RPF, and blue indicates less mRNA relative to RPF. n = 4 biological and technical replicates from independent differentiations. See also Table S2A. (C) Comparison of APOE4-associated changes in mRNA abundance and RPF. Dashed line at x=y shows expected region where changes in transcript levels are reflected in altered RPF levels. Color represents decoupling score, red indicates higher ribosomal occupancy relative to mRNA, and blue indicates lower ribosomal occupancy relative to mRNA. n = 4 biological and technical replicates from independent differentiations. See also Table S2B. (D) Volcano plot comparing translational efficiency (TE; RPF:mRNA ratio) between APOE4 and APOE3 iPSC-derived neurons. Genes of increased and decreased TE are colored red and blue, respectively. N = 4 biological and technical replicates from independent differentiations. See also Table S2C. (**E**-**G**) Example bar plots for concordance (E) and discordance (F-G) between transcriptional and translational changes in APOE4 neurons relative to APOE3 controls. n = 4 biological and technical replicates from independent differentiations. (H) Gene-set enrichment of transcripts that undergo TE changes in APOE4 neurons, based on a ranked STRING^39^ enrichment analysis. Low-FDRs GO terms were selected to avoid semantic redundancy. (I) Forest plot of features associated with transcripts that have altered TE in APOE4 neurons. X-axis position represents the odds ratio that features result in TE increase (red) or decrease (blue), color shades represent effect sizes. Point sizes represent the -Log_10_(P)-value of significant odds ratio enrichment, error bars represent the 95% confidence intervals. CAI: Codon adaptation index, IRES: Internal ribosome entry site, uORF: Upstream open reading frame, UTR: Untranslated region. (J) Amino acids associated with Ribo-seq-retrieved codons in APOE4 relative to APOE3 neurons. Black border data points indicate p<0.05, with inset asterisks indicating significance degree. Error bars indicate SEM. n = 4 biological and technical replicates from independent differentiations. See also Table S2D. (K) Comparison between expected *H.sapiens* codon frequency (x-axis) and observed codon frequency in iPSC-derived neurons (y-axis). Codons significantly differing between APOE4 and APOE3 neurons are highlighted in red (more in APOE4) and blue (more in APOE3). Data points were fitted by linear regression giving rise to indicated best-fit lines and 95% confidence intervals of the regression line. n = 4 biological and technical replicates from independent differentiations. See also Table S2E. (L) Codon bias coefficients along the *MAP2* transcript by APOE genotype. Top black line shows APOE3 codon bias, while the mirrored magenta line shows APOE4 codon bias. Inset heat-map indicates the average of Log_2_(APOE4/APOE3) codon bias values as a 10-codon rolling average. n = 4 biological and technical replicates from independent differentiations.

We could also observe cases of transcriptional decoupling. For example, the somatostatin (*SST*) neurotransmitter mRNA was overrepresented in ribosomes relative to available transcript, reflecting continuous synaptic protein synthesis to sustain interneuron signaling (Figures 2A-B).^40^ In contrast, we found that two genes (*ZNF560* and *PCDHB10*) were subject to the inverse dysregulation between APOE3 and APOE4 neurons, where APOE4-increased mRNA and decreased RPF transcript levels were observed (Figure S2D). Most genotype-associated decoupling events occurred as RPF changes in the absence of mRNA changes (Figures 2C, S2E). Normalization of RPF values by mRNA transcript levels to calculate translational efficiency (TE, Figure 2D) revealed that while some transcripts such as *USP54* showed concordant transcriptional-translational upregulation (Figure 2E), others such as *GRIK2* and *PSENEN* exhibited ribosomal occupancy changes in the absence of mRNA alterations (Figures 2F-G).

To highlight patterns underlying TE changes, we performed a gene set enrichment analysis (GSEA) of transcripts with APOE4-sensitive translational efficiencies. APOE4 decreased the TE of ribosome, exosome, and nucleosome transcripts, and increased the TE of transcripts associated with phosphoproteins and alternative splicing (Figure 2H). We next assessed mRNA features associated with decreased translational efficiency. The strongest predictor of low TE was mRNA coding sequence (CDS) length (Figure 2I), consistent with the possibility that longer transcripts provide more opportunities forribosomal pausing during elongation.^41^ Because codon composition influences translation elongation through differential decoding and tRNA utilization,^42–44^ we compared codon occupancy between APOE4 and APOE3 ribosomes (Figure 2J). APOE4 ribosomes showed decreased occupancy of glycine (GGC) and proline (CCC, CCG) codons (Figure 2K), consistent with faster decoding of glycine and proline codons, which are established determinants of translational pausing. Importantly, glycine- and proline-associated pausing events facilitate co-translational protein folding,^45^ thereby maintaining proteome integrity. In contrast, APOE4 ribosomes exhibited increased occupancy at a subset of codons encoding glutamate (GAA), lysine (AAG, AAA), aspartate (GAC), and glutamine (CAA; Figure 2K). Notably, four of the five enriched codons were adenosine-rich triplets, suggesting that APOE4 selectively alters ribosome transit through A-rich coding contexts. To test whether we could observe stalling along neuronal transcripts, we analyzed regional ribosomal pausing along *MAP2* (Figure 2L). Although we did not observe changes in ribosomal stalling associated with gene length or position, we found that codon bias coefficients associated with *MAP2* were more variable in APOE4 neurons, consistent with an uneven ribosome distribution along the transcript. Importantly, unresolved translational stalling is a causative driver for neurodegeneration in the case of Charcot-Marie-Tooth disease.^46,47^ In AD, similar stalling events occur during co-translational translocation of the pathogenic APP C99 fragment, resulting in deposition of aggregation-prone APP species that nucleate Aβ plaque formation.^48^ These results identify altered ribosome transit and codon-specific pausing as a potential mechanism linking APOE4 to neuronal translation changes in AD.

### APOE4 neurons extensively accumulate long-lived proteins

We next continued along the central dogma to examine how APOE4 influences the neuronal proteome.^38–47^ Contrasting our transcriptomic and translational observations, we observed a biased distribution of fold-changes, in which proteins were generally more abundant in APOE4 neurons (Figures 3A, S3A, Table S3). We performed a weighted gene network correlation analysis, finding that 4 of 28 protein co-expression modules were affected by the APOE4 genotype, impacting mitochondrial and ribosomal proteins (Figure 3B). We identified several features significantly associated with protein accumulation in APOE4 neurons, including long half-lives, high baseline expression, high isoelectric points, low molecular weight, and mitochondrial or synaptic localization (Figure 3C). The strongest associated feature of dysregulation was baseline protein expression, where low-abundance proteins tended to be downregulated and abundant proteins tended to be upregulated in APOE4 neurons (Figure 3D). We also found that proteins significantly more abundant in APOE4 neurons were enriched for low molecular weight proteins (Figure 3E). This contrasts the enrichment of long-CDS transcripts by Ribo-seq, which would encode high molecular weight proteins. Taken together, these results suggest that in APOE4 neurons, long CDS-transcripts are inefficiently translated into large proteins.

**Figure 3.**
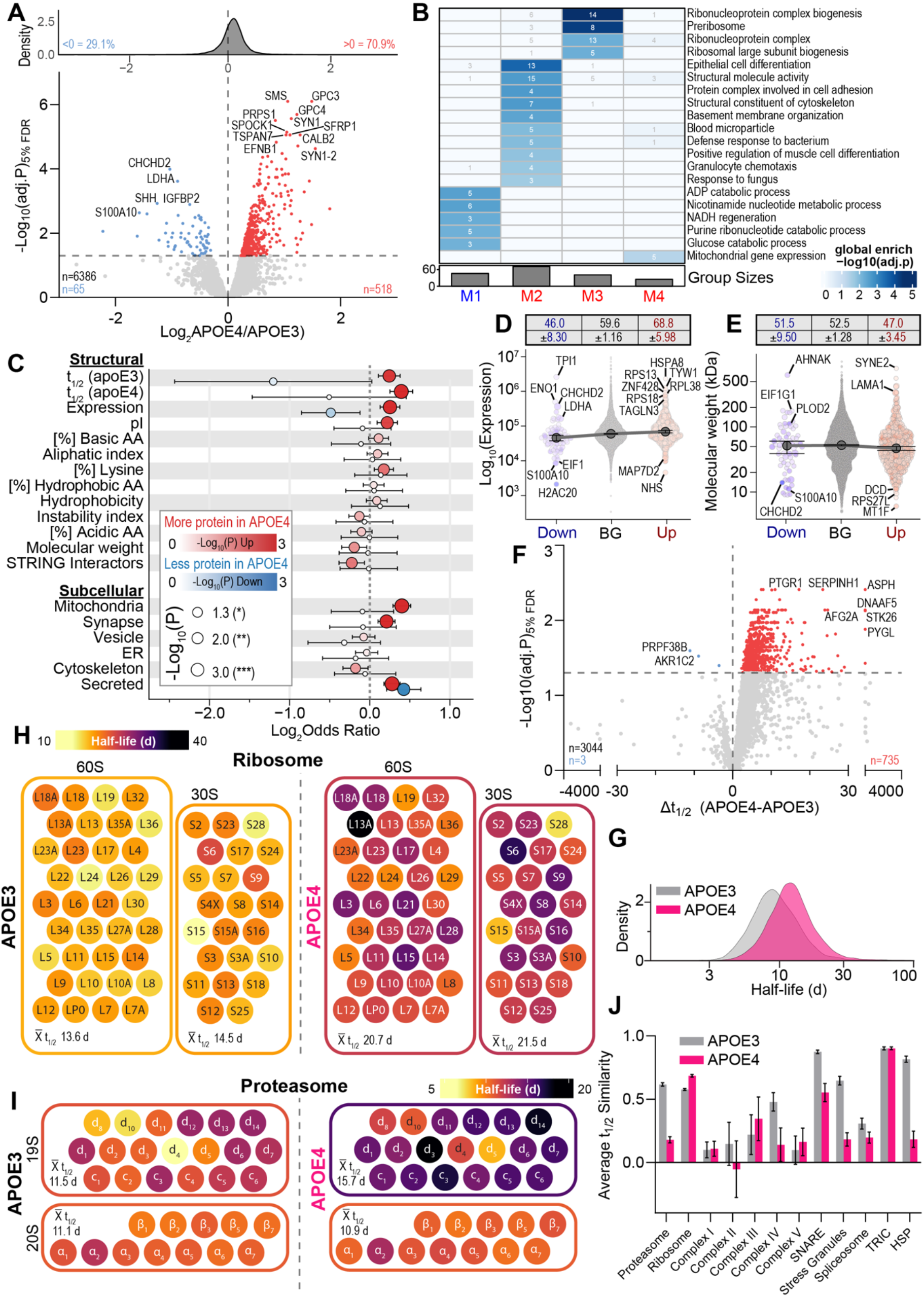
APOE4 causes widespread accumulation of long-lived neuronal proteins. (A) Volcano plot comparing protein abundances between APOE4 and APOE3 iPSC-derived neurons. Up- and down-regulated data points (adj. p<0.05) are colored red and blue, respectively. Overlaid density plot shows a positive skew of the total protein distribution. n = 6 biological and technical replicates from independent differentiations. See Table S3. (B) APOE4-sensitive co-expression modules based on weighted gene co-expression network analysis: One downregulated (M1, blue), and three upregulated (M2-4, red) modules. Cell shade indicates -Log_10_P values, inset numbers indicate the number of dysregulated genes per cell. (C) Forest plot of features associated with altered protein abundance in APOE4 neurons. X-axis position represents the odds ratio that features result in protein upregulation (red) or downregulation (blue), color shades represent effect sizes. Point sizes represent the -Log_10_(P)-value of significant odds ratio enrichment, error bars represent the 95% confidence intervals. AA: Amino acids, ER: Endoplasmic reticulum, pI: Isoelectric point. (D) Scatter plot of Log_10_-scaled protein expression level of all proteins (background, BG), upregulated (up, red) and downregulated (down, blue) protein sets. Large circles indicate population medians, error bars indicate the 95% confidence interval. Data point color intensity indicates Log_2_(APOE4/APOE3) values, circle size indicates statistical significance. n = 6 biological and technical replicates from independent differentiations. (E) Scatter plot of protein molecular weight of all proteins (background, BG), upregulated (up, red) and downregulated (down, blue) protein sets. Large circles indicate population medians, error bars indicate the 95% confidence interval. Data point color intensity indicates Log_2_(APOE4/APOE3) values, circle size indicates statistical significance. n = 6 biological and technical replicates from independent differentiations. (F) Volcano plot comparing protein half-lives between APOE4 and APOE3 iPSC-derived neurons. Proteins with significantly extended or shortened half-lives (adj. p<0.05) are colored red and blue, respectively. N ≥ 5 biological and technical replicates from independent differentiations. See Table S4A. (G) Density plot of average protein half-lives reported in days (d) in APOE3 and APOE4 neurons. See Table S4A. (H) 2D schematic of the protein half-lives reported in days (d) of 60S large and 30S small ribosomal subunit members from APOE3 (left) and APOE4 (right) neurons. Inset X^-^ values indicate mean protein half-lives in days (d) within the respective sub-complexes. (I) 2D schematic of the protein half-lives reported in days (d) of 20S catalytic and 19S regulatory proteasomal subunit members from APOE3 (left) and APOE4 (right) neurons. Inset X^-^ values indicate mean protein half-lives in days (d) within sub-complexes. (J) Average protein cosine similarity values based on protein half-lives within indicated protein complexes in APOE3 (gray) and APOE4 (magenta) neurons. High similarity values indicate complex co-regulation. Error bars indicate standard error of the mean. n ≥ 5 biological and technical replicates from independent differentiations. See Table S4B.

The observed proteomic shift toward protein accumulation in APOE4 neurons could be the result of a fundamental failure in protein homeostatic pathways. To further investigate this possibility, we complemented our snapshot of protein abundance by measuring how APOE4 affects the life-span of neuronal proteins by pulsed Stable Isotope Labeling by Amino acids in Cell culture (pSILAC).^49,50^ We treated neurons with heavy lysine for two weeks to obtain equivalent heavy:light peptide ratios (Figure S3B), and estimated protein half-lives (t_1/2_) assuming first-order synthesis/turnover kinetics. Similar to the global shift toward higher protein expression, we found that 88% of all detected proteins trended towards longer half-lives in APOE4 neurons (Figure 3F, Table S4), with the median protein half-life increasing from 8.9 days in APOE3 neurons to 12 days in APOE4 neurons (Figure 3G). The impacted proteins included several risk genes for AD (APP, CLU, CTSB, PICALM) and other NDDs,^51^ including NDDs whose progression is modulated by APOE4 (Figure S3C).^52,53^

Gene-set enrichment of proteins with significantly prolonged turnover rates^39^ revealed that proteins persisting in APOE4 cells were particularly enriched for regulators of gene expression and translation (n=134, FDR=1.39*10^-26^), in line with the ribosomal protein accumulation in APOE4 neurons (Figure 3B). While the entire ribosomal complex has a half-life of 14 days in APOE3 neurons, this extends to 20 days in APOE4 neurons (Figure 3H). Intriguingly, the half-lives of ribosomal subunits are very uniform within ribosomal complexes per genotype. This is consistent with the major ribosomal degradative process, ribophagy, in which assembled ribosomal complexes are turned over in unison.^54^ Proteasomal complexes contrast this observation. In APOE3 neurons we observe a similar half-life of both the catalytic 20S subunit and the regulatory 19S subunit that suggests a coordinated turnover of assembled proteasomes. In contrast, while the mean half-life of 20S subunit proteins in APOE4 neurons resembles that of APOE3 neurons (11 days), we observe a decoupling in turnover with the 19S subunit resulting in an extended mean half-life (16 days) in APOE4 neurons (Figures 3I, S3D). A similar proteasome subunit decoupling was recently observed in human dorsolateral prefrontal cortex tissue from AD patients, where 19S proteasomal subunits exhibited decreased correlation in protein abundance relative to healthy controls.^55^ We similarly profiled half-life-associated subunit decoupling by comparing cosine similarity scores of protein turnover within well-defined cellular complexes, finding evidence for APOE4-associated complex uncoupling in the spliceosome, HSP70-HSP90 co-chaperone, and stress granule protein complexes (Figures 3J, S3E). Taken together, these results indicate that APOE4 not only promotes protein longevity, but also disrupts the orchestrated turnover of specific protein complexes.

### Synergistic protein clearance defects determine the APOE4 neuronal proteome

To evaluate potential synergies between the transcriptional, translational, and proteomic landscapes, we integrated our RNA-seq, Ribo-seq, and proteomics data. We found that overlap in significantly regulated genes/proteins was rare (Figure 4A, Table S5) with only 20 genes/proteins being dysregulated at both the transcriptional and proteomic level. Overall, most cases of dysregulation occurred uniquely at the protein level (83.3% of all dysregulated genes/proteins) and were associated with protein accumulation (Figure 4B). This was particularly evident for several of the most dysregulated proteins, including ribosomal proteins such as RPL11 and synaptic proteins such as GPC4 and SYN1 (Figure 4C). These findings indicate that APOE4 disrupts the natural flow of the central dogma. Typically, DNA transcription into RNA and the subsequent translation by ribosomes into proteins dictates the cellular effectors in play. In APOE4 neurons, we observe a decoupling between transcriptomic and proteomic changes, resulting in an APOE4-associated bias towards protein accumulation. We hypothesized that this proteomic decoupling may be the result of inefficient protein clearance, which has previously been shown to be impacted early in AD.^31,56–60^

**Figure 4.**
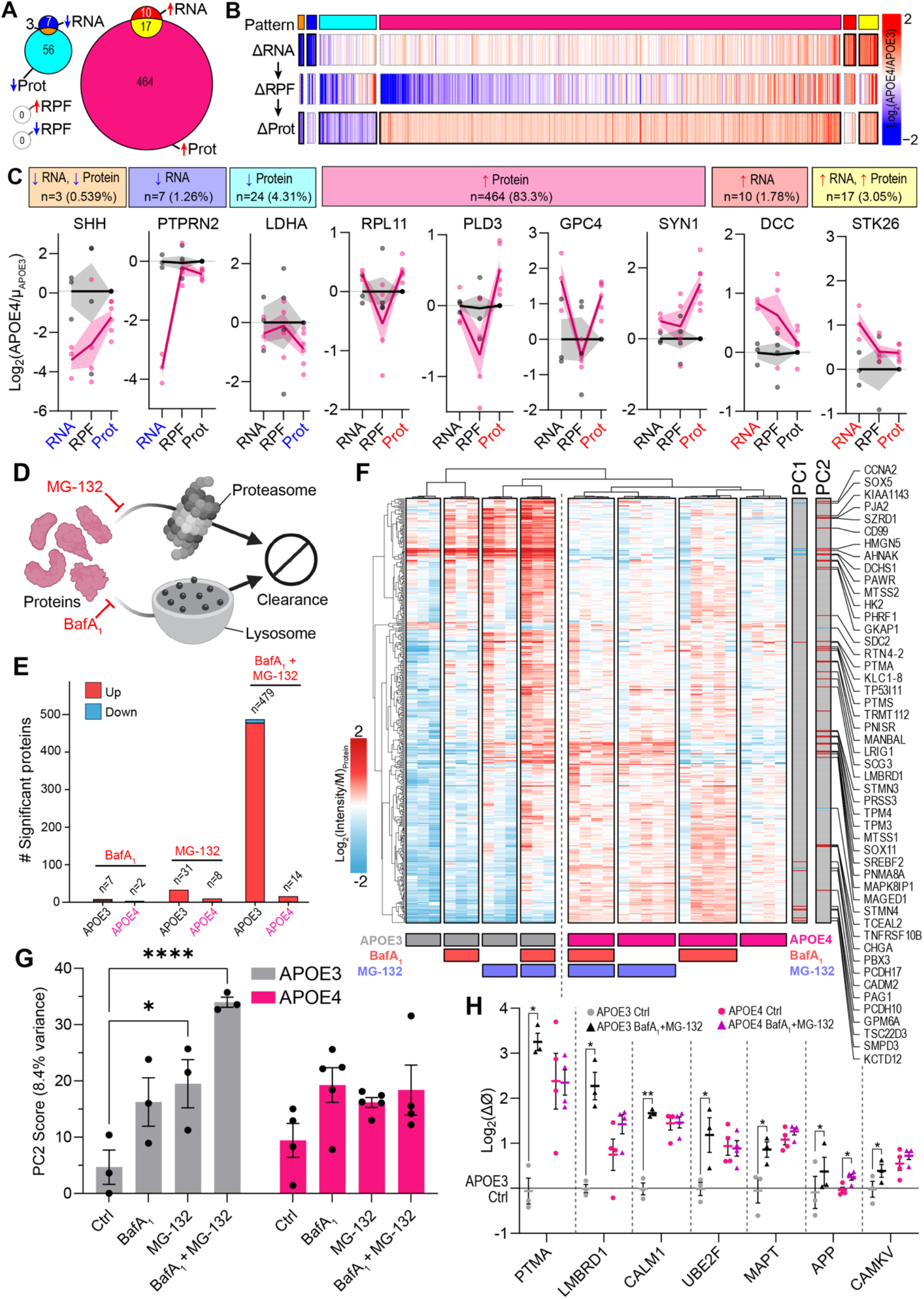
Global proteomic dysregulation in APOE4 neurons is driven by clearance defects. (A) Venn diagram of significantly dysregulated genes across RNA, RPF, and protein layers (adj. p<0.05), filtered for complete data and dysregulation in ≥1 layer. See Table S5. (B) Heat-map integrating RNA-seq, RPF, and proteomic changes in APOE4 relative to APOE3 neurons. Protein rows were filtered for complete data with significant dysregulation (adj. p<0.05) occurring in one of the RNA, RPF, or protein abundance levels, clustered based on the associated dysregulation, and ordered based on average extent of dysregulation. See Table S5. (C) Examples of cross-layer protein dysregulation, with dysregulation patterns indicated in overhead boxes. Inset numbers indicate gene count affected by the associated dysregulation. X-axis label font color denotes whether the associated metric was significantly increased (red) or decreased (blue) in APOE4 neurons (adj. p<0.05). Data points represent measurements from separate differentiations, line indicates measurement mean, shaded area indicates standard error of the mean. (D) Schematic indicating how blockade of proteasomal or lysosomal degradation by MG-132 or BafA_1_ for 3 hours, respectively, increases protein abundance (pink), measured by mass-spectrometry. (E) Bar plot of protein abundance changes upon clearance inhibition relative to DMSO-treated controls, filtered for proteins where adj. p<0.05 and |Log_2_FC|>0.58. n ≥ 3 biological and technical replicates from independent differentiations. See Table S6A. (F) Heat-map of proteins whose expression changed upon clearance inhibition. Colors indicate the median-scaled Log_2_(Intensity) per protein. Top 2% proteins (ranked by z-score) associated with the primary principal components (PC) are highlighted. (G) Bar plot showing the influence of clearance inhibition and APOE genotype on PC2 score across differentiation batches. n ≥ 3 biological and technical replicates from independent differentiations. See Table S6B. (H) Dot plot showing abundance changes of highlighted proteins by APOE genotype and clearance inhibition. n ≥ 3 biological and technical replicates from independent differentiations. See Table S6C.

To interrogate what pathway(s) may underly APOE4-associated protein accumulation, we systematically blocked protein clearance mechanisms and used mass spectrometry to measure the acute impact on the proteome (Figure 4D). We set out to measure changes in protein degradation by APOE4 by treating iPSC-derived neurons for 3 hours with BafA_1_ (inhibiting lysosomes) and/or MG-132 (inhibiting proteasomes) and measuring changes to the neuronal proteome. Each treatment individually had a modest impact on the proteome with a similar number of impacted proteins in both the APOE3 and APOE4 neurons (Figure S4A, Table S6). In contrast, simultaneous inhibition of proteasomal and lysosomal protein clearance synergistically resulted in protein accumulation in APOE3 cells beyond what can be expected of additive clearance inhibition, while protein accumulation was much less evident in APOE4 neurons (Figures 4E, S4A). This is likely the result of degradative compensation across the autophagic and proteasomal systems, where inhibition of one pathway is buffered by increasing the activity of the other.^61–63^ In contrast, when both pathways are inhibited simultaneously, this compensation is disrupted, resulting in the observed dramatic protein accumulation. We performed a principal component analysis (PCA) to visualize differential effects of clearance inhibitors on the neuronal proteome (Figures S4B-C). PC1 explained 68.4% of the observed variance (Figure S4B) and was primarily associated with the APOE genotype, while PC2 (8.4% of variance) was most associated with clearance blockade. Accordingly, while high PC2-scoring proteins reproducibly accumulated in APOE3 neurons upon blocking lysosomal and proteasomal degradation, the same proteins appeared resilient to clearance inhibition in APOE4 neurons (Figures 4G). By analyzing the clearance-sensitive proteome, we found that several of the proteins accumulating by clearance blockade in APOE3 neurons were elevated at baseline in the APOE4 context (Figure 4H, S4D). This particularly affected synaptic proteins (CALM1, CAMKV), but also extended to the central AD hallmark proteins amyloid precursor protein (APP) and Tau (MAPT; Figure 4H). These results indicate that synergistic clearance defects affecting both the lysosomal and proteasomal systems lay the foundation for APOE4-associated proteomic changes, both affecting neuronal connectivity and favoring the protein accumulation of aggregation-prone proteins associated with AD.

### APOE4 binds neuronal proteasomes and disrupts proteasomal function

While we^31^ and others^12,64^ previously described APOE4-associated lysosomal impairments and associated lysosomal retention of APOE4,^31,65^ the observed simultaneous inhibition of proteasomal and autophagic pathways by APOE4 is unexplored. We first queried our steady-state proteomics data to see if the expression of proteasomal subunits was changed by APOE4. We found that all proteasome components were slightly elevated in APOE4 neurons, with the exception of the PA28 immunoproteasome subunits PSME1 (PA28α) and PSME2 (PA28β) that appeared slightly less abundant (Table S3). We validated this by immunofluorescence, finding no overt change in the expression, colocalization, and distribution of the catalytic 20S PSMA and PSMB5 subunits, besides slightly lower PSMA association with PSMB5 (Figures 5A, S5A). In contrast, we confirmed that APOE4 neurons have a slightly decreased abundance of the regulatory immunoproteasome subunit PSME1 (PA28α), in particular in neurites (Figure 5B-C). Intriguingly, expression of the PA28αβ proteasome regulator complex supports cognitive function through maintaining protein clearance capacity upon aging.^66^ PA28αβ responds to oxidative stress in neurons to facilitate degradation of damaged, oxidized proteins, and was recently associated with tau proteolytic processing in neurons.^67,68^ The APOE4-associated neuronal downregulation of PA28αβ therefore is expected to lay the foundation for oxidative damage and associated accumulation of oxidatively damaged proteins. This has important implications in AD, where oxidative damage, alongside endocytic defects, constitutes a hallmark event in disease initiation.^69^

**Figure 5.**
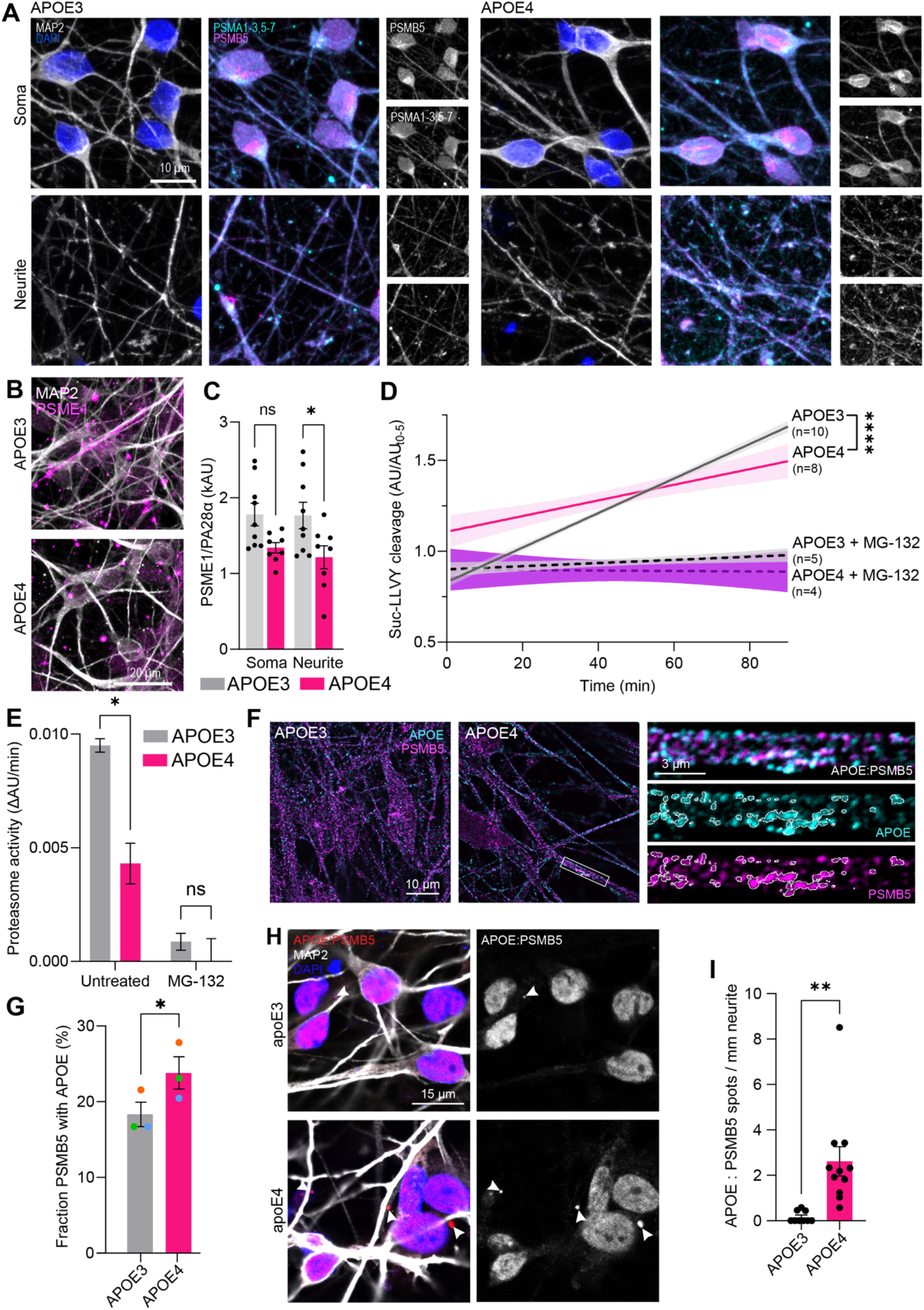
Proteasomal impairments in APOE4 neurons are associated with proteasomal APOE. (**A**) Immunofluorescence of catalytic 20S proteasome subunits (PSMB5 and PSMA1-3, 5-7) distribution in iPSC-derived neuronal neurites and soma. (**B** and **C**) Immunofluorescence (B) and quantification (C) of immunoproteasome subunit PSME1 expression levels in DIV49 iPSC-derived neuronal neurites and soma. n = 3 biological and technical replicates from independent differentiations. (**D** and **E**) Proteasome activity assay of fluorogenic Suc-LLVY cleavage (A) by APOE3 (black) and APOE4 (magenta) lysates ± the proteasome inhibitor MG-132 (dashed lines). Shaded regions denote standard error of the mean. Inset n numbers represents biological replicates from independent differentiations. Proteasomal degradation rates were quantified by linear regression. n ≥ 4 biological replicates from independent differentiations. (**F** and **G**) Co-immunofluorescence of 20S subunit PSMB5 and APOE in DIV49 iPSC-derived neurons by confocal microscopy and deconvolution (F). The zoomed-in neurite region (right panels) is indicated by a rectangular white box, highlighting APOE colocalization with PSMB5 along the neurite. Colocalization quantification, colored data points showing matched replicates (G). n = 3 biological and technical replicates from independent differentiations. (**H** and **I**) Proximity ligation assay between APOE and PSMB5 in iPSC-derived neurons, co-stained for MAP2 and nuclei (DAPI). White arrowheads indicate sites of proximity ligation signal (H). Quantification (I) of fluorescent PLA puncta indicative of proteasomal APOE (APOE:PSMB5-based proximity ligation) was normalized to neurite length. Each data point represents a field-of-view from one of three separate biological replicates per genotype. Background PLA signal in nuclei was excluded from the analysis. n = 3 biological and technical replicates from independent differentiations.

Since we could not observe conclusive evidence that the widespread proteasomal defects were caused by changes in the expression or distribution of the essential proteasomal subunits, we continued to assess whether proteasomal activity is influenced by the APOE allele status. We found that APOE4 neurons exhibit significantly lower proteasomal activity than their APOE3 counterparts, with APOE4 neuron lysates degrading the fluorogenic proteasome substrate Suc-LLVY 54.7% slower than corresponding APOE3 lysates (Figures 5D-E, S5B-C). As we had previously observed that APOE4-associated lysosomal defects were associated with accumulation of lysosomal APOE,^31^ we moved to test whether we could observe a similar association between APOE4 and proteasomes. This notion was particularly prompted by recent observations that the neuroproteasome, a specialized neuronal plasma-membrane proteasome, specifically binds to and is functionally regulated by APOE4.^70^ We found APOE to be distributed in puncta along neurites, with several of these puncta colocalized with the proteasomal subunit PSMB5, and that this colocalization was increased in APOE4 neurons (Figure 5F-G). To validate these findings by a more sensitive approach, we performed a proximity ligation assay (PLA) to measure the colocalization of APOE and PSMB5, bearing in mind the caveat that the PLA is best suited to profile known protein-protein interactions, requiring interactors to be within 40 nm of each-other. Nonetheless, we observed significantly more PLA signal between APOE and the proteasomal subunit PSMB5 in APOE4 neuronal cultures (Figures 5H-I, S5D), validating that APOE4 neurons show increased association of APOE with proteasomes.

### Post-mitotic dysregulation of the APOE4 proteome results in a neuronal senescence-like cell state

Having established clearance-associated proteomic defects in APOE4 neurons associated with concurrent lysosomal and proteasomal impairments, we considered a central unresolved question in the field: When does AD develop? This question was raised briefly following the initial disease description, spurred by observations that histopathological alterations occur in pre-symptomatic individuals.^71–73^ The pre-clinical phase was further described by efforts to stage the disease.^74^ This led to the observation that AD pathology begins in the twenties (around 40 years before the initial clinical diagnosis),^75,76^ associated with mild cognitive impairments.^77^ APOE4 modifies the pre-clinical disease phase, accelerating Aβ deposition and NFT formation by shifting the age of onset of the initial pre-clinical disease insult 12 years earlier.^78^ While molecular changes preceding classical AD pathology have been described,^79,80^ the detailed cellular events driving pre-clinical disease development, and how APOE4 shapes these events, are just recently being explored.^6^ We analyzed global protein expression from the iPSC-derived neurons each week over the course of the differentiation to assess if APOE modifies neuronal differentiation, and to pinpoint at what time we first observe neuronal proteome dysregulation, in particular relation to the expression of APOE4.

We defined DIV0 samples as iPSCs, DIV7-DIV21 (embryoid body, neurosphere, and neuroepithelium stages) as neuronal progenitor cells (NPCs), and plated DIV28-DIV49 cultures as post-mitotic neurons. We first identified markers associated with neuronal differentiation by highlighting clusters that were differentially regulated over time and associated with the aforementioned differentiation stages. Expectedly, we found a high correlation between differentiation lineage markers at the associated differentiation stages. Intriguingly, APOE itself appeared among the iPSC stage-enriched proteins (Figure S6A). We next assessed whether the APOE4 allele altered the differentiation trajectory, analyzing the temporal expression profiles of well-described stage markers per genotype. We did not see robust changes in the differentiation time-course in APOE4 samples, with stage markers being associated with their respective stages in both APOE3 and APOE4 samples (Figures 6A, S6B-D). We did however notice that stage markers persisted for longer in APOE4 cultures, and that particularly synaptic proteins showed an accelerated upregulation during the neuronal differentiation stage (Figure 6A). To identify the stage where neuronal proteins become dysregulated, we mapped their APOE4-associated dysregulation over time, and clustered similarly behaving proteins (Figure 6B). This yielded 7 dysregulated protein clusters that behaved in distinct ways, but all appeared normal during the iPSC stage. The largest and smallest protein clusters (C1, C7) both showed a gradual elevation of ribosomal, nucleolar, mitochondrial and vesicular proteins, starting at DIV14 and persisting to the end of the differentiation. Two APOE4-elevated protein clusters (C2, C5) showed a delayed timeline of protein accumulation, impacting synaptic proteins from DIV28 onwards. Uniquely, one protein cluster (C4) showed a progressive decline in vesicle-associated proteins during the neuronal phase starting at DIV28. Finally, the postsynaptic protein cluster (C6) was elevated during the NPC stage and upregulated by APOE4 from DIV14 until DIV42, after which the expression of their associated proteins was decreased. Taken together, our results indicate that the observed proteomic dysregulation of APOE4 neurons manifests as early as the neurogenesis phase, while APOE4-associated synaptic protein accumulation is particularly associated with the post-mitotic neuronal stage. While this aligns with APOE expression (higher during early differentiation; Figure 6C), we do not observe proteomic dysregulation in iPSCs, where APOE expression is the highest. We therefore postulate that the pathological effects of APOE4 are inversely associated with cell proliferation, wherein the proteome of highly proliferative iPSCs is governed by the proteomic rejuvenation through nascent protein synthesis. Meanwhile, the proteomes of NPCs and neurons appears to be dictated by an equilibrium of protein synthesis and protein degradation. APOE4 expression thus first appears pathogenic as cell division is halted and proteins are allowed to persist. Finally, we assessed whether APOE4 affects the expression trajectories of other AD-associated proteins, finding that the expression profiles of clusterin (Figure 6D), doublecortin (Figure 6E), and synapsin (Figure 6F) are all modified in APOE4 neurons.

**Figure 6.**
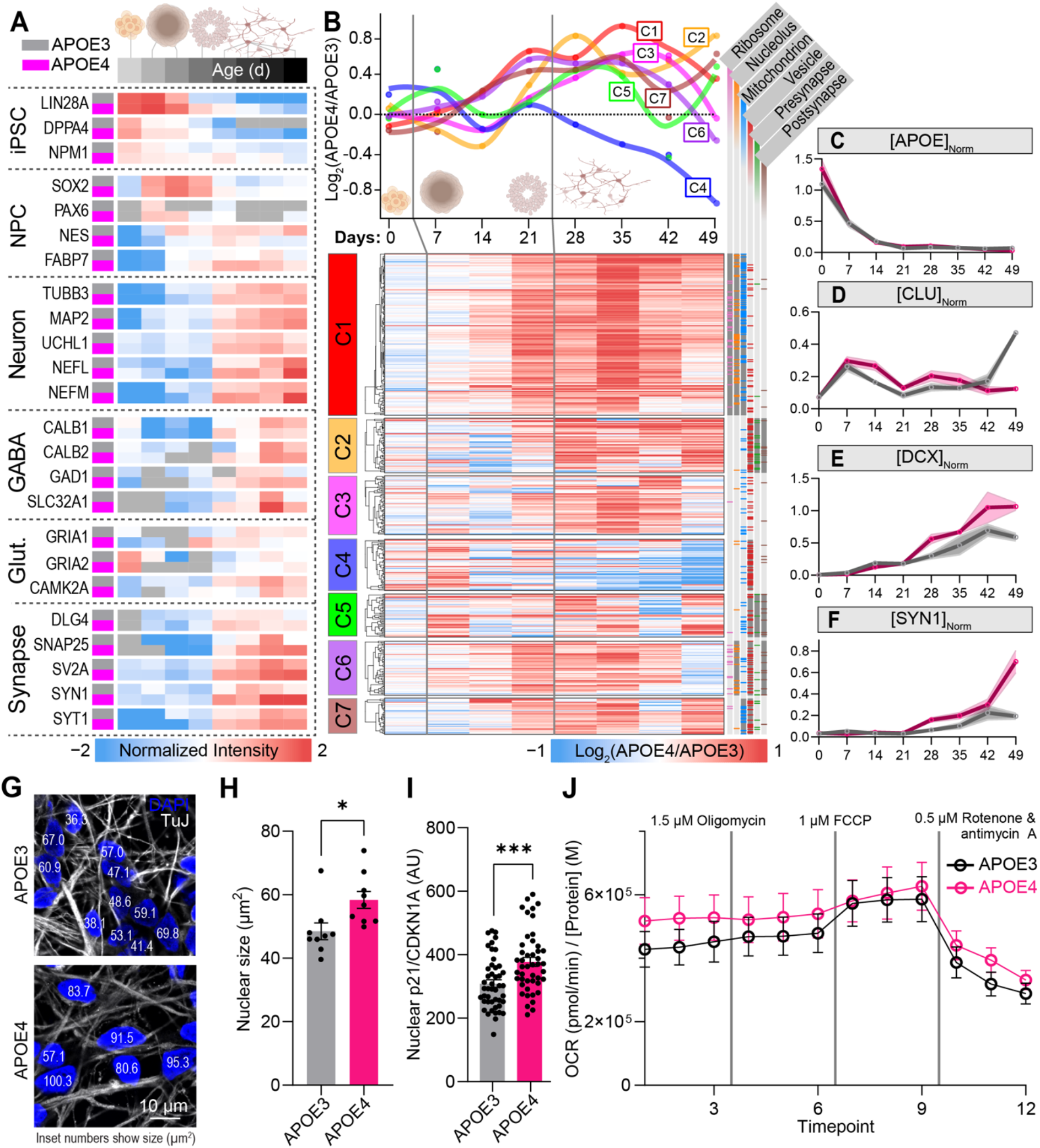
Longitudinal proteomics reveal early protein accumulation preceding senescence. (A) Average protein expression of differentiation-stage markers associated with neurogenesis, scaled to genotype-pooled expression minimum (blue) and maximum (red), centered on the median expression level for each protein. n ≥ 3 biological and technical replicates from independent differentiations. See Table S7A. (B) Time-course of neuronal protein dysregulation normalized to GAPDH and PPIA expression. Dysregulated proteins were clustered based on their co-dysregulation (APOE4 versus APOE3) over time. Top line plot shows LOESS-smoothed mean cluster trajectories over time. Lower heat-map shows individual APOE4/APOE3 protein changes over time, where colors represent Log_2_(APOE4/APOE3) values. Right-adjacent annotation lines indicate cellular compartment gene-set enrichments per cluster as identified using STRING,^39^ dark columns indicating statistically significant enrichment per cluster. n ≥ 3 biological and technical replicates from independent differentiations. See Table S7B. (**C**-**F**) Example trajectories of select proteins, indicating protein expression across differentiation stages in APOE3 (grey) and APOE4 (magenta) cells, expressed as a fraction relative to GAPDH/PPIA housekeeping genes. Shaded regions indicate the standard error of the mean. n ≥ 3 biological and technical replicates from independent differentiations. (**G** and **H**) Nuclear immunofluorescence (G) and average quantification (H) of Hoechst- and TuJ-stained DIV49 iPSC-derived neurons. White numbers within nuclei indicate the nuclear size (μm^2^). n = 8 biological and technical replicates from independent differentiations. (I) Quantification of the senescence marker CDKN1A/p21 in DIV49 neurons. n = 3 biological and technical replicates from independent differentiations. See also Figure S6F. (**J**) Mitochondrial stress test of DIV49 neurons showing change of oxygen consumption rates (OCR) as a proxy for aerobic respiration in iPSC-derived neurons, normalized to total protein amounts. n = 3 biological and technical replicates from independent differentiations. See also Figures S6G-I.

Several of the aforementioned molecular changes suggest that APOE4 neurons favor a senescence-like cell state. Traditionally, senescence represents a cellular stress response, epitomized by cell cycle arrest in the G_1/0_ phase. Nonetheless, the existence of neuronal senescence has been appreciated for half a decade, particularly in the context of AD.^81^ Recently, the molecular basis of neuronal senescence has started to be unveiled. Aged and diseased neurons become refractory to external stimuli, express senescence-associated proteins, and suffer from DNA damage, impaired oxidative phosphorylation, and oxidative stress.^82–84^ As these events are tightly associated with AD proteinopathy,^85,86^ a hypothesis of proteinopathy-induced neuronal senescence has been formulated as a proposed mechanism for AD development.^87^ We therefore continued to assess whether APOE4 neurons exhibited senescent phenotypes. This possibility was supported by an unexpected observation during the puromycin incorporation assay, where we found that APOE4 neurons lacked a population of tetraploid/G_2_-phase cells, as revealed by Hoechst counter-staining (Figure S6E). We orthogonally assessed complementary markers for neuronal senescence. First, we performed a retrospective analysis of nuclear size,^88^ and found that nuclei from APOE4 neurons are 20.2% larger than APOE3 nuclei (Figures 6G-H), a similar effect size as what has previously been reported for AD neurons (20.6%).^88^ We also observed a modest increase in transcript levels of the senescence-associated *CDKN1A*/p21 in APOE4 neurons (1.91-fold), while other senescence-associated transcripts (*CDKN2A*/p16, *CDKN2D*/p19)^83^ did not differ by APOE genotype (Table S1). By immunofluorescence, we confirmed that nuclear CDKN1A protein levels were elevated in APOE4 neurons (Figures 6I, S6F). We finally tested whether the senescent phenotype of APOE4 neurons extended to altered metabolic activity. In particular, the spare respiratory capacity of neurons is abolished upon cellular senescence.^89^ In line with this, while we did not observe changes in basal respiration nor non-mitochondrial oxygen consumption, we found that the spare respiratory capacity (following FCCP treatment) was reduced in APOE4 neurons (Figures 6J, S6G). We also found a trend toward higher glutamine/pyruvate respiratory dependency in APOE4 neurons (Figure S6H). In contrast, the glycolytic rates were not impacted by the APOE genotype, with equal extracellular acidification rates (ECAR) between APOE3 and APOE4 cells (Figure S6I). Taken together, these results indicate that APOE4 neurons suffering from impaired proteostasis enter a senescence-like stress state, impacting respiratory adaptation to neuronal stress.

## DISCUSSION

While APOE4 is the strongest risk factor for late-onset AD, the molecular basis predisposing APOE4 neurons to neurodegeneration remains incompletely understood.^5,6^ Here, by integrating transcriptomic, translatomic, and various proteomic profiling approaches, we identify impaired proteome renewal as a unifying mechanism underlying APOE4-associated neuronal vulnerability. While APOE4 showed modest influences on the neuronal transcriptome, its impact became increasingly pronounced at successive stages of the central dogma. Ribosomal occupancy was uncoupled from transcript abundance by APOE4 and was associated with stalling events previously attributed to AD.^48^ Simultaneously, we found that APOE4 neurons show decreased protein clearance, extending the half-lives of neuronal proteins and culminating in widespread protein accumulation. These findings suggest that neuronal vulnerability to APOE4 may not be primarily driven by altered gene expression programs, but is rather governed by post-transcriptional mechanisms.

A growing body of evidence indicates that decoupling of transcription, translation, and protein homeostasis represents a central hallmark of aging and neurodegeneration.^90–93^ This has significant implications for neurons, which are highly sensitive to proteostatic defects as they represent the most long-lived post-mitotic cell-type of the human body. Unlike proliferative cells, neurons cannot dilute damaged proteins through cell division, and instead rely on continuous proteomic maintenance by protein quality-control systems. In line with this framework, we observed a global extension of protein half-lives in APOE4 neurons, accompanied by accumulation of highly expressed and long-lived proteins. Notably, the median protein half-life was 35% longer in APOE4 neurons, suggesting that APOE4 fundamentally slows neuronal proteome renewal. These findings establish impaired protein turnover as a dominant molecular consequence of neuronal APOE4 expression.

Our data indicate that APOE4-asscoiated proteostatic defects arise through simultaneous disruption of the two principal cellular degradative pathways. Lysosomal defects have emerged as an early, initiating event in AD pathogenesis.^58,94–98^ We recently demonstrated that APOE4 disrupts lysosomes by functionally rewiring the lysosome, including lysosomal TMED5 accumulation.^31^ Here, we corroborate and extend these observations by demonstrating that the APOE4 proteome resembles the proteomic state influenced by simultaneous lysosomal and proteasomal blockade in APOE3 neurons. Importantly, blockade of these pathways showed negligible proteomic effects in APOE4 cells, suggesting that APOE4 neurons suffer from substantial clearance defects already at baseline. Taken together, our findings show that APOE4 causes proteostatic insufficiency through the synergistic impairment of lysosomal and proteasomal degradation pathways.

Lysosomal dysfunction has previously been linked to APOE4 and AD pathology, largely owing to significant genetic enrichment of risk factors for neurodegeneration in the endolysosomal pathway.^99–101^ In comparison, the influence of genetic risk factors for AD on proteasomal function is less well understood. Pioneering studies have, however, demonstrated that proteasomes are both impacted by, and functionally important in AD pathogenesis.^59,60,102,103^ We found that APOE4 neurons exhibit reduced proteasomal activity, independent of subunit expression levels. These observations are particularly intriguing in the context of recent results demonstrating functionally relevant interactions between exogenous APOE4 and neuronal membrane-associated proteasomes.^70^ We similarly find that neuronal APOE4 colocalizes with neuronal proteasomes, particularly within neurites. Supporting a functional influence of APOE4 on proteasomes, protein turnover measurements revealed coordinated turnover of proteasome complexes in APOE3 neurons, whereas APOE4 selectively extended the half-life of the 19S regulatory complex with little effect on the 20S core particle, suggesting a loss of coordinated proteasome homeostasis in the neuronal APOE4 context. While the mechanistic consequences of this interaction should be addressed further, our findings support a model in which neuronal APOE4 impairs proteasomal function through altering its regulation rather than expression. Of note, it remains unclear whether the increased association between APOE4 and proteasomes reflects enhanced recruitment of APOE destined for degradation or a direct inhibitory interaction contributing to proteasomal dysfunction.

Through longitudinal proteomics, we found that the differential impact of APOE4 begins as early as the neuronal progenitor stage and is more tightly associated with decreasing proliferation rates than the high APOE expression during the iPSC stage. This observation suggests that the neuronal pathogenic consequences of APOE4 are not only determined by APOE abundance but also by the cellular capacity to renew the proteome. As proliferation rates decline, APOE4-associated defects in protein turnover become progressively unmasked, resulting in widespread accumulation of long-lived proteins and disruption of protein homeostasis. Notably, many molecular phenotypes emerging during this transition resemble established hallmarks of cellular aging, including impaired proteostasis, ribosomal protein accumulation, and reduced metabolic flexibility. Consistent with this framework, APOE4 neurons exhibited enlarged nuclei, elevated p21 expression, and reduced spare respiratory capacity, indicative of a stressed senescence-like cell state. Together, these findings suggest that APOE4 accelerates biological aging of neurons by impairing proteome renewal and drives neurons toward a premature senescence-like state preceding the emergence of overt neurodegenerative pathology.

Impaired proteome renewal provides a mechanistic framework linking neuronal APOE4 to AD and neurodegeneration. Rather than influencing individual disease-associated proteins, we find that APOE4 slows the kinetics of neuronal proteome maintenance. The broad extension of protein half-lives observed in APOE4 neurons is not only restricted to AD-associated proteins but also extends to proteins implicated across neurodegenerative disorders including ALS, FTD, PD, and lysosomal storage disorders. Since long-lived proteins accumulate pathological modifications such as persistent aberrant post-translational modifications (PTMs), oxidation, isomerization, and misfolding, slowing their turnover lays the foundation for diverse proteinopathies. These findings therefore indicate that beyond the direct, causative role of APOE4 in AD, APOE4 potentially modifies the course of other neurological proteinopathies, positioning impaired proteome renewal as a fundamental determinant of neuronal vulnerability during aging.^104–107^

## Limitations of the study

This study has several limitations warranting further investigation. First, transcriptomic analyses were performed using bulk RNA sequencing of mixed neuronal cultures, which may mask neuronal cell-type-specific responses to APOE4. Future single-cell approaches will be required to determine whether distinct neuronal populations differ in their susceptibility to impaired proteome renewal and will facilitate direct comparison with single-cell data from mouse models and human AD brain tissues. Second, the experiments were performed in iPSC-derived neurons in culture, which incompletely recapitulate the aged cellular environment in which AD develops. Validation in aged human neurons, transdifferentiated neurons, and *in vivo* chimeric mouse models would therefore be a valuable future complement to this. Third, all experiments were conducted using female-derived isogenic cell lines, and sex-dependent effects of APOE4 remain to be explored. Fourth, although we identify proteasomal dysfunction and increased APOE-proteasome association in APOE4 neurons, the specific proteasome assemblies affected by APOE4 and the molecular consequences of this interaction remain unresolved. Fifth, while our data reveal concurrent defects in protein clearance and translation, the mechanistic relationship between these processes remains unclear. Future studies directly perturbing lysosomal and proteasomal function while monitoring ribosomal dynamics will be required to determine whether impaired protein clearance contributes to the translational defects observed in APOE4 neurons. Finally, we only explored APOE4 effects on proteome renewal in neurons, future studies should also be done in glial cells with much higher levels of APOE4 expression to obtain a more comprehensive view of the effect of APOE4 across different types of cells within the brains.

## RESOURCE AVAILABILITY

### Lead contact

Further information and requests for resources and reagents should be directed to and will be fulfilled by the lead contact, Danielle Swaney.

## ACKNOWLEDGMENTS

We thank Robert W. Mahley, Ryan Corces, James S. Fraser, Hanna Martens, Ken Nakamura, Maria Barna, Kirsten Obernier, Robyn Kaake, Zelda Love and Michael Ward for helpful discussions. We also thank Iris Avellano, Antara Rao, Oscar Yip, Tushar Raskar, Anastasia Zhurukhina, Neal Bennett, Siyu Chen, Austin Holub, Benjamin J. Polacco, Michael McGregor, and Ronald Babu for methodological support. We are grateful for support with lab infrastructure by Margaret Soucheray and Holli Deval, and the rest of the UCSF Quantitative Biosciences Institute (QBI) administrative personnel. We thank the Gladstone Flow Cytometry, Histology & Light Microscopy (HLMC), and Stem Cell Cores (SCC), and the UCSF Center for Advanced Technology (CAT), for equipment support and technical assistance. This work was supported by the California Institute for Regenerative Medicine (CIRM; Scholars Program to E.K.K.), the UCSF Weill Institute for Neuroscience (UCSF Catalyst Program to E.K.K. and D.L.S.), and the National Institute on Aging (NIA; R01AG092454 to D.L.S.; R01AG085357 to D.L.S.; R03AG091095 to D.L.S.; R01AG071697 to Y.H.; R01AG076647 to Y.H.).

## AUTHOR CONTRIBUTIONS

Conceptualization, E.K.K. and D.L.S.; methodology, E.K.K., J.M., L.L., A.L.R., D.L.S.; Investigation, E.K.K., J.M., L.L., I.L., A.L.R., V.S., K.Y., M.G., P.B.M., E.S., H.Q., D.L.S.; writing—original draft, E.K.K. and D.L.S.; writing—review & editing, E.K.K., Y.H., H.G., D.L.S.; funding acquisition, E.K.K., Y.H., H.G., N.J.K., D.L.S.; resources, H.G., N.J.K., D.L.S.; supervision, Y.H., H.G., D.L.S.

## DECLARATION OF GENERATIVE AI AND AI-ASSISTED TECHNOLOGIES

During the preparation of this work, the author(s) used ChatGPT 5.5 and Claude Sonnet 4 for review of literature and ensure consistency of written protocols, and to prepare code for data analysis and graphical representation of transcriptomic and proteomic datasets. After using this tool or service, the authors reviewed and edited the content as needed and take full responsibility for the content of the publication.

## SUPPLEMENTAL INFORMATION

### Document S1. Figures S1–S6

**Table S1. RNA-seq results from DIV49 APOE3 and APOE4 neurons** Contains raw RNA-counts, processed RNA counts, and downstream transcriptomic analysis results obtained from DIV49 APOE3 and APOE4 neurons, associated with Figure 1.

**Table S2. Ribo-seq results from DIV49 APOE3 and APOE4 neurons** Contains raw RPF-counts, processed RPF counts, and downstream ribo-seq analysis results obtained from DIV49 APOE3 and APOE4 neurons, associated with Figure 2.

**Table S3. Abundance proteomics results from DIV49 APOE3 and APOE4 neurons** Contains raw protein intensities, normalized protein intensities, and downstream abundance proteomic analysis results obtained from DIV49 APOE3 and APOE4 neurons, associated with Figures 3A-E.

**Table S4. Turnover (pSILAC) proteomics results from DIV49 APOE3 and APOE4 neurons** Contains raw heavy/light protein intensities, heavy/light ratios of each protein, calculated protein half-lives, and downstream pSILAC proteomic analysis results obtained from DIV49 APOE3 and APOE4 neurons, associated with Figures 3F-J.

**Table S5. Omics integration results from DIV49 APOE3 and APOE4 neurons** Contains integrated data across modalities and downstream analysis results obtained from DIV49 APOE3 and APOE4 neurons, associated with Table S1-S3.

**Table S6. Clearance inhibition proteomics results from DIV49 APOE3 and APOE4 neurons** Contains raw protein intensities, normalized protein intensities, and downstream abundance proteomic analysis results obtained from DIV49 APOE3 and APOE4 neurons treated with protein clearance inhibitors, associated with Figures 4A-F.

**Table S7. Longitudinal proteomic results from APOE3 and APOE4 neuronal differentiations** Contains raw protein intensities, normalized protein intensities, and downstream abundance proteomic analysis results obtained from DIV0-DIV49 APOE3 and APOE4 neurons, associated with Figure 6.

## STAR★METHODS

### KEY RESOURCES TABLE

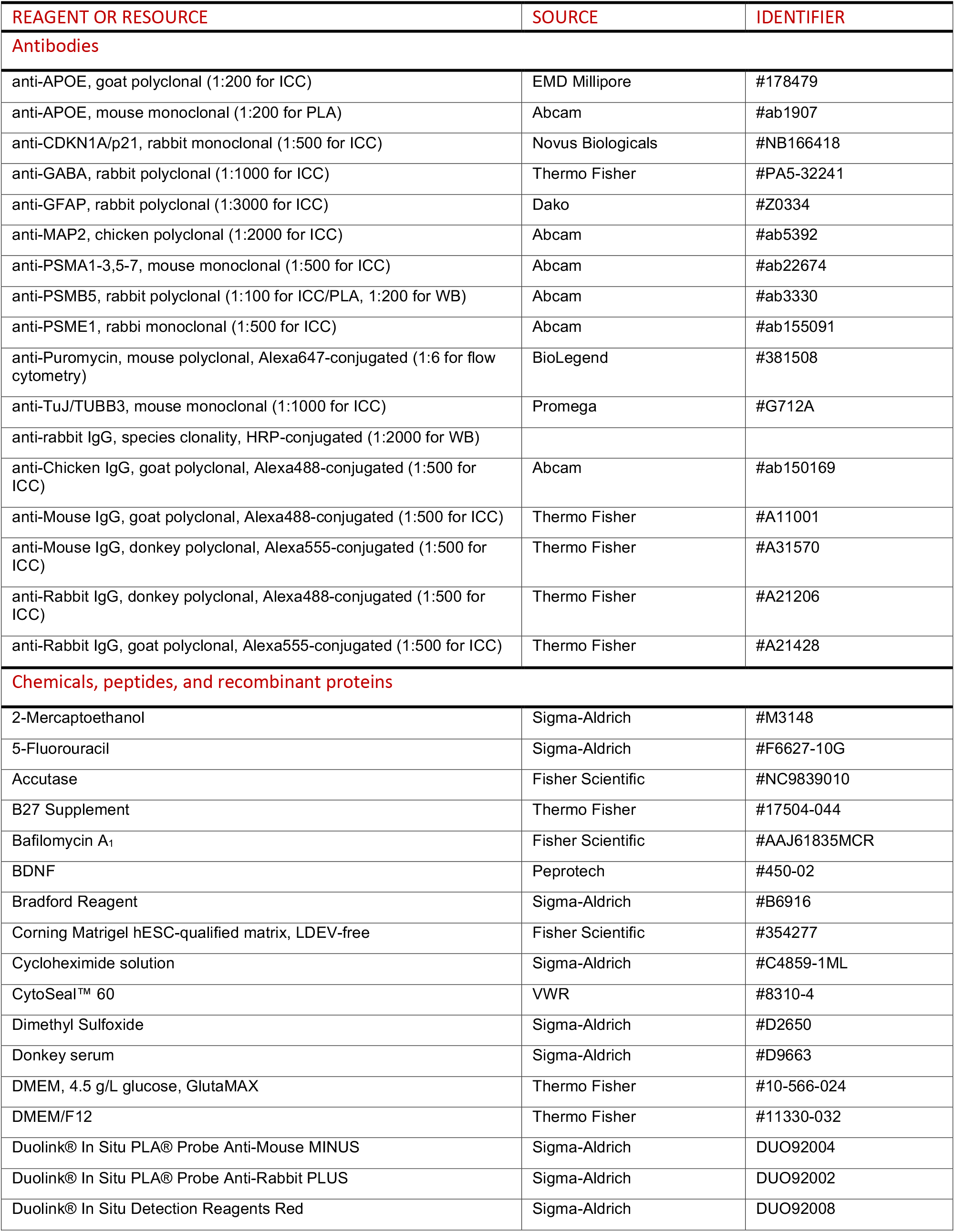
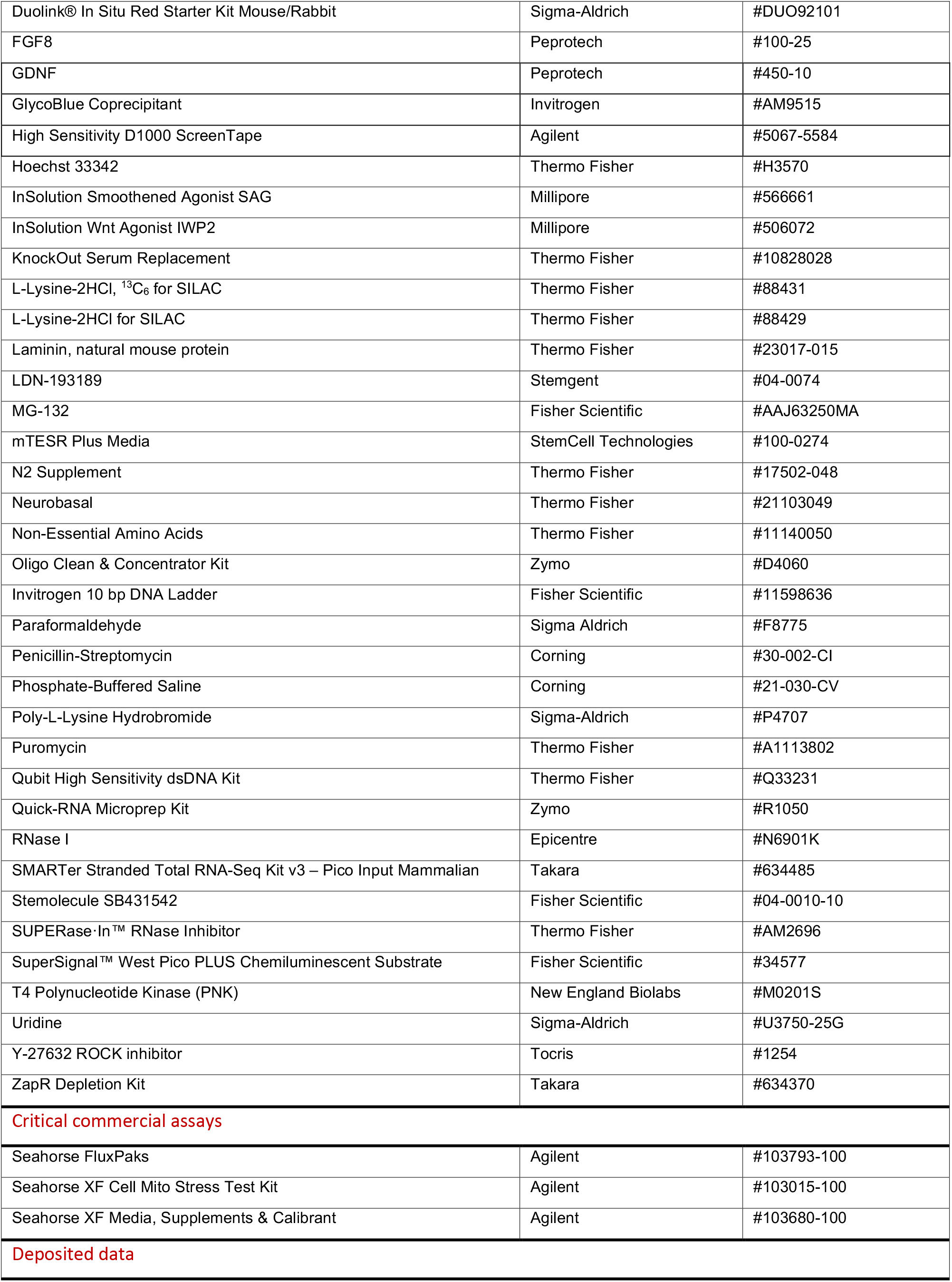
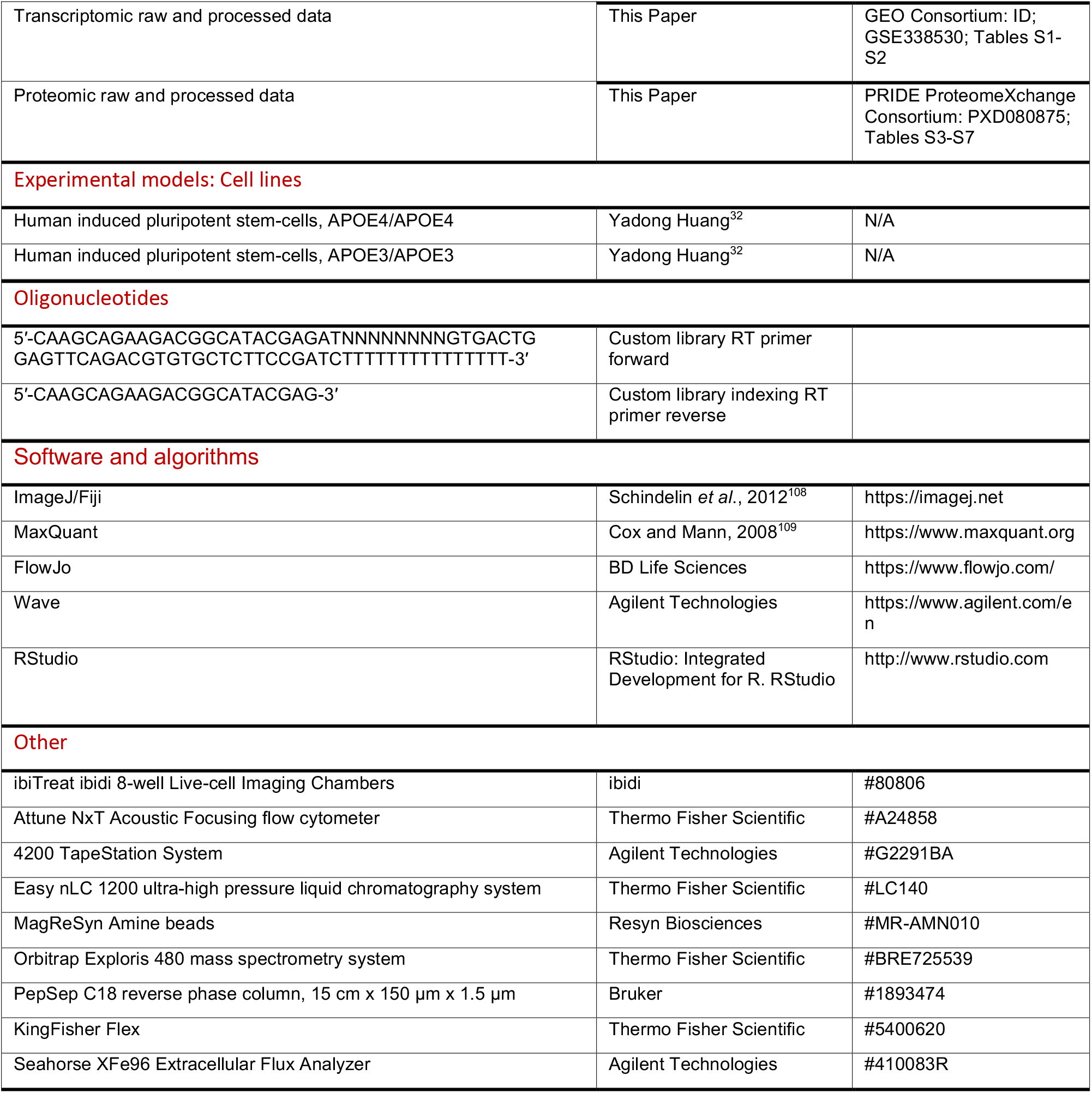

## METHOD DETAILS

### EXPERIMENTAL MODEL AND STUDY PARTICIPANT DETAILS

#### Induced pluripotent stem cells

Human induced pluripotent stem cells (iPSCs) were obtained from the laboratory of Yadong Huang (UCSF/Gladstone Institutes), as previously described.^32^ In short, skin fibroblasts were obtained from a 64-year-old female donor homozygous for APOE4 with confirmed Alzheimer’s disease. These were reprogrammed by retroviral transduction of Oct3/4, Sox2, Klf4, and c-Myc to obtain induced pluripotent stem cells.^110^ Isogenic homozygous APOE3 iPSCs were generated by ZFN-mediated gene editing, followed by monoclonal expansion and subsequent quality control (targeted sequencing of *APOE*, *APP*, *MAPT*, *BACE1*, *PSEN1*, and the potential off-target *NPLOC4*, karyotype, immunofluorescence, teratoma formation). iPSCs were maintained on Matrigel-coated plastic under feeder-free conditions using mTESR Plus medium, and routinely passaged: Accutase was added for 5 minutes, cells were gently triturated in DMEM/F12, centrifuged at 400g for 4 minutes, and seeded onto Matrigel at 1:2-1:6 with 10 µM Y-27632 ROCK inhibitor. All cells used herein were cultured in a humidified incubator kept at 37°C with 5% CO_2_.

#### SMALL MOLECULE-MEDIATED DIFFERENTIATION OF HUMAN IPSC-DERIVED NEURONS

Mixed neuronal cultures were generated as previously described with minor modifications.^31,32^ A graphical representation of the differentiation protocol is provided in Figure 1A. Briefly, iPSCs from a confluent 100-mm dish were dissociated with Accutase (4 mL, 5 min), collected by centrifugation (200 g, 2 min), and resuspended in 15 mL knockout serum replacement (KSR) medium consisting of 80% DMEM, 20% knockout serum replacement, 1X non-essential amino acids (NEAA), 1X GlutaMAX, 0.5X penicillin-streptomycin, and 10 μM β-mercaptoethanol. During the first week of differentiation (DIV0–7), KSR medium was supplemented with 10 μM SB-431542, 250 nM LDN-193189, 5 μM IWP2, and 100 nM smoothened agonist (SAG) (KSR-I). Additionally, 10 μM Y-27632 was included from DIV0–1. Cells were transferred to suspension culture and maintained as neurospheres. At DIV1, neurospheres were collected by centrifugation, resuspended in fresh KSR-I lacking Y-27632, and transferred to T175 flasks. Cultures were fed every other day with KSR-I until DIV7. From DIV7–14, neurospheres were maintained in KSR-II, consisting of KSR medium supplemented with 250 nM LDN-193189, 5 μM IWP2, 100 nM SAG, and 100 ng/mL FGF8. At DIV14, neurospheres were partially dissociated with Accutase, collected by centrifugation, and replated in N2 medium containing DMEM/F12 supplemented with 0.5X N2, 1X NEAA, 1X GlutaMAX, 0.5X penicillin-streptomycin, 100 ng/mL FGF8, and 100 nM SAG. Cells from each differentiation were distributed across two poly-L-lysine/laminin-coated 100-mm dishes and maintained until DIV21 with half-medium changes every other day. At DIV21, neuroepithelial cultures were dissociated to single cells using Accutase, filtered through a 40-μm cell strainer, and counted. Cells were seeded at 200,000 cells/mL onto poly-L-lysine/laminin-coated culture vessels for downstream applications. Neurons were maintained in N2/B27 medium consisting of 50% DMEM/F12, 50% Neurobasal, 0.5X N2 supplement, 0.5X B27 supplement, 1X NEAA, 1X GlutaMAX, and 0.5X penicillin-streptomycin supplemented with 10 ng/mL BDNF and 10 ng/mL GDNF. During the first week following neuronal plating (DIV21–28), cultures additionally received 5 μM 5-fluorouracil and 5 μM uridine to suppress proliferating cells. Neurons were maintained by twice-weekly half-medium changes and used for experiments at DIV49 unless otherwise indicated.

#### IMMUNOFLUORESCENCE AND CONFOCAL MICROSCOPY

For microscopic analysis, 200,000 iPSC-derived neurons were seeded per 12 mm poly-L-lysine-coated coverslip for confocal microscopy, and cultured as previously described to DIV49 for microscopy analysis. Cells were briefly washed once with PBS before they were fixed with 4% paraformaldehyde for 15 minutes at room temperature. Cells were washed twice with PBS and stored in PBS with 0.1% NaN_3_ at 4°C before blocking and permeabilization.

To detect differentiation markers (TUBB3/TuJ, GABA, GAD67, GFAP, Ki67, MAP2), the neurons were washed once with washing buffer (DPBS with Ca^2+^ and Mg^2+^, 0.1% Tween 20) for 5 minutes. Neurons were blocked with blocking buffer (DPBS with Ca^2+^ and Mg^2+^, 10% normal donkey serum, 0.5% Triton-X) for 1 hour at room temperature. After blocking, neurons were stained with primary antibodies at concentrations indicated in the key resources table, diluted in antibody dilution buffer (99% DPBS with Ca^2+^ and Mg^2+^, 1% normal donkey serum). Samples were incubated with primary antibodies at 4°C overnight. Neurons were next washed twice with washing buffer for 7 minutes each, and stained with secondary antibodies at concentrations indicated in the key resources table, diluted in antibody dilution buffer, for 2 hours at room temperature. Following secondary antibody incubation, samples were washed twice in washing buffer for 7 minutes each, before continuing with nuclear counterstaining as detailed further below.

To detect APOE, MAP2, and proteasomal markers (PSMA1-3,5-7, PSMB5, PSME1), the neurons were permeabilized and blocked with saponin blocking buffer (BB-S; PBS with 1% BSA, 0.05% saponin, 50 mM NH_4_Cl, 0.01% NaN_3_) for 1 hour at room temperature. Samples were incubated with primary antibodies at 4°C overnight, diluted in BB-S. Neurons were washed three times with PBS for 5 minutes each, and stained with secondary antibodies at concentrations indicated in the key resources table, diluted in BB-S, for 2 hours at room temperature. Following secondary antibody incubation, samples were washed twice in PBS for 5 minutes each, before continuing with nuclear counterstaining as detailed further below.

To detect senescence markers (MAP2, CDKN1A/p21), cells were permeabilized with permeabilization solution (TBS-T, 0.1% Triton X) at room temperature for 10 minutes. Neurons were next blocked with blocking solution (TBS-T, 5% normal donkey serum) at room temperature for 2 hours. After blocking, neurons were stained with primary antibodies at concentrations indicated in the key resources table, diluted in blocking solution. Samples were incubated with primary antibodies at 4°C overnight. Neurons were next washed twice with TBS-T for 5 minutes each, and stained with secondary antibodies at concentrations indicated in the key resources table, diluted in blocking solution, for 2 hours at room temperature. Following secondary antibody incubation, samples were washed twice in TBS-T for 5 minutes each, before continuing with nuclear counterstaining as detailed below.

For all immunocytochemistry experiments, nuclei were counterstained using Hoechst 33342 (200 ng/mL in DPBS) for 30 minutes at room temperature. Samples were washed once before being mounted onto microscope slides using Cytoseal™ 60. Images were captured using a confocal laser scanning microscope (Zeiss LSM880) fitted with a Plan-Apochromat 20X, 0.8 Air M27 objective and PMT detector, with the pinhole set to 3.46 AU for overview images, or a plan-apochromat 63X, 1.4 Oil DIC M27 objective and PMT detector, with the pinhole set to 1 AU for high-resolution images. Images were captured using standard excitation/emission settings for Hoechst/DAPI and Alexa fluorophores.

#### BULK RNA SEQUENCING AND RIBOSOME FOOTPRINTING

Mixed neuronal cultures were used at DIV49 for RNA sequencing and ribosome footprinting analysis. Four matched differentiations were used for each genotype, with each differentiation including 5*12-well plates containing 400,000 cells per well (for a total of total 24 million cells per differentiation). The ribosome sequencing approach was adapted from previously established protocols^111,112^ as follows: Neurons were treated with 100 µg/mL cycloheximide for 2 min prior to RNA extraction. Medium was replaced with cold PBS, and plates placed on ice. Cells were pipetted down from plates and transferred to falcon tubes, before centrifugation at 150 g, 4°C for 5 min. Supernatant PBS was aspirated, and 400 μL lysis buffer (20 mM Tris pH 7.5, 150 mM NaCl, 5 mM MgCl_2_, 1 mM DTT, 1% Triton X, 25 U/mL Turbo DNase, 100 µg/mL cycloheximide) added to the tube. Cell pellets were resuspended in lysis buffer by trituration, and lysed on ice for 10 min. Cells were triturated 10 times through a 26 G needle, and lysate clarified by centrifugation at 20,000 g, 4°C for 10 min. The supernatant was recovered, and flash-frozen by immersion in liquid nitrogen.

#### RNA-seq library preparation

Total RNA was extracted from cell lysates using the Quick-RNA Microprep Kit. Strand-specific RNA-seq libraries were prepared using the SMARTer Stranded Total RNA-Seq Kit v3 – Pico Input Mammalian. Final libraries were quantified with the Qubit High Sensitivity dsDNA Kit and the High Sensitivity D1000 ScreenTape on a 4200 TapeStation System and sequenced on an Illumina NovaSeq X Plus with a 10B (51 × 12 × 24 × 51) configuration by the UCSF Center for Advanced Technology (CAT) to generate 50 bp paired-end reads.

#### Nuclease footprinting and ribosome recovery

Polysome buffer with sucrose was prepared (20 mM Tris pH 7.5, 150 mM NaCl, 5 mM MgCl₂, 100 μg/mL cycloheximide, 1 mM DTT, supplemented with 1M sucrose) and stored at 4°C until use. 400 μL cell lysate was diluted to 1 mL with polysome buffer and digested with 100 U/mL RNase I at 25°C for 45 min with gentle agitation. The digested lysate was layered onto a sucrose cushion in a polycarbonate ultracentrifuge tube. Ultracentrifugation was performed at 236,000 rpm for 2 h at 4°C. Following centrifugation, the supernatant was aspirated and the ribosomal pellet was recovered and purified using the Oligo Clean & Concentrator kit according to the manufacturer’s protocol.

#### Ribosome profiling library preparation

Following purification of Ribosome-protected RNA fragments, RNA was resuspended in 10 mM Tris (pH 8.0). Approximately 5 μg of purified RNA was resolved on a 15% TBE-urea polyacrylamide gel in RNase-free 1× TBE buffer at 200 V for 65 minutes. Oligoribonucleotide size markers run in adjacent lanes were used to guide excision of gel slices corresponding to 17–34 nt ribosome footprints. Gel slices were passively eluted overnight at 4°C in 0.3 M NaCl, RNA eluate was precipitated with GlycoBlue coprecipitant and 55% isopropanol, and purified RNA was resuspended in 7 μL of 10 mM Tris (pH 8.0). Dephosphorylation was performed using T4 Polynucleotide Kinase (PNK) alongside SUPERase·In RNase inhibitor for 40 min at 37°C. RNA was subsequently purified using the Oligo Clean & Concentrator kit and eluted in nuclease-free water. Libraries were prepared using the SMARTer smRNA-Seq Kit for Illumina with a custom RT primer, incorporating a unique molecular identifier (UMI) and a custom indexing PCR reverse primer. Ribosomal RNA contaminants were depleted using the ZapR Depletion Kit. Final libraries were quantified with the Qubit High Sensitivity dsDNA Kit and the High Sensitivity D1000 ScreenTape on a 4200 TapeStation System and sequenced on an Illumina NovaSeq X Plus 10B SE100 by the UCSF Center for Advanced Technology (CAT) to generate 100 bp single-end reads.

#### RNA sequencing data processing

For total RNA read preprocessing, raw total RNA paired-end sequencing data from APOE3 and APOE4 samples were first processed by extracting UMIs located in the first 8 nts of R2 to read headers using *umi_tools107109* (v1.1.6), following the “SMARTer Stranded Total RNA-Seq Kit v3 - Pico Input Mammalian” manual guidelines. Reads were then aligned to the human genome using *STAR108110* (v2.7.1a), and then *UMICollapse*^115^ was used for deduplication. The deduplicated reads were subsequently aligned to the same GRCh38-based mRNA transcriptome (Ensembl release v96) in the RPF read preprocessing pipeline using *bowtie2* (v2.5.3).^116^ Following preprocessing, transcript-level count matrices were analyzed using *DESeq2* (v1.50.2).^117^ Features with at least 10 counts in a minimum of two samples were retained for downstream analysis. Library size normalization and differential expression analysis were performed using the *DESeq2* workflow. Differential expression was assessed by fitting negative binomial generalized linear models and comparing APOE4 to APOE3 samples. P values were adjusted using the Benjamini–Hochberg correction for multiple testing. Normalized counts, log2 fold changes, and adjusted P values were exported for downstream analyses. Features with adj.p<0.05 were considered significantly differentially expressed.

For RPF read preprocessing, raw sequencing data from APOE3 and APOE4 samples were first processed by appending unique molecular identifiers (UMIs) to read headers using a custom Python script. Raw single-end Ribo-seq reads were adapter-trimmed using *cutadapt109114* (v3.5) with options “*-m 18 -M 40 - j 4 -u 3 -a AAAAAAAAAAAAAAAAAA.”* Trimmed reads were then aligned in two steps using *bowtie2*^116^ (v2.5.3). First, reads mapping to a custom rRNA/tRNA index were discarded. The remaining reads were then aligned end-to-end to a GRCh38-based mRNA transcriptome (Ensembl release v96), in which each gene is represented by the isoform containing the longest annotated coding sequences (CDS). After alignment, *UMICollapse*^115^ was used to deduplicate reads.

For translation efficiency (TE) analysis, transcript-level Ribo-seq and matched RNA-seq count matrices were merged by transcript identifier and analyzed using *DESeq2* (v1.50.2). Features were retained for analysis if they contained at least one RPF count in a minimum of four Ribo-seq samples and at least five RNA-seq counts in a minimum of two RNA-seq samples. Raw counts were modeled using a negative binomial generalized linear model with genotype (APOE3 versus APOE4), assay type (RNA-seq versus Ribo-seq), and interactions included in the model (∼ genotype + assay + genotype:assay). Library-size normalization, dispersion estimation, and statistical testing were performed using the standard *DESeq2* workflow. Differential translation efficiency between APOE4 and APOE3 samples was assessed using the genotype-by-assay interaction term, identifying transcripts exhibiting altered ribosome occupancy relative to transcript abundance. P values were adjusted by Benjamini–Hochberg multiple testing correction. Transcripts with an adj.p < 0.05 were considered significantly differentially translated.

For codon-level ribosome occupancy analyses, Ribo-seq and RNA-seq BAM files were imported into *Ribolog* (v0.0.0.9).*112115* Ribo-seq reads were filtered to retain only in-frame ribosome-protected fragments (RPFs) 24-32 nucleotides long. Codon-level counts derived from these RPFs were corrected for ribosome stalling using the CELP (Consistent Excess of Loess Preds) method implemented in *Ribolog*. For global codon-level analysis, codon counts were extracted from the output of the *CELP_bias* function in Ribolog and summed across all detected transcripts per sample. Differential codon usage analysis was performed using *DESeq2* (v.1.42.1) to compare APOE3 and APOE4 samples. For differential stalling analysis, CELP bias coefficients were calculated separately for APOE3 and APOE4 samples and extracted for each transcript. Codons with large bias coefficients represent a consistent excess of ribosome-protected reads relative to the transcript background, indicative of translational stalling.

#### GENOMIC FEATURE ANALYSIS OF DIFFERENTIALLY EXPRESSED GENES

To identify genomic features associated with transcriptional dysregulation, we performed a gene-level feature enrichment analysis across all genes detected by RNA sequencing. Differential expression statistics were obtained from DESeq2^117^ analyses, and genes were classified as upregulated (adj.p<0.05, Log_2_(APOE4/APOE3)>0), downregulated (adj.p<0.05, Log_2_(APOE4/APOE3)<0), or background (all remaining genes). Gene annotations were retrieved from the Ensembl human gene database (GRCh38 release)114,115116,117 using the Bioconductor package *biomaRt*.116118 For each Ensembl gene identifier, genomic coordinates, gene biotype, transcript count, chromosome assignment, strand orientation, and additional annotation metadata were obtained from Ensembl. When multiple Ensembl records mapped to a single gene identifier, the longest annotated genomic interval was retained for downstream analyses. Standard autosomes and sex chromosomes were included in the analysis. Genomic sequence analyses were performed using the UCSC hg38 reference genome implemented in BSgenome.Hsapiens.UCSC.hg38.117119 Gene-body GC content was calculated from the full annotated genomic span of each gene. Gene structural features included gene length, transcript count, transcript density (transcripts per kilobase), and log-transformed gene length and transcript count. Promoter characteristics were quantified using strand-aware promoter regions defined as −2,000 bp upstream to +500 bp downstream of the transcription start site. Promoter sequences were extracted from the hg38 reference genome and analyzed for CpG content. Calculated sequence features included gene-body GC content, gene-body CpG observed/expected ratio, promoter GC content, promoter CpG density, and promoter CpG observed/expected ratio. Promoters were classified as CpG-rich when they satisfied established CpG island-like criteria of CpG observed/expected ratio > 0.6, GC content > 50%, and CpG frequency > 4%.118120 Additional genomic features included chromosome X localization, strand orientation, distance to the nearest chromosome end, gene biotype categories. Baseline expression level was estimated as Log_10_-transformed normalized mean expression across all samples.

To quantify associations between genomic features and differential expression status, separate univariate logistic regression models were fit for upregulated-versus-background and downregulated-versus-background comparisons. For continuous variables, feature values were standardized prior to modeling to facilitate comparison of effect sizes across features. Binary genomic features were analyzed without transformation. Odds ratios (ORs) and 95% confidence intervals were calculated from model coefficients. Statistical significance was assessed using Wald tests, and resulting p-values were corrected for multiple hypothesis testing within each comparison using the Benjamini–Hochberg false discovery rate (FDR).

#### TRANSCRIPT GENE SET ENRICHMENT AND OVER-REPRESENTATION ANALYSES

To identify biological pathways associated with transcriptional changes, we performed gene set enrichment analysis (GSEA) using the *fgsea*119121 package. Differential expression statistics were obtained from DESeq2 analyses. Genes were ranked using the DESeq2 Wald statistic when available. For datasets lacking the Wald statistic, genes were ranked according to the signed significance score, defined as Sign(Log_2_(APOE4/APOE3)) * -Log_10_(adj.p). Ensembl gene identifiers were mapped to HGNC gene symbols using Ensembl annotations accessed through *biomaRt*.116118 When multiple Ensembl identifiers mapped to the same gene symbol, duplicate entries were collapsed by retaining the gene with the largest absolute ranking score. Ranked gene lists were ordered from highest to lowest enrichment score. Gene sets were obtained from the Molecular Signatures Database (MSigDB)120122 using the *msigdbr* package. Analyses were performed using both the Hallmark collection and Gene Ontology Biological Process (GO BP) gene sets. Preranked GSEA was performed with *fgsea* using 10,000 permutations and gene set size thresholds of 15–500 genes. Enrichment significance was evaluated using normalized enrichment scores (NES) and Benjamini–Hochberg-adjusted false discovery rates (FDR). Positive NES values indicate enrichment among genes increased in APOE4 neurons relative to APOE3 neurons, whereas negative NES values indicate enrichment among genes decreased in APOE4 neurons. To complement rank-based enrichment analyses, over-representation analysis (ORA) was performed using the *clusterProfiler121123* package. Significantly upregulated and downregulated genes were analyzed separately, defined as genes with adj.p<0.05 and Log_2_(APOE4/APOE3)>0 or <0, respectively. The background universe consisted of all genes included in the RNA-seq dataset following identifier mapping and filtering. Hallmark and GO Biological Process gene sets were tested independently using hypergeometric enrichment analysis implemented in the *enricher* function. Benjamini–Hochberg correction was applied to account for multiple hypothesis testing, and pathways with FDR < 0.05 were considered significantly enriched. For visualization of GSEA results, the top significantly enriched pathways were ranked by their adj.p-value and normalized enrichment score. Pathways were displayed according to NES, with point size proportional to enrichment significance (-Log_10_FDR). For over-representation analysis of transcription factors associated with dysregulated transcripts, up- and downregulated genes (adj.p<0.05 and Log_2_(APOE4/APOE3)>0 or <0, respectively) were separately analyzed using Enrichr^128–130^ for ENCODE/ChEA consensus TFs identified by ChIP-X. The top five enriched transcription factors across all dysregulated transcripts, as determined by statistical significance (P<0.05) and ranked by their associated combined score, are reported in Figure S1E.

#### TRANSCRIPT FEATURE ANALYSIS OF DIFFERENTIALLY TRANSLATED GENES

To identify transcript features associated with altered translation efficiency (TE), we performed a transcript-level feature enrichment analysis across all transcripts included in the TE analysis. Differential TE statistics were obtained from *DESeq2* interaction-model analyses, and transcripts were classified as TE-upregulated (adj.p < 0.05, Log_2_(APOE4/APOE3) > 0), TE-downregulated (adj.p < 0.05, Log_2_(APOE4/APOE3) < 0), or background (all remaining transcripts). Transcript structural annotations were obtained from the Ensembl GRCh38 reference annotation (release 115)114116 using a transcript database generated with *GenomicFeatures*.125127 Transcript coding sequences (CDS), 5′ untranslated regions (5′UTRs), and 3′ untranslated regions (3′UTRs) were extracted from the reference genome using *Biostrings126128* and *Rsamtools*.127129 Transcript sequences were reconstructed in a strand-aware manner from chromosome-level FASTA files.

For each transcript, structural and sequence-derived features were calculated. Structural features included 5′UTR length, CDS length, and 3′UTR length. Sequence composition features included GC content within the 5′UTR, CDS, and 3′UTR. Codon adaptation index (CAI)128130 was calculated from CDS sequences using human codon usage frequencies, and the fraction of rare codons was quantified as the proportion of codons with relative adaptiveness values below 0.25. Upstream open reading frames (uORFs) were identified by scanning 5′UTR sequences for AUG start codons followed by an in-frame termination codon. Associations between transcript features and differential translation status were assessed using separate logistic regression models for TE-upregulated-versus-background and TE-downregulated-versus-background comparisons. Continuous features were standardized prior to modeling to facilitate comparison of effect sizes across features. For CDS length and CDS GC content, univariate logistic regression models were fit. All other transcript features were analyzed using logistic regression models adjusted for CDS length and CDS GC content. Odds ratios (ORs), 95% confidence intervals, and Wald test P values were calculated from model coefficients. P values from each comparison were separately adjusted using the Benjamini– Hochberg false discovery rate (FDR) multiple test correction.

#### FLOW CYTOMETRY TO MEASURE PUROMYCIN INCORPORATION AND CELL CYCLE STAGE

Global protein synthesis was measured using a puromycin incorporation assay based on the SUnSET (Surface Sensing of Translation) method.129131 DIV49 iPSC-derived neuronal cultures were either left untreated or incubated with 10 μM puromycin for 10 min or 60 min at 37°C and 5% CO2. Following puromycin exposure, cultures were washed once with PBS and maintained in puromycin-free medium for an additional 1 h to allow clearance of unincorporated puromycin. Untreated cultures served as negative controls. Neurons were dissociated using Accutase, transferred to round-bottom 96-well plates, and washed twice with cold PBS containing 0.1% BSA. Cells were fixed in 4% paraformaldehyde for 30 min at room temperature and subsequently blocked in PBS containing 1% BSA. Cells were incubated with anti-puromycin antibody (clone 12D10; 1:6 dilution) for 30 min at 4°C, washed with PBS/0.1% BSA, stained with Hoechst 33342 (200 ng/mL in PBS/0.1% BSA), washed again, and resuspended in PBS/0.1% BSA for flow cytometric analysis. Samples were acquired on an Attune NxT acoustic focusing flow cytometer equipped with an Attune Auto Sampler and BRVY lasers. Data were analyzed using FlowJo (v10.8.1). A uniform gating strategy was applied across all samples. Cellular debris was excluded using FSC-A and SSC-A parameters, singlets were identified using forward-scatter pulse characteristics, and Hoechst fluorescence was used to assess DNA content and cell-cycle distribution. Puromycin incorporation was quantified from Alexa Fluor 633 fluorescence intensity in single cells. Translation rates were assessed by comparing puromycin fluorescence intensity and the proportion of puromycin-positive cells between APOE3 and APOE4 neuronal cultures under matched labeling conditions. Puromycin-positivity thresholds were defined using untreated control samples, yielding 0% puromycin-positive cells in the negative-control population.

#### PROTEIN EXTRACTION FOR PROTEOMICS

For each APOE genotype-matched differentiation batch, approximately 1.2*10^6^ neurons were cultured across six wells of a 12-well plate and harvested at DIV49. Cells were washed once with ice-cold PBS and lysed directly in modified SP3 lysis buffer containing 50 mM Tris-HCl (pH 8.0), 50 mM NaCl, 1% SDS, 1% NP-40, 1% Tween-20, 1% glycerol, 1% sodium deoxycholate, 5 mM EDTA, 5 mM dithiothreitol (DTT), 0.5 KU/mL benzonase, cOmplete protease inhibitor, and PhosSTOP phosphatase inhibitor. Lysates were incubated on ice for 30 min, collected by scraping, and transferred to LoBind microcentrifuge tubes. Samples were further lysed at 65°C with continuous shaking (1200 rpm) for 30 min. Proteins were alkylated with iodoacetamide (10 mM final concentration) for 30 min in the dark. Samples were then clarified by centrifugation (16,000 g, 10 min, 4°C), and supernatants were transferred to fresh tubes. Proteins were precipitated by addition of ice-cold acetone to a final concentration of 25% (v/v) and incubation at −80°C for 1 h. Protein precipitates were recovered by centrifugation (2,000 g, 15 min, 4°C), washed twice with ice-cold acetone, air-dried for 30 min, and stored at −80°C until further processing.

Following protein extraction, protein pellets were resuspended in 40 µl of trifluoroacetic acid at approximately three times pellet volume and incubated at room temperature for 5 min. The acid was neutralized with 10x acid volume of 2 M Tris before heating at 95 °C for 5 min. Samples were diluted 1:2 with water, allowed to cool and quantified using the Protein 660 assay. Proteins were digested by adding trypsin and Lys-C at a 1:100 enzyme-to-protein ratio (each) and incubated on a thermomixer at 37 °C with 750 rpm shaking overnight.

Digested peptides (∼60 µg per sample) were acidified with 40 µl of 99% TFA (∼4% final concentration) to quench digestion before desalting on NEST Group 96-well C18 plates (20 mg PROTO 300 C18). Each well was wetted with 400 µl of wetting buffer (80% acetonitrile (ACN), 0.1% TFA), spun for 1 min at 2,000 rpm, then washed three times with 400 µl of wash buffer (0.1% TFA), spinning for 1 min at 2,400 rpm between washes. Acidified peptides were loaded onto the plate and spun through at 2,400 rpm, then washed three further times with wash buffer. Peptides were eluted into a fresh PCR plate by two sequential additions of 55 µl of elution buffer (50% ACN, 0.25% formic acid (FA)), spinning for 1 min at 2,400 rpm after each addition. Eluates were frozen at −80 °C and dried in a SpeedVac. Dried peptides were resuspended in 0.1% FA at an approximate concentration of 0.5 ug/ul and 500 nl was injected onto the LCMS platform.

In an effort to increase the proteomic depth of the spectral library, a pooled sample was fractionated using a silica C18, UltraMicroSpin Column from the NEST Group (catalog number SUM SS18V). The column was activated by addition of 100 ul of 100% ACN followed by 1 min centrifugation at 300 g before two equilibration cycles that included addition of 200 ul 0.1% TFA followed by 1 min centrifugation at 300 g. Approximately 30 ug of acidified digested peptides were added to each column, followed by two washes with 200 ul of 0.1% TFA with 1 min centrifugation at 300 g in between. Peptides were then eluted by addition of 100ul of 0.1% triethylamine (TEA) in water with increasing concentrations of ACN: 5%, 7.5%, 10%, 12.5%, 15%, 17.5%, 20%, and 50%, with the first two fractions eluted into the same tube (7 final fractions). Eluted fractions were dried down in the SpeedVac vacuum concentrator before being resuspended in 0.1% FA at a concentration of ∼200 ng/ul with ∼ 500 ng loaded on to the LCMS system.

Samples were analyzed on an Orbitrap Exploris 480 mass spectrometry system (Thermo Fisher Scientific) equipped with a Thermo EASY-nLC ultra-high-pressure liquid chromatography system (Thermo Fisher Scientific) interfaced via a Nanospray Flex source. Mobile phase A was composed of 0.1% FA and mobile phase B was composed of 0.1% FA in 80% ACN. Peptides were separated using a 15-cm long Bruker PepSep column with a 150-µm inner diameter pre-packed with 1.5 um Reprosil C18 particles (Bruker, 1893474). Peptides were separated at a flow rate of 300 nl min⁻¹ using a gradient of 4% to 30% mobile phase B over 60 min, followed by an increase to 45% B over 10 min, ending with a wash at 95% B for 6 min, for a total method duration of 80 min.

Quantitative spectra were acquired in a data-independent manner (DIA). The ion transfer tube was held at 275 °C with a spray voltages of 2000 V. Full scans (MS1) were taken from 350–1,100 *m/z* in the Orbitrap at 60,000 resolving power with a normalized auto gain control (AGC) target of 300%, a radio frequency (RF) setting of 40% and the maximum injection time set to ‘Auto’. DIA isolation windows covered 350–1,100 *m/z* space with 20-*m/z*-wide windows and 2-*m/z* overlap across 37 scan events, with window placement optimization enabled. Peptide ions were fragmented using a normalized HCD energy of 30%, with the AGC target set to ‘standard’, the maximum injection time set to ‘Auto’. MS2 scan range mode was set to ‘auto’ with a resolution of 15,000.

A spectral library was built by acquiring data-dependent spectra from select set of individual samples as well as tip-based fractions described above. Full scans were performed at 60,000 resolution from 350– 1,100 *m/z*, with a normalized AGC target of 300%, an RF setting of 40% and the maximum injection time set to ‘Auto’. Advanced peak determination was enabled. Charge states of 2–6 were selected for fragmentation using a normalized HCD energy of 30%, with a 1.6-*m/z* isolation window, an AGC target of 200%, and a maximum injection time of 22 ms at a resolution of 15,000 resolution. Dynamic exclusion of 45s was used with a mass error of 10 ppm, and cycle defined by 20 MS2 scans. DDA files were used to build a spectral library in Spectronaut using default settings searched against the full reviewed human proteome with isoforms downloaded from Uniprot(downloaded May 23, 2022) supplemented by an in-house generated human library from previous experiments. DIA files were searched against this spectral library in Spectronaut using default settings with the exception of cross-run normalization and imputation which were both disengaged. The MS Stats Format output report from Spectronaut was exported. After filtering out low intensity features, peptide features were summarized to protein-level abundances using the MSstats function dataProcess without imputation and with global standard normalization using the five housekeeping proteins described below.

#### PULSED STABLE ISOTOPE LABELING WITH AMINO ACIDS IN CELL CULTURE

Protein turnover was assessed using pulsed stable isotope labeling with amino acids in cell culture (pSILAC).^136–138^ Neuronal cultures were maintained under standard differentiation conditions until DIV35, at which point culture medium was supplemented with heavy ^13^C_6_ L-lysine to achieve a 15-fold molar excess relative to endogenous lysine present in the culture medium. Labeling was initiated by complete medium replacement at DIV35, followed by routine half-medium changes twice weekly until sample collection at DIV49. Neurons were harvested at DIV49 by washing once with ice-cold PBS before lysis in modified SP3 lysis buffer (50 mM Tris-HCl (pH 8.0), 50 mM NaCl, 1% SDS, 1% NP-40, 1% Tween-20, 1% glycerol, 1% sodium deoxycholate, 5 mM EDTA, 5 mM DTT, 0.5 KU/mL benzonase, cOmplete protease inhibitor, PhosSTOP phosphatase inhibitor). Lysates were heated at 95°C for 5 min followed by incubation at 60°C for 30 min and sonicated briefly (5 s, 20% amplitude). Protein concentrations were determined by Bradford assay. Proteins were alkylated by addition of iodoacetamide (10 mM final concentration) and incubated for 30 min, followed by addition of 10 mM DTT for 30 min at room temperature. Protein cleanup and digestion were performed on a KingFisher Flex automated purification platform using MagReSyn Amine magnetic beads and a protein aggregation capture workflow. For each sample, 100 µg protein was adjusted to a final volume of 100 µL and transferred to a 96-well plate. MagReSyn Amine beads (20 µL) were added, followed by 280 µL acetonitrile to induce protein aggregation onto the beads. Automated washing was performed using three sequential washes in 150 µL of 95% acetonitrile followed by two washes in 150 µL of 70% ethanol. Following washes, beads were transferred into 150 µL of 50 mM ammonium bicarbonate containing Lys-C protease at a 1:50 enzyme-to-protein ratio. Samples were incubated overnight at 37°C with agitation (800 rpm) to achieve proteolytic digestion. Following digestion, peptide-containing supernatants were separated from the magnetic beads and the beads were washed once with 100 µL HPLC-grade water. Washes were combined with the corresponding peptide eluates and acidified with formic acid. Samples were subsequently filtered, dried by vacuum centrifugation, and resuspended in 0.1% formic acid to a final concentration of 500 ng/µL. Peptide concentrations were determined by absorbance at 280 nm, and 1 µL of peptide solution was injected for LC-MS/MS analysis.

Samples were analyzed on an Orbitrap Exploris 480 mass spectrometry system equipped with an Easy nLC 1200 ultra-high pressure liquid chromatography system interfaced via a Nanospray Flex nanoelectrospray source. For all analyses, samples were injected on a C18 reverse phase column operated at 0.600 µL/min. Mobile phase A consisted of 0.1% formic acid (FA), and mobile phase B consisted of 0.1% FA / 80% acetonitrile (ACN). Peptides were separated using a 120 min method beginning at 3% solvent B, followed by a linear gradient to 30% B over 104 min and then to 40% B from 104 to 112 min. The column was subsequently washed at 95% B from 112 to 120 min before the run ended. The mass spectrometer was operated in positive ion mode using data-dependent acquisition. The method used a 120 min acquisition with a 2 s cycle time. Full MS1 scans were acquired in the Orbitrap over an m/z range of 350–1250 at 120,000 resolution, with an AGC target of 300% and automatic maximum injection time. MS/MS spectra were acquired for precursors with charge states 2–6, while undetermined charge states were excluded. Precursors were isolated with a 1.6 m/z isolation window and fragmented by higher-energy collisional dissociation using a normalized collision energy of 28%. MS2 spectra were acquired in the Orbitrap at 15,000 resolution with an AGC target of 200% and automatic maximum injection time. Dynamic exclusion was enabled after one selection for 20 s using a ± 10 ppm mass tolerance, and isotope exclusion was enabled. Raw LC-MS/MS data were processed using MaxQuant (v2.5.1.0). Spectra were searched against a human canonical UniProt protein database (LysC digestion enzyme, ≤ 2 missed cleavages). Carbamidomethylation of cysteine was specified as a fixed modification, while methionine oxidation and protein N-terminal acetylation were included as variable modifications. Peptide, protein, and site-level false discovery rates were controlled at 1%. Quantification was performed using a two-channel SILAC search strategy with an unlabeled/light channel and a Lys6-labeled channel. Match between runs was enabled to transfer identifications across LC-MS/MS runs, while requantification and dependent peptide searching were disabled.

Fractional protein replacement was estimated from H ratios according to F=H/(H/L), assuming complete heavy-label availability throughout the labeling period. Protein half-lives were then calculated from the 14-day labeling interval using first-order kinetics: Half-life (d)=(14*Ln(2)) / -Ln(1-F). Protein half-life estimates derived from pSILAC experiments were analyzed using the *limma* framework.133135 Prior to statistical analysis, proteins with missing half-life measurements in more than 50% of samples were excluded. Remaining protein half-life values were organized into a protein-by-sample matrix and analyzed without batch correction. Differential protein half-life analysis between APOE4 and APOE3 neuronal cultures was performed using linear models implemented in *limma*. A design matrix containing genotype as the sole explanatory variable was fitted to protein half-life values, applying empirical Bayes variance moderation. Differential statistics were extracted for the APOE4 versus APOE3 comparison, including estimated effect sizes and adj.p values based on Benjamini–Hochberg false discovery rate (FDR) correction for multiple testing. For visualization, volcano plots were generated using estimated APOE4-APOE3 effect sizes and - Log_10_(adj.p). Proteins with adj.p<0.05 were considered to exhibit significantly altered half-lives between APOE4 and APOE3 neurons. For mapping half-life changes of neurodegeneration-associated genes, gene symbols were retrieved from OpenTargets.134136

We used cosine similarity scores within defined protein complexes to assess coordinated protein turnover of complexes, as previously reported.132134 We analyzed proteins that contained (H:L) measurements in at least two APOE3 samples, at least two APOE4 samples, and at least four samples overall. Pairwise cosine similarity was calculated between proteins using their Log_2_(H:L) labeling profiles across samples. Cosine similarity matrices were generated for combined datasets as well as separately for APOE3 and APOE4 samples. Hierarchical clustering was performed using Ward’s D2 method, and cosine similarity heatmaps were generated for visualization of protein-complex organization. For each protein complex, mean pairwise cosine similarity values were calculated from the upper triangle of the corresponding cosine similarity matrix. Differences between APOE3 and APOE4 cosine similarity distributions were evaluated using both Student’s t test alongside the Benjamini–Hochberg false discovery rate (FDR) correction for multiple tests. Complex-level cosine similarity statistics were visualized as heatmaps and bar plots displaying mean ± SEM.

#### IDENTIFICATION OF HOUSEKEEPING GENES FOR NORMALIZING PROTEOMIC DATA

We processed our proteomic analyses to account for two special circumstances: (1) we accounted for variability in neuronal differentiations by applying a linear model for batch correction, and (2) previous studies have suggested that APOE4 iPSC-derived interneurons degenerate during the course of differentiation.^32^ Simultaneously, previous studies showing lysosomal impairments in APOE4 neurons suggest that the total protein content per cell might differ from APOE3 neuronal cultures.^31^ These factors would profoundly impact the proteomic results, where protein levels are analyzed based on their relative abundance compared to the protein injected into the instrument, rather than measuring the absolute number of proteins per cell. These issues are typically counteracted in analogous experiments such as RT-qPCR and Western blotting by comparison to housekeeping genes. The expression of housekeeping genes is however also subject to variation, and the ideal housekeeping protein can differ between cell-types. We therefore identified a set of ideal housekeeping proteins for normalization in iPSC-derived neurons based on mass-spectrometric analyses presented herein: First, we identified proteins whose expression levels did not change dependent on the APOE genotype based on abundance proteomics. Since later experiments would assess protein turnover that ideally should not affect the levels of housekeeping genes, we used pulsed stable isotope labelling by amino acids in cell culture (pSILAC) to identify proteins whose turnover did not differ by APOE genotype. From these proteins, we identified the proteins that were expressed at the highest levels permitting their consistent retrieval from proteomic analyses, and had the lowest turnover rendering them robust for experiments acutely altering cellular protein turnover rates. We profiled the behavior of these in a larger proteomic experiment including proteostasis inhibitors, finding that five of these behaved similarly to each other. This unbiased approach revealed five outstanding housekeeping genes for normalization: PPIA, GAPDH, TUBB3, and the histone proteins H1-5, and H2AX.

#### PROTEOMIC WEIGHTED GENE CORRELATION NETWORK ANALYSIS

After summarization, proteins were required to be identified in at least two-thirds (24/36) of all samples to be included in the weighted gene co-expression network analysis (WGCNA). A weighted protein co-expression networks was constructed on the full protein matrix after this filtering step (6668 protein groups) using the blockwiseModules function in the WGCNA package in R. The soft thresholding power was selected by evaluating the scale-free topology model fit (signed R^2^) and mean connectivity across a set of candidate powers using the pickSoftThreshold function. A power of β = 10 was chosen as the value at which the scale-free topology fit approached its asymptote while maintaining a fit > 0.7. Modules were detected using an unsigned topological overlap matrix (TOMType = “unsigned”), a minimum module size of 20, and a merge cut height of 0.10, with the PAM stage disabled with respect to the dendrogram (pamRespectsDendro = FALSE) and a reassignment threshold of 0. Clustering was performed in two blocks (maxBlockSize = 6000). This yielded 29 modules of co-expressed proteins, with proteins not assigned to any module placed in the grey module. Eigengenes for each module (excluding grey) were extracted and compared between APOE3 and APOE4 using Welch’s t-test to calculate significance. After Benjamini-Hochberg multiple hypothesis correction, four modules were found to be significant at a false discovery rate (FDR) of 5%. The module gene sets were tested for enrichment of Gene Ontology (GO Biological Process, Molecular Function and Cellular Component) terms. The over-representation analysis (ORA) was performed using the enricher function from R package clusterProfiler (v4.16.1).^127^ The gene ontology terms and annotations were obtained from the R annotation package org.Hs.eg.db (v3.22.0). In an effort to select non-redundant GO terms, we first constructed a term tree based on distances (1-Jaccard Similarity Coefficients of shared genes in GO) between the significant terms using the R function hclust. The term tree was cut at a specific level (R function cutree, h = 0.99) to identify clusters of redundant gene sets. For results with multiple significant terms belonging to the same cluster, we selected the most significant term (ranked by adjusted p value).

#### PROTEIN FEATURE ANALYSIS OF DIFFERENTIALLY EXPRESSED PROTEINS

To identify protein features associated with differential protein abundance, we performed a protein-level feature enrichment analysis across all proteins quantified by mass spectrometry. Differential protein expression statistics were obtained from housekeeping-normalized proteomic analyses, and proteins were classified as upregulated (adj.p < 0.05, Log_2_(APOE4/APOE3) > 0), downregulated (adj.p < 0.05, Log_2_(APOE4/APOE3) < 0), or background (all remaining proteins). Protein sequence information was retrieved from UniProt135137 using the UniProt REST API, using the primary accession for downstream analyses. Amino acid sequences were used to calculate protein length, molecular weight, theoretical isoelectric point (pI), grand average of hydropathy (GRAVY),136138 aliphatic index,137139 instability index,138140 and amino acid composition metrics. Theoretical isoelectric points were calculated using the EMBOSS pKa scale implemented in the *Peptides139141* R package. Protein half-life estimates derived from pSILAC experiments were incorporated where available. Subcellular localization annotations were similarly retrieved from UniProt. Baseline protein abundance was estimated as the Log_10_-transformed median raw protein abundance across all samples. To quantify associations between protein features and differential protein abundance, separate univariate logistic regression models were fit for up-or downregulated proteins versus background comparisons. Continuous features were standardized prior to modeling to facilitate comparison of effect sizes across features. Odds ratios (ORs) and 95% confidence intervals were calculated from model coefficients. Statistical significance was assessed using Wald tests, applying the Benjamini–Hochberg false discovery rate (FDR) correction for multiple testing separately for each comparison. Feature enrichment results were visualized as forest plots displaying Log_2_-transformed odds ratios and corresponding confidence intervals.

#### PROTEOMIC MEASUREMENT OF NEURONAL PROTEIN CLEARANCE

Protein clearance was measured by abundance proteomics following inhibition of protein clearance pathways. Neuronal cultures were maintained under standard differentiation conditions until DIV49, at which point culture medium was supplemented with either the lysosomal inhibitor bafilomycin A_1_ (BafA1, 200 nM), the proteasomal inhibitor MG-132 (10 μM), or the combination of both BafA_1_ and MG-132. Neurons were cultured for another 3 hours at 37°C, 5% CO_2_. Cells were washed once with ice-cold PBS and lysed directly in modified SP3 lysis buffer. Lysates were incubated on ice for 30 min, collected by scraping, and transferred to LoBind microcentrifuge tubes. Samples were further lysed at 65°C with continuous shaking (1200 rpm) for 30 min. Samples were sonicated for 20s on ice, and protein content estimated by a Bradford assay. Proteins were alkylated with 10 mM iodoacetamide, incubated in the dark for 45 minutes, and reduced by adding 5 mM dithiothreitol before being frozen at −80°C. Subsequent protein quantification, SP3 detergent removal using the KingFisher Flex system, and peptide digestion was performed in analogy to the abundance proteomics measurements detailed in the section “Protein extraction for proteomics.”

Samples were analyzed on an Orbitrap Exploris 480 mass spectrometry system equipped with an Easy nLC 1200 ultra-high pressure liquid chromatography system interfaced via a Nanospray Flex nanoelectrospray source. Samples were injected onto a PepSep C18 reverse-phase column (150 µm inner diameter * 15 cm, 1.5 µm particle size) operated at 0.500 µL/min. Mobile phase A consisted of 0.1% formic acid (FA), and mobile phase B consisted of 0.1% FA / 80% acetonitrile (ACN). Peptides were separated using an 80 min method beginning at 4% solvent B, followed by a linear gradient to 16% B over 40 min, to 28% B from 40 to 66 min, and to 40% B from 66 to 70 min. The column was then washed at 95% B from 70 to 80 min, with the flow rate increased to 0.600 µL/min during the high-organic wash. The mass spectrometer was operated in positive ion mode using data-independent acquisition (DIA). The nanospray voltage was set to 2.0 kV, and the ion transfer tube temperature was 275 °C. Full MS scans were acquired in the Orbitrap over a range of 350–1250 m/z at 60,000 resolution, with a normalized AGC target of 300%, an RF lens setting of 40%, and automatic maximum injection time. DIA MS/MS scans were acquired over a precursor mass range of m/z 350–950 using custom isolation windows of 15 m/z with 0.5 m/z overlap and window placement optimization enabled. The DIA method used a 3 s cycle time with 39 scan events. Precursors were fragmented by higher-energy collisional dissociation using a normalized collision energy of 28%. DIA MS/MS spectra were acquired in the Orbitrap at 15,000 resolution with standard AGC target, automatic maximum injection time, and centroid data collection Raw DIA LC-MS/MS files were analyzed in Spectronaut (v19.7.250203.62635) using a peptide-centric workflow and BGS factory settings. Spectra were searched against a sample-specific spectral library searched with a reviewed human UniProt database with isoforms. Precursor identifications were filtered at a Q value cutoff of 0.01, and protein identifications were filtered using an experiment-level Q value cutoff of 0.01. Protein inference was performed using the IDPicker workflow. Proteins were grouped by protein group ID. Cross-run normalization was disabled in Spectronaut. Quantitative analysis was performed on the exported Spectronaut report in the R statistical programming language (v4.1.3). Initial quality control analyses were completed with the R package *artMS* (v1.12.1). Contaminants were filtered, data were log2-transformed, and globalStandards normalization was applied using the aforementioned housekeeping proteins for normalization.

All statistical analysis was performed in the R statistical programming language (v4.5.3). Briefly, housekeeping protein-normalized protein intensities were imported into R using the fread function from the data.table package (v1.18.2.1). Protein intensities were modeled as a function of genotype, treatment and batch, with a genotype:batch interaction term to account for genotype-dependent treatment responses using base R lm function. The full model for each protein is described by LogIntensities∼genotype+treatment+batch+genotype:treatment. We used the *emmeans* package (v2.0.2) to extract pairwise contrast coefficients and associated p-values from the models for within-genotype treatment vs control comparisons. P-values were corrected for multiple testing using the Benjamini-Hochberg method, and proteins with Log_2_(APOE4/APOE3) >= 0.58 and adj.p < 0.05 were considered differentially expressed among treatment groups. Proteins demonstrating a genotype-specific treatment response (i.e an interaction effect) were identified at an adjusted p-value < 0.1. The *enricher* function from the clusterprofiler package (v4.18.4)121123 was used to identify enriched gene ontology (GO) terms among the differential sets at an adjusted p-value threshold < 0.05. We used the *removebatchEffects* function from limma (v3.66.0)133135 to correct protein intensities for known batch effects, and performed principal component analysis (PCA) on the batch-corrected Log_2_ values using the *prcomp* function to identify the strongest drivers of variance in our data. Since principal component 2 (PC2) was inversely correlated with PC1, its scores were multiplied with −1 to maintain directionality and facilitate cross-comparison across principal components. Visualizations were created using either *ggplot2* (v4.0.2)140142 or *ComplexHeatmap* (v2.26.1).141143 All analysis was performed using custom R scripts and wrapper functions available at https://github.com/kroganlab/bp_utils/blob/master/LinearModels.R.

#### INTEGRATION OF TRANSCRIPTOMIC, RIBO-SEQ, AND PROTEOMIC DATASETS

To identify molecular changes across transcriptional, translational, and proteomic layers, differential expression datasets from RNA-seq, Ribo-seq, and proteomics were integrated at the gene-symbol level. For each dataset, Log_2_(APOE4/APOE3) values and corresponding adj.p significance statistics were extracted from the primary differential analyses. When multiple entries mapped to the same gene symbol, the entry with the strongest statistical support was retained. A gene-level integration matrix was constructed containing RNA-seq, Ribo-seq, and proteomic effect sizes. Statistical significance was defined using adj.p values. Genes were retained for downstream integration if measurements were available for all three molecular layers (RNA, Ribo-seq, and protein abundance) and at least one of these layers exhibited significant differential regulation (adj.p < 0.05). Genes were classified according to their direction of regulation within the RNA-seq, Ribo-seq, and proteomic datasets, generating discrete multi-omics regulatory patterns. Cluster populations were visualized using Euler diagrams generated with the *eulerr*142144 R package. Euler diagrams were fitted directly from gene-set membership and displayed both set sizes and overlap counts, allowing visualization of concordant and discordant regulation between RNA and protein abundance changes. Genes sharing regulatory patterns were assigned to the same cluster. Within each cluster, genes were ranked according to the summed standardized effect sizes across the three molecular layers. Integrated datasets were visualized using ComplexHeatmap.141143 Heatmap values corresponded to Log_2_(APOE4/APOE3) values, and genes were grouped according to multi-omics regulatory clusters.

#### PROTEASOME ACTIVITY ASSAY

Proteasome activity was measured using a fluorogenic substrate assay adapted from an established native proteasome activity protocol.143145 Briefly, 400,000 neurons cultured in a single well of a 6-well plate were maintained until DIV49, washed with ice-cold PBS, collected into LoBind tubes, and pelleted by centrifugation (500 × g, 5 min, 4°C). Cell pellets were lysed in OK lysis buffer (50 mM Tris-HCl, pH 7.5; 2 mM DTT; 5 mM MgCl2; 10% glycerol; 2 mM ATP; 0.05% digitonin) on ice for 20 min with intermittent mixing. Lysates were clarified by centrifugation (21,300 × g, 20 min, 4°C), and protein concentrations were determined using a BCA assay. Proteasome abundance was estimated by immunoblotting for the integral, catalytic β5 proteasome subunit (PSMB5). Equal amounts of protein (7.5 μg) were resolved on Criterion TGX precast gels and transferred to PVDF membranes using a Trans-Blot Turbo system. Membranes were blocked in TBS-T containing 5% (w/v) milk and incubated with anti-PSMB5 primary antibody (1:500) overnight at 4°C followed by HRP-conjugated anti-rabbit secondary antibody (1:1000). Immunoreactive bands were visualized using SuperSignal™ West Pico substrate and imaged on a ChemiDoc XRS+ system. PSMB5 band intensities were quantified using Fiji/ImageJ. For proteasome activity measurements, lysate equivalent to 1.5 μg total protein was combined with reaction buffer (50 mM Tris-HCl, pH 7.5; 10 mM MgCl2; 1 mM ATP; 1 mM DTT; 48 μM Suc-LLVY-AMC) in a final volume of 100 μL in flat-bottom 96-well plates. Where indicated, 10 μM MG-132 was included as a negative control. Fluorescence was measured using a SpectraMax ID8 plate reader (380 nm excitation, 460 nm emission) at 1-min intervals for 90 min. Fluorescence signals were normalized to the mean baseline signal recorded during the first 5 min and subsequently normalized to PSMB5 abundance determined by densitometry. Proteasome activity was quantified as the slope of fluorescence increase (ΔAU/min) determined by linear regression. Differences between APOE3 and APOE4 lysates were evaluated using restricted maximum likelihood (REML) mixed-effects models with fixed effects for time, genotype, and the time:genotype interaction, implemented in GraphPad Prism (v11.0.0).

#### PROXIMITY LIGATION ASSAY TO DETECT PROTEASOMAL APOE

To measure the spatial proximity between APOE and proteasomes, neurons were cultured on glass coverslips until DIV49 and fixed with 4% PFA for 20 min at room temperature. Following two PBS washes, cells were permeabilized and blocked for 1 h at room temperature in BB-S (PBS containing 1% BSA, 0.05% saponin, 50 mM NH_4_Cl, and 0.01% NaN_3_). Samples were incubated overnight at 4°C with mouse anti-APOE and rabbit anti-PSMB5 primary antibodies for proximity ligation, together with chicken anti-MAP2 for neuronal counterstaining. The proximity ligation assay was performed using the Duolink® In Situ Red Starter Kit Mouse/Rabbit according to the manufacturer’s instructions with minor modifications for gentle permeabilization.^31^ Following primary antibody incubation, samples were washed twice in a modified Wash Buffer A (10 mM Tris (pH 7.4), 150 mM NaCl, 0.05% saponin). Samples were next incubated with anti-chicken Alexa-488 secondary antibodies for MAP2 staining, and anti-rabbit PLUS and anti-mouse MINUS PLA probes diluted in BB-S for 1 h at 37°C. After probe binding, samples were subjected to ligation for 30 min at 37°C followed by rolling-circle amplification for 100 min at 37°C with Detection Reagents Red. Slides were subsequently washed in Wash Buffer B and mounted using Duolink® In Situ Mounting Medium containing DAPI for confocal microscopy. Images were captured using a confocal laser scanning microscope (Zeiss LSM880) fitted with a plan-apochromat 63X/1.4 Oil DIC M27 objective and PMT detector. Nuclei were imaged by 405 nm excitation/459 emission, MAP2 by 488 nm excitation/549 emission, and PLA by 594 nm excitation/645 emission, at a pixel resolution of 70 nm.

#### LONGITUDINAL PROTEOMICS

For each APOE genotype-matched differentiation batch throughout the differentiation protocol, cell pellets (2-10*10^5^ cells) were collected in LoBind tubes by Accutase dissociation and centrifugation at 400 g, 4 min (DIV0), washing and centrifugation 200 g, 2 min (DIV7-DIV21), or flushing off wells and centrifugation for 500 g, 5 min (DIV28-DIV49). After collection by centrifugation, cells were washed once with ice-cold PBS and lysed in modified SP3 lysis buffer containing 50 mM Tris-HCl (pH 8.0), 50 mM NaCl, 1% SDS, 1% NP-40, 1% Tween-20, 1% glycerol, 1% sodium deoxycholate, 5 mM EDTA, 5 mM dithiothreitol (DTT), 0.5 KU/mL benzonase, cOmplete protease inhibitor, and PhosSTOP phosphatase inhibitor. Cells were lysed by heating at 95 C for 5 mins in lysis buffer, before performing two freeze-thaw cycles and probe sonication at 20% amplitude for 15 seconds. Protein concentration was determined by Bradford assay before proteins were alkylated with iodoacetamide (10 mM final concentration) for 30 min in the dark. Alkylation was quenched by addition of DTT to final concentration of 20 mM and incubation for 30 mins at room temperature.

Following protein extraction, cleanup and digestion were performed on a KingFisher Flex automated purification system using a protein aggregation capture (PAC/SP3-like) workflow with MagReSyn Amine magnetic beads. For each sample, 100 µL of protein lysate containing 50 µg total protein was transferred to a 96-well plate and combined with 20 µL MagReSyn Amine beads. Protein aggregation onto the beads was induced by addition of 280 µL acetonitrile (final concentration 70% v/v), followed by automated mixing and bead capture using the KingFisher Flex instrument. Bound proteins were washed three times in 150 µL of 95% acetonitrile and twice in 150 µL of 70% ethanol to remove detergents, salts, and other contaminants. Following washing, beads were transferred into 150 µL of 50 mM ammonium bicarbonate (ABC; pH 7.8) containing Lys-C and trypsin (1:100 enzyme-to-protein ratio). Samples were incubated overnight at 37°C with continuous agitation (800 rpm) to achieve proteolytic digestion. Following digestion, peptide-containing supernatants were separated from the magnetic beads and the beads were washed once with HPLC-grade water. Washes were combined with the corresponding peptide eluates and acidified with formic acid. Samples were subsequently filtered through 0.45 µm membranes, dried by vacuum centrifugation, and stored at −80°C until LC-MS/MS analysis. Peptide samples were resuspended in 0.1% formic acid to a final concentration of 500 ng/µL, with LCMS injections of 1 uL.

Samples were analyzed on an Orbitrap Exploris 480 mass spectrometry system (Thermo Fisher Scientific) equipped with a Vanquish Neo ultra-high-pressure liquid chromatography system (Thermo Fisher Scientific) interfaced via a Nanospray Flex source. Mobile phase A was composed of 0.1% FA and mobile phase B was composed of 0.1% FA in 80% ACN. Peptides were separated using a 15-cm long Bruker PepSep column with a 150-µm inner diameter pre-packed with 1.5 µm Reprosil C18 particles (Bruker, 1893474). Peptides were separated at a flow rate of 600 nl min⁻¹ using a gradient of 3% to 28% mobile phase B over 67 min, followed by an increase to 40% B over 5 min, ending with a wash at 95% B for 8 min, for a total method duration of 80 min. Sample loading was pressure-controlled with a maximum pressure of 300 bar.

Spectra were acquired in a data-independent manner (DIA). The ion transfer tube was held at 275 °C, a spray voltage of 2000 V. Full scans (MS1) were taken from 350–1,250 m/z in the Orbitrap at 60,000 resolution with a normalized AGC target of 300%, an RF setting of 40% and the maximum injection time set to ‘Auto’. DIA isolation windows covered 350–1,250 m/z space with 15-m/z-wide windows and 0.5-m/z overlap across 59 scan events, with window placement optimization enabled. Peptide ions were fragmented using a normalized HCD energy of 30%, with the AGC target set to ‘standard’, the maximum injection time set to ‘Auto’. MS2 scan range mode was set to ‘auto’ with a resolution of 15,000. DIA files were searched against the same spectral library described above in Spectronaut using default settings with the exception of cross-run normalization and imputation, which were both disengaged.

Protein group intensities from longitudinal iPSC-derived neuronal differentiations were post-processed to generate an APOE-agnostic temporal expression matrix. Protein group intensities were normalized within each sample to the housekeeping proteins GAPDH and PPIA, which were found to be stable throughout differentiation. Protein identifiers were mapped to gene symbols and collapsed to gene-level values by taking the median normalized intensity across technical replicates within each subject, differentiation day, and genotype. Differentiation days were assigned to broad developmental bins corresponding to iPSC (DIV0), neural progenitor cell (NPC; DIV7–DIV21), and neuronal stages (DIV28–DIV49). Genes detected in fewer than two developmental bins were excluded from downstream analysis. To identify stage-associated proteins, gene-level median abundances were summarized within iPSC, NPC, and neuronal stages and tested for stage association using an APOE-agnostic linear model. Genes exhibiting significant stage dependence following multiple-testing correction (BH-adjusted p < 0.05) were assigned to the developmental stage in which they showed the greatest relative enrichment. Stage specificity was quantified as the difference between the median abundance in the assigned stage and the highest median abundance observed across the remaining stages. Stage-enriched proteins were further filtered for minimum stage-specific detection, requiring detection in at least one iPSC sample, two NPC samples, or three neuronal samples, depending on the assigned stage. For heatmap generation, technical replicates were collapsed within each differentiation day, preserving one value per gene per DIV. The resulting matrix was ordered by developmental stage and stage-specificity score, then row-centered by subtracting the median abundance of each gene across differentiation time points. Rows were clustered within stage-specific blocks using complete-linkage hierarchical clustering following median-based imputation of missing values. The final heatmap displayed normalized, row-centered protein abundances across differentiation. To visualize temporal patterns of APOE4-associated proteomic dysregulation, proteins assigned to longitudinal dysregulation clusters were analyzed across differentiation time points. Log_2_(APOE4/APOE3) values were summarized by cluster and differentiation day (DIV), and mean cluster trajectories were calculated as the average Log_2_(APOE4/APOE3) value of all proteins within each cluster at each time point. Smoothed cluster trajectories were generated using LOESS regression. Cluster-associated gene-set enrichment analysis was performed separately for each cluster using STRING,^39^ and non-redundant terms showing significant cluster enrichment based on the FDR presented along the heatmap.

#### SEAHORSE MEASUREMENT OF NEURONAL RESPIRATION

APOE3 and APOE4 iPSCs were cultured until DIV21, at which point 40,000 neuronal progenitor cells were seeded into wells of poly-L-lysine/laminin-coated Seahorse XFe96 FluxPak 96-well plates. Cells were maintained in N2/B27 medium and fed by half-feeds twice per week. The Seahorse assay was carried out at DIV49 according to the manufacturer’s instructions. The sensor cartridge was calibrated by probe immersion in Seahorse XF calibrant in a CO_2_-free incubator overnight. Seahorse medium was prepared by supplementing Seahorse XF DMEM with 1 mM pyruvate, 2 mM glutamine, 10 mM glucose, 0.5X B27, 0.5X N2, 10 ng/mL BDNF, and 10 ng/mL GDNF (optionally removing pyruvate and glutamine for restriction analysis, where indicated). Oligomycin, FCCP, and rotenone/antimycin A (AA) resuspended in assay medium and loaded into sensor cartridge injection ports. Neuronal culture medium was exchanged for Seahorse medium and placed in a non-CO_2_ incubator 1 hour prior to the assay. The sensor plate was calibrated in the Seahorse XFe96 Extracellular Flux Analyzer for 30 minutes prior to the assay. The sensor plate was mounted onto the XFe96 FluxPak 96-well plate and samples loaded into the Seahorse XFe96 Extracellular Flux Analyzer controlled by the Agilent Wave (v2.6.4.24) software. Oxygen consumption rate (OCR) and extracellular acidification rate (ECAR) was measured at baseline, and again after sequential addition of 1.5 µM oligomycin, 1 µM FCCP, and 0.5 µM rotenone/AA. Three measurements were taken at baseline and following each treatment at intervals of 400 s. Following the assay, cells were washed once with PBS before estimating cell content by the Bradford assay. Respiration was calculated relative to the protein content and the average basal OCR of each well.

## QUANTIFICATION AND STATISTICAL ANALYSIS

Data was processed using a combination of Microsoft Excel and RStudio/R version 4.3.2.144146 R packages used are indicated in the associated methods sections. Steps taken to process and normalize data are specified in the associated methods sections, figure axes, and figure legends. Statistical tests and associated metrics are described in the figure legends of their associated graphs. All data presented was collected from at least three separate technical and biological replicates, where a biological replicate is defined as a separate differentiation batch, started at different days from distinct iPSC passage numbers.

### Fiji analysis pipelines of microscopy images

Confocal microscopy images were analyzed in a blinded manner using Fiji.^108,151^ Where applicable, we used automated image analysis scripts for consistent measurements across experimental conditions. All image processing parameters, including thresholding, morphological operations, and ROI generation, were applied identically across all experimental conditions.

To calculate the proportions of Ki67- and GABA-immunoreactive neurons by APOE genotype, 20X images of GABA/MAP2-stained neuronal cultures were captured, and a nuclear mask was generated based on Hoechst-staining by mean filtering (block radius=5) followed by thresholding using Li AutoThreshold. Nuclei were segmented by watershed, followed by EDM Binary Operations (5 iterations, open). Nuclear masks were segmented by analyze particles (>500 pixel units) and added to the ROI Manager. Nuclear GABA and Ki67 immunoreactivity analyzed using the roiManager measure function.

To calculate GABAergic neuron proportions per differentiation batch, 63X images of GABA/TuJ-stained neuronal cultures were captured, and a nuclear mask was generated based on Hoechst-staining by mean filtering (block radius=20) followed by thresholding using Li AutoThreshold. Nuclei were segmented by watershed, followed by EDM Binary Operations (10 iterations, open). The nuclear mask selection was added to the ROI Manager, and nuclear GABA immunoreactivity analyzed using the roiManager measure function. The area measurement of the nuclear masks was used for retrospective analyses of nuclear size.

To calculate colocalization of proteasomal subunits with each-other from immunofluorescent images, 63X images of stained neuronal cultures were captured. To calculate 20S assembly, neuronal regions were identified from MAP2 immunostaining and nuclei were identified using DAPI staining. For each image, MAP2 and DAPI channels were smoothed using a mean filter (10-pixel radius) and segmented using the Li automatic thresholding method. Binary masks were generated and converted to regions of interest (ROIs). To define neuronal soma regions, MAP2-positive and DAPI-positive ROIs were combined and subjected to morphological opening and dilation operations to generate a soma mask. Neurite regions were defined as MAP2-positive areas excluding the soma mask. Compartment-specific image exports were generated by masking either soma or neurite regions and splitting fluorescence channels. The % subunit association was calculated using the ImageJ *JACoP* plugin146148 to obtain Manders overlap coefficients with automatic thresholding.

To calculate neuronal PSME1 abundance and distribution, 63X images of PSME1/MAP2-stained neuronal cultures were captured. Individual fluorescence channels were smoothed using a mean filter (3-pixel radius) and segmented using the Default automatic thresholding algorithm to generate binary masks. Binary masks from all channels were combined to generate a composite cellular mask representing the total labeled cellular area. Cells were identified by particle analysis (>10,000 pixel units). A nuclear mask was generated based on Hoechst staining, expanded to cover cell bodies, and converted to a region of interest (ROI). The soma ROI was intersected with the cellular ROI to restrict analysis to cell bodies. A complementary cellular ROI outside the cell body was generated by subtracting punctate regions from the total cellular area, giving rise to the neurite mask. Mean PSME1 fluorescence was measured independently within punctate and non-punctate cellular compartments.

For object-based colocalization analysis of APOE and PSMB5, 63X images of PSMB5/APOE-stained neuronal cultures were analyzed in IamgeJ/Fiji. Multi-channel images were split into individual channels. For each fluorescence channel, a theoretical point-spread function (PSF) was generated using the microscope parameters (refractive index 1.518, numerical aperture 1.4). Channel-specific deconvolution was then performed using iterative deconvolution (termination threshold 0.005, maximum 100 iterations). Deconvolved channels were re-merged and saved for downstream analysis and figure representation. To define the cellular region of interest (ROI), the MAP2 and DAPI channels were segmented separately using Li thresholding after smoothing, converted to binary masks, and morphologically dilated. The resulting masks were combined to generate a single cell ROI, which was applied to the PSMB5 and APOE channels to restrict analysis to the cellular area. Object-based colocalization analysis146148 was performed on the deconvolved PSMB5 and APOE channels. Each channel was thresholded using the Moments algorithm to generate binary object masks. Puncta were identified using particle analysis with a size range of 0.03–1.5 µm². For each channel, the number of objects was quantified, and overlap with the reciprocal channel was determined by measuring mean intensity within each object ROI and scoring objects as overlapping when the mean reciprocal-channel intensity was greater than zero. Puncta density was calculated as the number of objects per unit cell area, and the percentage overlap was calculated as the fraction of objects showing signal in the reciprocal channel.

For quantification of APOE:PSMB5 proximity ligation signal, 63X confocal images containing APOE:PSMB5 PLA signal, MAP2 immunofluorescence, and DAPI nuclear staining were analyzed using Fiji/ImageJ. Due to elevated background PLA signal within nuclei, a nuclear mask was generated from the DAPI channel following mean filtering (radius = 20 pixels) and segmentation using the Li automatic thresholding algorithm. PLA signal within the nuclear mask was excluded from all subsequent analyses. A neuronal mask was generated from MAP2 immunofluorescence to restrict quantification to neuronal structures. APOE PLA puncta were identified using the *Find Maxima* algorithm and quantified within the non-nuclear MAP2-positive area. Neurite length was determined from skeletonized MAP2 staining, and PLA puncta counts were normalized to total neurite length. Statistical analysis and data visualization were performed using GraphPad Prism.

### Statistical Analysis

The following statistical approaches were used for hypothesis testing, unless otherwise indicated: For comparisons between two conditions (one independent variable), a two-way t-test was performed. In cases where genotype-matched experiments were performed, a paired two-way t-test was performed. For comparisons between multiple conditions across one independent variable, a one-way ANOVA was performed, followed *post-hoc* by Bonferroni’s multiple comparison test. For comparisons between multiple conditions across more than one independent variables, a two-way ANOVA was performed, followed *post-hoc* by Bonferroni’s multiple comparison test. Results were considered statistically significant when the P value was below 0.05. All bar graphs present data as the mean ± SEM, unless otherwise stated.

## REFERENCES

1. WHO. WHO Fact Sheets: Dementia. WHO Fact Sheets (31AD).

2. Zhu, X. C. et al. Rate of early onset Alzheimer’s disease: A systematic review and meta-analysis. Ann. Transl. Med. 3, (2015).

3. Reitz, C. & Mayeux, R. Use of genetic variation as biomarkers for mild cognitive impairment and progression of mild cognitive impairment to dementia. Journal of Alzheimer’s Disease vol. 19 229–251 Preprint at 10.3233/JAD-2010-1255 (2010).

4. Hunsberger, H. C., Pinky, P. D., Smith, W., Suppiramaniam, V. & Reed, M. N. The role of APOE4 in Alzheimer’s disease: strategies for future therapeutic interventions. Neuronal Signal. 3, 1–15 (2019).

5. Blumenfeld, J., Yip, O., Kim, M. J. & Huang, Y. Cell type-specific roles of APOE4 in Alzheimer disease. Nature Reviews Neuroscience vol. 25 91–110 Preprint at 10.1038/s41583-023-00776-9 (2024).

6. Knoferle, J. et al. Apolipoprotein E4 produced in GABAergic interneurons causes learning and memory deficits in mice. Journal of Neuroscience 34, 14069–14078 (2014).

7. Wang, C. et al. Selective removal of astrocytic APOE4 strongly protects against tau-mediated neurodegeneration and decreases synaptic phagocytosis by microglia. Neuron 109, 1657–1674.e7 (2021).

8. Xiong, M. et al. Astrocytic APOE4 removal confers cerebrovascular protection despite increased cerebral amyloid angiopathy. Mol. Neurodegener. 18, (2023).

9. Harris, F. M. et al. Astroglial regulation of apolipoprotein E expression in neuronal cells: Implications for Alzheimer’s disease. Journal of Biological Chemistry 279, 3862–3868 (2004).

10. Lanfranco, M. F., Sepulveda, J., Kopetsky, G. & Rebeck, G. W. Expression and secretion of apoE isoforms in astrocytes and microglia during inflammation. Glia 69, 1478–1493 (2021).

11. Lee, H. et al. ApoE4-dependent lysosomal cholesterol accumulation impairs mitochondrial homeostasis and oxidative phosphorylation in human astrocytes. Cell Rep. 42, (2023).

12. de Leeuw, S. M. et al. APOE2, E3, and E4 differentially modulate cellular homeostasis, cholesterol metabolism, and inflammatory response in isogenic iPSC-derived astrocytes. Stem Cell Reports 17, 110–126 (2022).

13. Boschert, U., Merlo-Pich, E., Higgins, G., Roses, A. D. & Catsicas, S. Apolipoprotein E Expression by Neurons Surviving Excitotoxic Stress. http://www.idealibrary.com (1999).

14. Xu, Q. et al. Profile and regulation of apolipoprotein E (ApoE) expression in the CNS in mice with targeting of green fluorescent protein gene to the ApoE locus. Journal of Neuroscience 26, 4985–4994 (2006).

15. Xu, Q. et al. Intron-3 retention/splicing controls neuronal expression of apolipoprotein E in the CNS. Journal of Neuroscience 28, 1452–1459 (2008).

16. Keren-Shaul, H. et al. A Unique Microglia Type Associated with Restricting Development of Alzheimer’s Disease. Cell 169, 1276–1290.e17 (2017).

17. Rangaraju, S. et al. Quantitative proteomics of acutely-isolated mouse microglia identifies novel immune Alzheimer’s disease-related proteins. Mol. Neurodegener. 13, (2018).

18. Jiang, Q. et al. ApoE Promotes the Proteolytic Degradation of Aβ. Neuron 58, 681–693 (2008).

19. Lee, C. Y. D., Tse, W., Smith, J. D. & Landreth, G. E. Apolipoprotein E promotes β-amyloid trafficking and degradation by modulating microglial cholesterol levels. Journal of Biological Chemistry 287, 2032–2044 (2012).

20. Liu, C. C. et al. Cell-autonomous effects of APOE4 in restricting microglial response in brain homeostasis and Alzheimer’s disease. Nat. Immunol. 24, 1854–1866 (2023).

21. Bassal, R. et al. APOE4 impairs autophagy and Aβ clearance by microglial cells. Inflammation Research 74, (2025).

22. Serrano-Pozo, A. et al. Effect of APOE alleles on the glial transcriptome in normal aging and Alzheimer’s disease. *Nat*. Aging 1, 919–931 (2021).

23. Koutsodendris, N. et al. Neuronal APOE4 removal protects against tau-mediated gliosis, neurodegeneration and myelin deficits. *Nat*. Aging (2023) doi:10.1038/s43587-023-00368-3.

24. Tong, L. M. et al. Inhibitory interneuron progenitor transplantation restores normal learning and memory in ApoE4 knock-in mice without or with Aβ accumulation. Journal of Neuroscience 34, 9506– 9515 (2014).

25. Huang, Y. et al. Apolipoprotein E fragments present in Alzheimer’s disease brains induce neurofibrillary tangle-like intracellular inclusions in neurons. Proc. Natl. Acad. Sci. U. S. A. 98, 8838–8843 (2001).

26. Chang, S. et al. Lipid- and receptor-binding regions of apolipoprotein E4 fragments act in concert to cause mitochondrial dysfunction and neurotoxicity. Proc. Natl. Acad. Sci. U. S. A. 102, 18694–18699 (2005).

27. Nakamura, T., Watanabe, A., Fujino, T., Hosono, T. & Michikawa, M. Apolipoprotein E4 (1-272) fragment is associated with mitochondrial proteins and affects mitochondrial function in neuronal cells. Mol. Neurodegener. 4, 1–11 (2009).

28. Orr, A. L. et al. Neuronal Apolipoprotein E4 Expression Results in Proteome-Wide Alterations and Compromises Bioenergetic Capacity by Disrupting Mitochondrial Function. Journal of Alzheimer’s Disease 68, 991–1011 (2019).

29. Cakir, Z. et al. Quantitative Proteomic Analysis Reveals apoE4-Dependent Phosphorylation of the Actin-Regulating Protein VASP. Mol. Cell. Proteomics 22, 100541 (2023).

30. Krogsaeter, E. K. et al. Lysosomal proteomics reveals mechanisms of neuronal apoE4-associated lysosomal dysfunction. Autophagy 2023.10.02.560519v1, 1–26 (2025).

31. Wang, C. et al. Gain of toxic apolipoprotein E4 effects in human iPSC-derived neurons is ameliorated by a small-molecule structure corrector. Nat. Med. 24, 647–657 (2018).

32. Ahn, S., Kim, T.-G., Kim, K.-S. & Chung, S. Differentiation of human pluripotent stem cells into Medial Ganglionic Eminence vs. Caudal Ganglionic Eminence cells. Methods 101, 103–112 (2016).

33. Altmann, A., Tian, L., Henderson, V. W. & Greicius, M. D. Sex modifies the APOE-related risk of developing Alzheimer disease. Ann. Neurol. 75, 563–573 (2014).

34. Neu, S. C. et al. Apolipoprotein E genotype and sex risk factors for Alzheimer disease: A meta-analysis. JAMA Neurol. 74, 1178–1189 (2017).

35. Shi, X. et al. An epigenetic switch induced by Shh signalling regulates gene activation during development and medulloblastoma growth. Nat. Commun. 5, (2014).

36. Salcedo-Tacuma, D. et al. Differential Methylation Levels in CpGs of the BIN1 Gene in Individuals With Alzheimer Disease. Alzheimer Dis. Assoc. Disord. 33, 321–326 (2019).

37. Walker, R. M. et al. Identification of epigenome-wide DNA methylation differences between carriers of APOE ε4 and APOE ε2 alleles. Genome Med. 13, (2021).

38. Szklarczyk, D. et al. The STRING database in 2023: protein–protein association networks and functional enrichment analyses for any sequenced genome of interest. Nucleic Acids Res. 51, D638– D646 (2023).

39. Ouwenga, R., Lake, A. M., Aryal, S., Lagunas, T. & Dougherty, J. D. The differences in local translatome across distinct neuron types is mediated by both baseline cellular differences and posttranscriptional mechanisms. eNeuro 5, (2018).

40. Lyu, X., Yang, Q., Zhao, F. & Liu, Y. Codon usage and protein length-dependent feedback from translation elongation regulates translation initiation and elongation speed. Nucleic Acids Res. 49, 9404– 9423 (2021).

41. Ishimura, R. et al. Ribosome stalling induced by mutation of a CNS-specific tRNA causes neurodegeneration. Science (1979). 345, 455–459 (2014).

42. Rimal, S. et al. Inefficient quality control of ribosome stalling during APP synthesis generates CAT-tailed species that precipitate hallmarks of Alzheimer’s disease. Acta Neuropathol. Commun. 9, (2021).

43. Ong, S. E. et al. Stable isotope labeling by amino acids in cell culture, SILAC, as a simple and accurate approach to expression proteomics. Mol. Cell. Proteomics 1, 376–386 (2002).

44. Boisvert, F. M. et al. A quantitative spatial proteomics analysis of proteome turnover in human cells. Molecular and Cellular Proteomics 11, (2012).

45. Bellenguez, C. et al. New insights into the genetic etiology of Alzheimer’s disease and related dementias. Nat. Genet. 54, 412–436 (2022).

46. Moulard, B., Sefiani, A., Laamri, A., Malafosse, A. & Camu, W. NEUROLOGICAL SCIENCES Apolipoprotein E Genotyping in Sporadic Amyotrophic Lateral Sclerosis: Evidence for a Major Influence on the Clinical Presentation and Prognosis. Journal of the Neurological Sciences vol. 139 (1996).

47. Di Biase, E. et al. ApoE4 requires lipidation enhancement to resolve cellular lipid and protein abnormalities following NPC1 inhibition. Sci. Rep. 15, (2025).

48. Frankel, L. B., Lubas, M. & Lund, A. H. Emerging connections between RNA and autophagy. Autophagy 13, 3–23 (2017).

49. Korolchuk, V. I., Menzies, F. M. & Rubinsztein, D. C. Mechanisms of cross-talk between the ubiquitin-proteasome and autophagy-lysosome systems. FEBS Letters vol. 584 1393–1398 Preprint at 10.1016/j.febslet.2009.12.047 (2010).

50. Li, C. et al. Proteasome Inhibition Activates Autophagy-Lysosome Pathway Associated With TFEB Dephosphorylation and Nuclear Translocation. Front. Cell Dev. Biol. 7, (2019).

51. Bendiske, J. & Bahr, B. A. Lysosomal activation is a compensatory response against protein accumulation and associated synaptopathogenesis - An approach for slowing Alzheimer disease? J. Neuropathol. Exp. Neurol. 62, 451–463 (2003).

52. Nuriel, T. et al. The endosomal-lysosomal pathway is dysregulated by APOE4 expression in vivo. Front. Neurosci. 11, 1–12 (2017).

53. Morrow, J. A. et al. Apolipoprotein E4 forms a molten globule: A potential basis for its association with disease. Journal of Biological Chemistry 277, 50380–50385 (2002).

54. Wu, D. gui, Wang, Y. na, Zhou, Y., Gao, H. & Zhao, B. Inhibition of the Proteasome Regulator PA28 Aggravates Oxidized Protein Overload in the Diabetic Rat Brain. Cell. Mol. Neurobiol. 43, 2857– 2869 (2023).

55. Pickering, A. M. & Davies, K. J. A. Differential roles of proteasome and immunoproteasome regulators Pa28αβ, Pa28γ and Pa200 in the degradation of oxidized proteins. Arch. Biochem. Biophys. 523, 181–190 (2012).

56. Nunomura, A., et al. Oxidative Damage Is the Earliest Event in Alzheimer Disease. Journal of Neuropathology and Experimental Neurology vol. 60 https://academic.oup.com/jnen/article/60/8/759/2916237 (2001).

57. Samelson, A. J. et al. CRISPR screens in iPSC-derived neurons reveal principles of tau proteostasis. Cell 189, 1517–1534.e19 (2026).

58. Paradise, V. et al. Neuroproteasomes regulate endogenous tau paired helical filament formation in an APOE genotype- and age-dependent manner. Nat. Neurosci.(2026) doi:10.1038/s41593-026-02297-x.

59. Gruenthal, E. Klinisch-anatomisch vergleichende Untersuchungen ueber den Greisenbloedsinn. Zeitschrift für die gesamte Neurologie und Psychiatrie 111, 763–818 (1927).

60. Ulrich, J. Alzheimer changes in nondemented patients younger than sixty-five: Possible early stages of Alzheimer’s disease and senile dementia of Alzheimer type. Ann. Neurol. 17, 273–277 (1985).

61. Tomlinson, B. E. & Roth, M. Observations on the Brains of Non-Demented Old People. J. Neurol. Sci. 7, 331–356 (1968).

62. Braak, H. & Braak, E. Acta H’ Pathologica Neuropathological Stageing of Alzheimer-Related Changes. Acta Neuropathol vol. 82 (1991).

63. Ohm, T. Does Alzheimer’s disease start early in life? Mol. Psychiatry 2, 21–25 (1997).

64. Ohm, T. G., Moller, H., Braakt, H. & Bohl, J. CLOSE-MESHED PREVALENCE RATES OF DIFFERENT STAGES AS A TOOL TO UNCOVER THE RATE OF ALZHEIMER’S DISEASE-RELATED NEUROFIBRILLARY CHANGES. vol. 64 (1995).

65. Snowdon, D. A. et al. Linguistic Ability in Early Life and Cognitive Function and Alzheimer’s Disease in Late Life: Findings From the Nun Study. JAMA 7, 528–532 (1996).

66. Ohm, T. G., Scharnagl, H., März, W. & Bohl, J. Apolipoprotein E isoforms and the development of low and high Braak stages of Alzheimer’s disease-related lesions. Acta Neuropathol 98, 273–280 (1999).

67. Cataldo, A. M., et al. Gene Expression and Cellular Content of Cathepsin D in Alzheimer’s Disease Brain: Evidence for Early Up-Regulation of the EndosomaI-Lysosomal System. Neuron vol. 14 (1995).

68. Cataldo, A. M., Barnett, J. L., Pieroni, C. & Nixon, R. A. Increased Neuronal Endocytosis and Protease Delivery to Early Endosomes in Sporadic Alzheimer’s Disease: Neuropathologic Evidence for a Mechanism of Increased-Amyloidogenesis. (1997).

69. Selkoe, D. J. Altered protein composition of isolated human cortical neurons in alzheimer disease. Ann. Neurol. 8, 468–478 (1980).

70. Jurk, D. et al. Postmitotic neurons develop a p21-dependent senescence-like phenotype driven by a DNA damage response. Aging Cell 11, 996–1004 (2012).

71. Herdy, J. R., Mertens, J. & Gage, F. H. Neuronal senescence may drive brain aging. Science (1979). 384, 1404–1406 (2024).

72. Kang, C. et al. The DNA damage response induces inflammation and senescence by inhibiting autophagy of GATA4. Science (1979). 349, (2015).

73. Musi, N. et al. Tau protein aggregation is associated with cellular senescence in the brain. Aging Cell 17, (2018).

74. Dehkordi, S. K. et al. Profiling senescent cells in human brains reveals neurons with CDKN2D/p19 and tau neuropathology. *Nat*. Aging 1, 1107–1116 (2021).

75. Golde, T. E. & Miller, V. M. Proteinopathy-induced neuronal senescence: a hypothesis for brain failure in Alzheimer’s and other neurodegenerative diseases. Alzheimers Res. Ther. 1, 5 (2009).

76. Herdy, J. R. et al. Increased post-mitotic senescence in aged human neurons is a pathological feature of Alzheimer’s disease. Cell Stem Cell 29, 1637–1652.e6 (2022).

77. Sutherland, T. C. et al. Age-dependent decline in neuron growth potential and mitochondria functions in cortical neurons. Cells 10, (2021).

78. Di Fraia, D. et al. Altered translation elongation contributes to key hallmarks of aging in the killifish brain. Science (1979). 389, (2025).

79. Solyga, M., Majumdar, A. & Besse, F. Regulating translation in aging: from global to gene-specific mechanisms. EMBO Reports vol. 25 5265–5276 Preprint at 10.1038/s44319-024-00315-2 (2024).

80. Mangleburg, C. G. et al. Integrated analysis of the aging brain transcriptome and proteome in tauopathy. Mol. Neurodegener. 15, (2020).

81. Dick, F., Tysnes, O. B., Alves, G. W., Nido, G. S. & Tzoulis, C. Altered transcriptome-proteome coupling indicates aberrant proteostasis in Parkinson’s disease. iScience 26, (2023).

82. Nixon, R. A. Amyloid precursor protein & endosomal-lysosomal dysfunction in Alzheimer’s disease: Inseparable partners in a multifactorial disease. FASEB Journal 31, 2729–2743 (2017).

83. Colacurcio, D. J. & Nixon, R. A. Disorders of lysosomal acidification—The emerging role of v-ATPase in aging and neurodegenerative disease. Ageing Res. Rev. 32, 75–88 (2016).

84. Lee, S., Sato, Y. & Nixon, R. A. Lysosomal proteolysis inhibition selectively disrupts axonal transport of degradative organelles and causes an Alzheimer’s-like axonal dystrophy. Journal of Neuroscience 31, 7817–7830 (2011).

85. Hung, C. O. Y. & Livesey, F. J. Altered γ-Secretase Processing of APP Disrupts Lysosome and Autophagosome Function in Monogenic Alzheimer’s Disease. Cell Rep. 25, 3647–3660.e2 (2018).

86. Gowrishankar, S. et al. Massive accumulation of luminal protease-deficient axonal lysosomes at Alzheimer’s disease amyloid plaques. Proc. Natl. Acad. Sci. U. S. A. 112, E3699–E3708 (2015).

87. Lee, J. H. et al. Faulty autolysosome acidification in Alzheimer’s disease mouse models induces autophagic build-up of Aβ in neurons, yielding senile plaques. Nat. Neurosci. 25, 688–701 (2022).

88. Keller, J. N., Hanni, K. B. & Markesbery, W. R. Impaired proteasome function in Alzheimer’s disease. J. Neurochem. 75, 436–439 (2000).

89. Tseng, B. P., Green, K. N., Chan, J. L., Blurton-Jones, M. & LaFerla, F. M. Aβ inhibits the proteasome and enhances amyloid and tau accumulation. Neurobiol. Aging 29, 1607–1618 (2008).

90. Ribeiro, F. C. et al. Synaptic proteasome is inhibited in Alzheimer’s disease models and associates with memory impairment in mice. *Commun*. Biol. 6, (2023).

91. Koo, B. G., et al. Association of circulating proteasome activity with Alzheimer’s pathology and cognitive functions in APOE ε4 carriers. Alzheimer’s Research and Therapy 18, (2026).

92. Jin, Y. et al. APOE4 exacerbates α-synuclein seeding activity and contributes to neurotoxicity in Alzheimer’s disease with Lewy body pathology. Acta Neuropathol. 143, 641–662 (2022).

93. Wennberg, A. M. et al. Association of Apolipoprotein e ɛ4 with Transactive Response DNA-Binding Protein 43. JAMA Neurol. 75, 1347–1354 (2018).

94. Josephs, K. A., Tsuboi, Y., Cookson, N., Watt, H. & Dickson, D. W. Apolipoprotein E 4 Is a Determinant for Alzheimer-Type Pathologic Features in Tauopathies, Synucleinopathies, and Frontotemporal Degeneration.

95. Shvetcov, A. et al. APOE ε4 carriers share immune-related proteomic changes across neurodegenerative diseases. Nat. Med. 31, 2590–2601 (2025).

96. Takahashi, K. & Yamanaka, S. Induction of Pluripotent Stem Cells from Mouse Embryonic and Adult Fibroblast Cultures by Defined Factors. Cell 126, 663–676 (2006).

97. Smith, T., Heger, A. & Sudbery, I. UMI-tools: Modeling sequencing errors in Unique Molecular Identifiers to improve quantification accuracy. Genome Res. 27, 491–499 (2017).

98. Dobin, A. et al. STAR: Ultrafast universal RNA-seq aligner. Bioinformatics 29, 15–21 (2013).

99. Martin, M. Cutadapt removes adapter sequences from high-throughput sequencing reads. EMBnet. J. 17, 10–12 (2011).

100. Langmead, B. & Salzberg, S. L. Fast gapped-read alignment with Bowtie 2. Nat. Methods 9, 357– 359 (2012).

101. Liu, D. Algorithms for efficiently collapsing reads with Unique Molecular Identifiers. PeerJ 7, (2019).

102. Navickas, A. et al. An mRNA processing pathway suppresses metastasis by governing translational control from the nucleus. Nat. Cell Biol. 25, 892–903 (2023).

103. Love, M. I., Huber, W. & Anders, S. Moderated estimation of fold change and dispersion for RNA-seq data with DESeq2. Genome Biol. 15, (2014).

104. Lawrence, M. et al. Software for Computing and Annotating Genomic Ranges. PLoS Comput. Biol. 9, (2013).

105. Yates, A. D. et al. Ensembl 2020. Nucleic Acids Res. 48, D682–D688 (2020).

106. Durinck, S., Spellman, P. T., Birney, E. & Huber, W. Mapping identifiers for the integration of genomic datasets with the R/ Bioconductor package biomaRt. Nat. Protoc. 4, 1184–1191 (2009).

107. James Kent, W., et al. The human genome browser at UCSC. Genome Res. 12, 996–1006 (2002).

108. Gardiner-Garden’, M. & Frommer1v2, M. CpG Islands in Vertebrate Genomes. J. Mol. Biol vol. 196 (1987).

109. Korotkevich, G. et al. Fast gene set enrichment analysis. biorXiv 060012v3 (2021) doi:10.1101/060012.

110. Liberzon, A. et al. The Molecular Signatures Database Hallmark Gene Set Collection. Cell Syst. 1, 417–425 (2015).

111. Yu, G., Wang, L. G., Han, Y. & He, Q. Y. ClusterProfiler: An R package for comparing biological themes among gene clusters. OMICS 16, 284–287 (2012).

112. Xie, Z. et al. Gene Set Knowledge Discovery with Enrichr. Curr. Protoc. 1, (2021).

113. Chen, E. Y., et al. Enrichr: Interactive and Collaborative HTML5 Gene List Enrichment Analysis Tool. http://amp.pharm.mssm.edu/Enrichr. (2013).

114. Kuleshov, M. V. et al. Enrichr: a comprehensive gene set enrichment analysis web server 2016 update. Nucleic Acids Res. 44, W90–W97 (2016).

115. Carlson, A. M., et al. Package ‘GenomicFeatures’. BSgenome.Hsapiens.UCSC 1–79 Preprint at https://bioconductor.org/packages/GenomicFeatures (2021).

116. Pages, H., Aboyoun, P., Gentleman, R. & DebRoy, S. Biostrings: Efficient manipulation of biological strings. Preprint at https://bioconductor.org/packages/Biostrings (2025).

117. Morgan, M., Pages, H., Obenchain, V. & Hayden, N. Rsamtools: Binary alignment (BAM), FASTA, variant call (BCF), and tabix file import. Preprint at https://bioconductor.org/packages/Rsamtools.

118. Sharp, P. M. & Li, W. H. The codon adaptation index-a measure of directional synonymous codon usage bias, and its potential applications. Nucleic Acids Res. 15, 1281–1295 (1987).

119. Schmidt, E. K., Clavarino, G., Ceppi, M. & Pierre, P. SUnSET, a nonradioactive method to monitor protein synthesis. Nat. Methods 6, 275–277 (2009).

120. R Core Team. R: A language and environment for statistical computing. R Foundation for Statistical Computing, Vienna, Austria. https://www.R-project.org/ (2021).

121. Schindelin, J., et al. Fiji: An open-source platform for biological-image analysis. Nature Methods vol. 9 676–682 Preprint at 10.1038/nmeth.2019 (2012).

