## Supplemental Figures for "APOE4 disrupts the central dogma by arresting neuronal proteome dynamics"

### Document S1

#### FIGURE TITLES AND LEGENDS

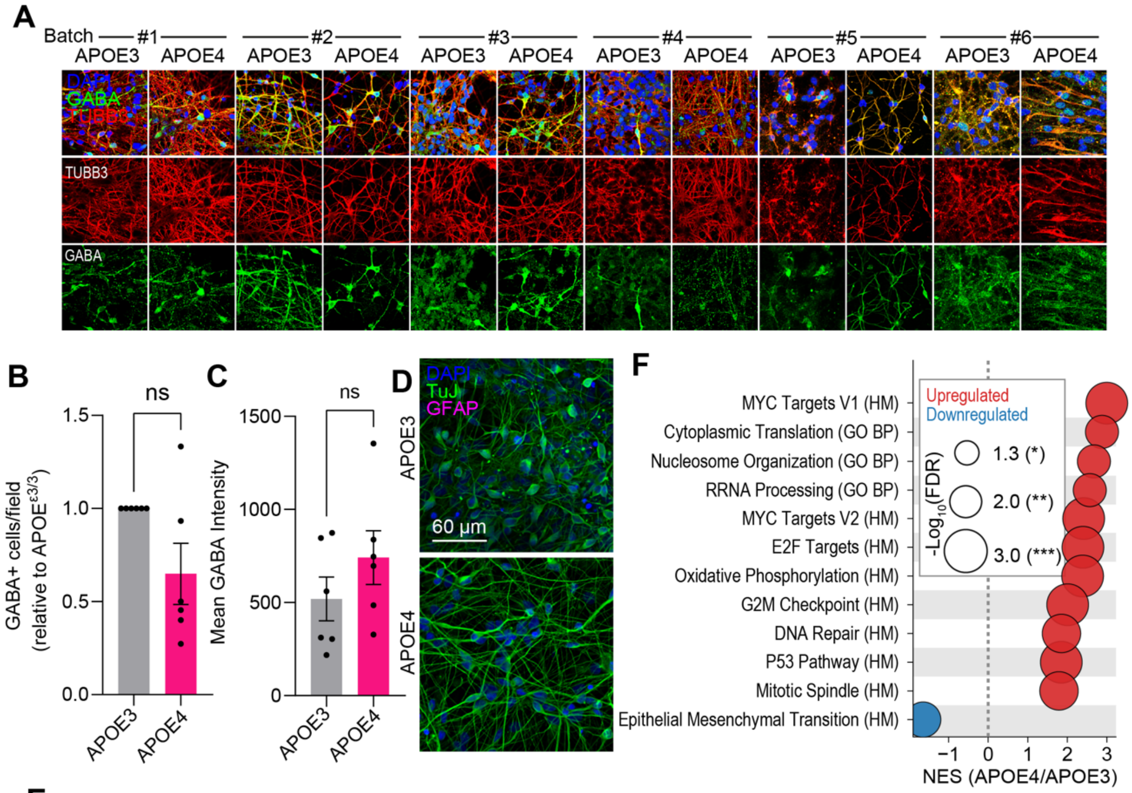


**E** ENCODE and ChEA Consensus TFs from ChIP-X (Ma'ayan Lab Enrichr)

| Transcription factor | Direction | Function | Targets | P value | Combined score |
|---|---|---|---|---|---|
| E2F1 (ChEA) | Up | Activator | TCEAL8, PRPS1, PRDX4, SMS, CHM, SMC1A, PJA1 | $5.86 \times 10^{-3}$ | 18.3 |
| TCF3 (ChEA) | Up | Repressor | TCEAL8, SH3KBP1, DCC, FUNDC1, SMS, HIST1H1A, HS6ST2 | $1.33 \times 10^{-2}$ | 13.0 |
| SOX2 (ChEA) | Up | Repressor | PRPS2, FUNDC1, PHACTR3, MTUS1, HIST1H1A, HS6ST2 | $1.36 \times 10^{-2}$ | 14.3 |
| ZBTB33 (ENCODE) | Up | Variable | HIST1H2BM, PRDX4, HIST1H2AJ, TAB3, SMC1A | $3.27 \times 10^{-2}$ | 10.3 |
| SUZ12 (ChEA) | Down | Repressor | SPON1, SHH, DPP6, LHX6, NKX2-4, POU3F4 | $3.67 \times 10^{-3}$ | 28.2 |

**Figure S1. Differentiation characterization by immunofluorescence and transcript enrichment**

(**A**) Immunofluorescence of DIV49 iPSC-derived neurons, indicating GABAergic interneurons (GABA), total neurons (TUBB3/TuJ), and total cells (DAPI). Column groups indicate separate differentiation batches.

(**B** and **C**) Figure panels associated with Figure 1B-C. Quantification of interneuron density, indicating the relative density of GABA$^{+}$ interneurons compared to apoE3 controls (B), and mean cellular GABA immunofluorescence intensity per cell, in arbitrary fluorescence units (C). n ≥ 5 biological and technical replicates from independent differentiations.

(**D**) Immunofluorescence of DIV35 iPSC-derived neurons, showing immunoreactivity for the neuronal marker TuJ and the astrocyte marker GFAP.

(**E**) Transcription factor enrichment of subset upregulated and downregulated transcripts genes (adj.p<0.05) as determined by RNA-sequencing (associated with Figure panel 1F) among ENCODE/ChEA consensus transcription factors (Ma'ayan Lab Enrichr). The top five most enriched transcription actors, ranked by statistically significance, are shown.

(**F**) Transcript gene-set enrichment analysis (GSEA) results of up- and downregulated transcripts (adj.p<0.05) among the Hallmark (HM) and Gene Ontology Biological Process (GO BP) gene sets.

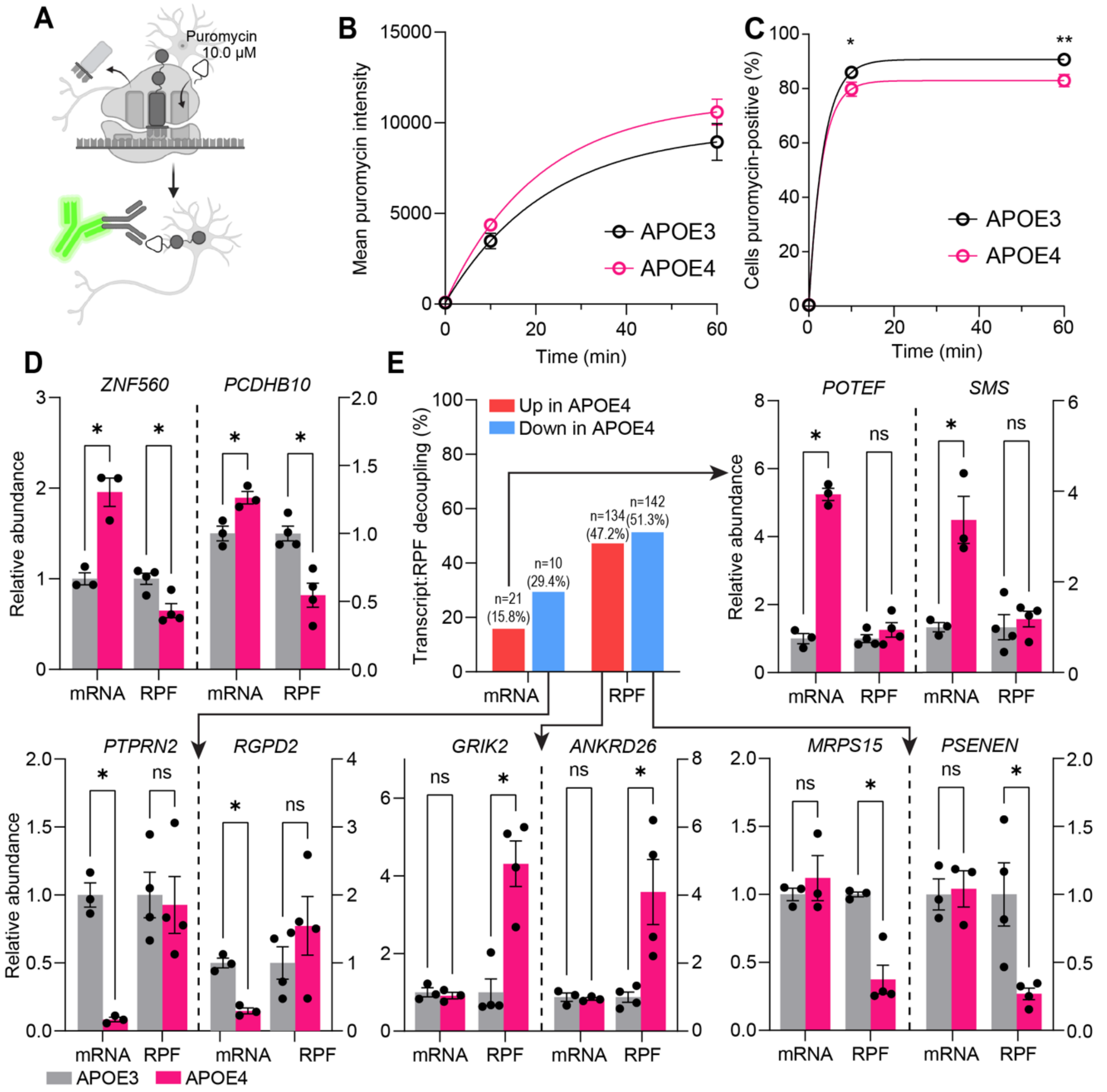


#### Figure S2. The influence of apoE4 on ribosomal translation in apoE4 neurons

(**A**) Schematic of the SUrface SEnsing of Translation (SuNSET) assay performed in iPSC-derived neurons, indicating how puromycin (white triangle) is incorporated into nascent peptides (grey circles), and detected by a fluorophore-coupled anti-puromycin antibody.

(**B** and **C**) Flow cytometry-based analyses indicating the average cellular puromycin intensity (B) and proportion neurons incorporating puromycin (C) over time, dependent on apoE genotype. n = 3 biological and technical replicates from independent differentiations.

(**D**) Bar plots showing of the two discordant genes across transcript and RPF layers (adj.p<0.05) in the form up mRNA upregulation (RNA-seq) and decreased ribosome occupancy (ribo-seq) in apoE4 neurons.

(**E**) Example plots for transcript:RPF decoupling. Summary bar plot indicates the fraction of dysregulated (adj.p) events within each category, with most changes affecting RPF/translation alone. Arrows point towards bar plots that represent such discordant changes for each of the illustrated categories.

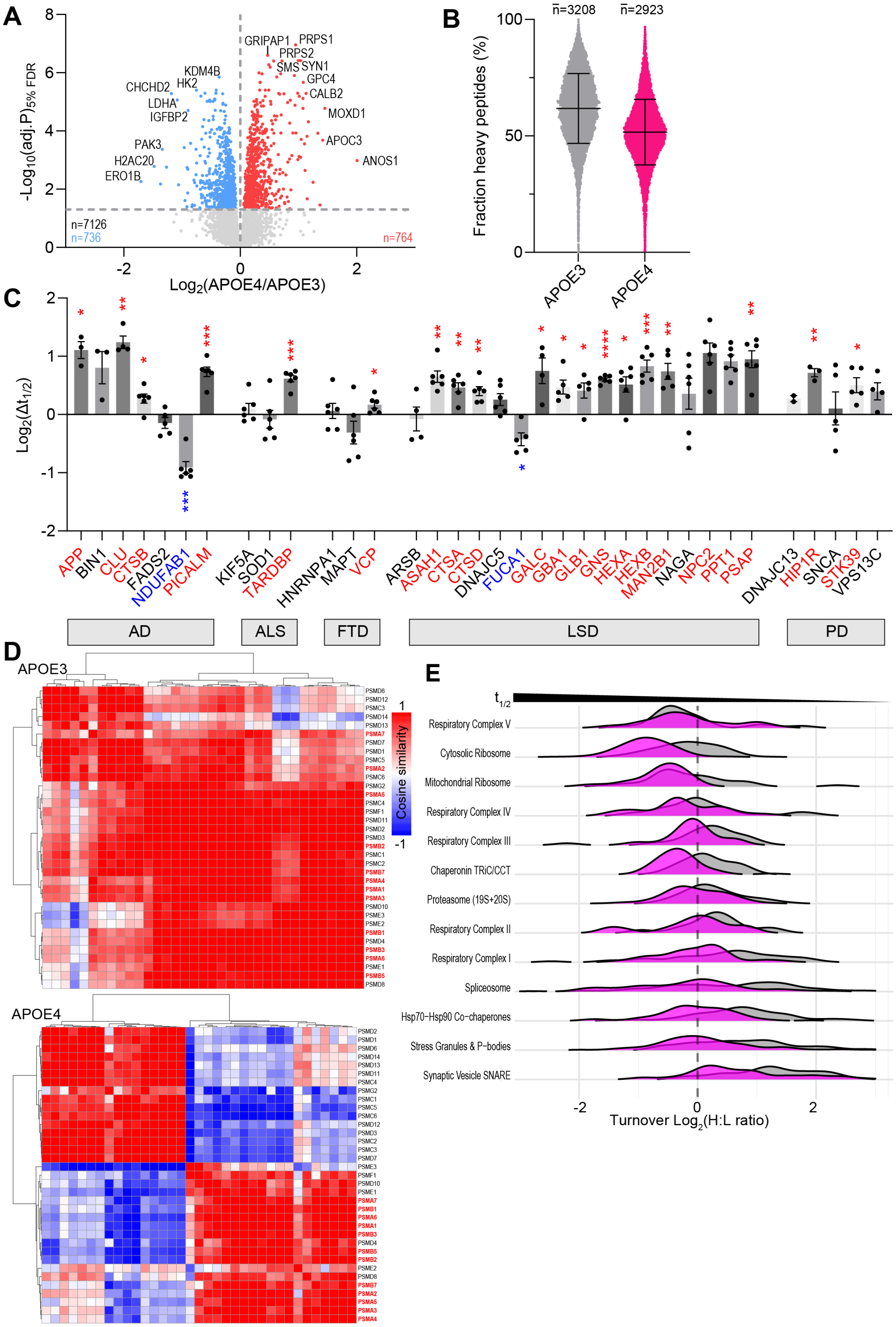


**Figure S3. Protein turnover changes associated with neuronal apoE4**

(**A**) Volcano plot comparing protein abundance between apoE4 and apoE3 iPSC-derived neurons normalized to median protein intensity. Associated with Figure 3A. Up- and down-regulated data points (adj. $p<0.05$) are colored red and blue, respectively. n = 6 biological and technical replicates from independent differentiations. See Table S3B.

(**B**) Fraction heavy (H) protein (protein group) relative to total protein (light/L + heavy) in apoE3 and apoE4 samples following 2 weeks of heavy $^{13}C_6$ L-Lysine incorporation. Each data point represents an individual protein. Line indicates mean, error bars indicate standard deviation. n ≥ 5 biological and technical replicates from independent differentiations.

(**C**) $Log_2$-transformed half-life change ($Log_2\Delta t_{1/2}$) in apoE4 neurons relative to apoE3 neurons of risk factors associated with the neurodegenerative proteinopathies Alzheimer's disease (AD), amyotropic lateral sclerosis (ALS), frontotemporal dementia (FTD), lysosomal storage diseases (LSDs), and Parkinson's disease (PD). Gene symbols and significance labels are colored red for extended half-lives, black for no change in half-life, and blue for decreased half-lives by apoE4. Each data point represents protein half-life changes in individual differentiations, relative to matched apoE3 controls. n ≥ 5 biological and technical replicates from independent differentiations.

(**D**) Representative cosine similarity matrices showing the relationships between protein turnover profiles within proteasome complexes of apoE3 (top) and apoE4 (bottom) samples. Proteins are ordered by hierarchical clustering, with red indicating greater similarity in turnover profiles and blue indicating lower similarity. n ≥ 5 biological and technical replicates from independent differentiations.

(**E**) Ridge plots showing the distribution of pairwise cosine similarity values for proteins within each indicated protein complex in apoE3 and apoE4 samples. Rightward shifts indicate greater overall similarity in protein turnover profiles within a complex (coordinated turnover), whereas leftward shifts indicate greater divergence (dissociated turnover). n ≥ 5 biological and technical replicates from independent differentiations.

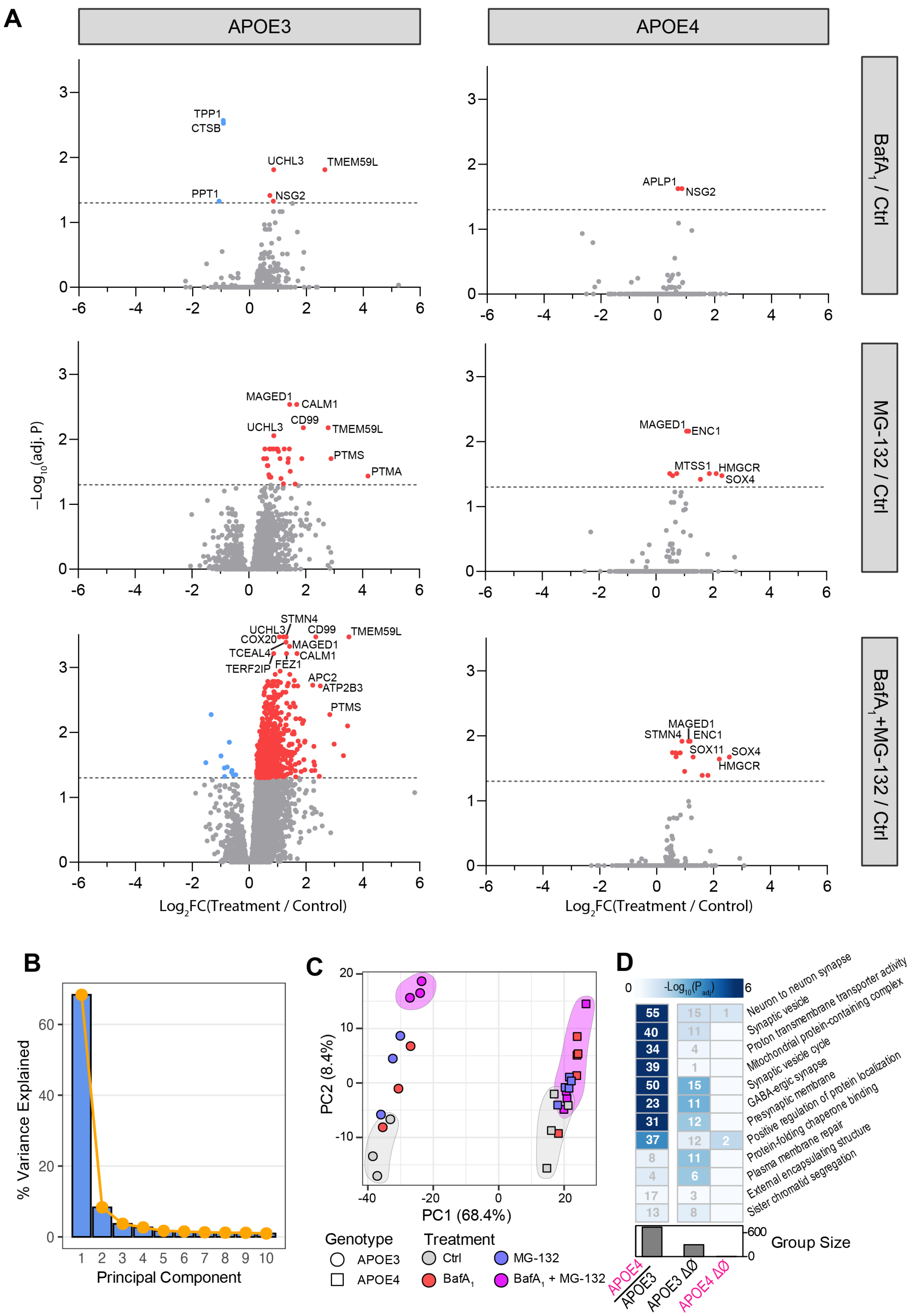


**Figure S4. Genotype-dependent proteome response to protein clearance inhibition**

(**A**) Volcano plots comparing housekeeping-normalized protein abundance following 3 h treatment with lysosomal ($BafA_1$, top), proteasomal (MG-132, middle), combined lysosomal and proteasomal (bottom) clearance inhibition in apoE3 (left) and apoE4 (right) neurons. Associated with Figure 4D-E. Up- and downregulated data points (adj. $p<0.05$) are colored red and blue, respectively. $n \geq 3$ biological and technical replicates from independent differentiations. See Table S4A.

(**B**) Relative variance explained by each principal component of the clearance inhibitor dataset, across all treatment and genotype conditions.

(**C**) Principal component analysis (PCA) of proteomic profiles from apoE3 and apOE4 neurons under control conditions, or following treatment with $BafA_1$, MG-132, or the combination of both clearance inhibitors. Shaded regions indicate the range of control (grey) and $BafA_1$ + MG-132 co-treatment (purple) for each genotype.

(**D**) Functional enrichment analysis of proteins significantly altered in apoE4 relative to apoE3 neurons, or following combined clearance inhibition (ΔØ) in apoE3 and apoE4 neurons. Heat-map shows the most significantly enriched gene ontology terms, with inset numbers indicating the number of associated proteins.

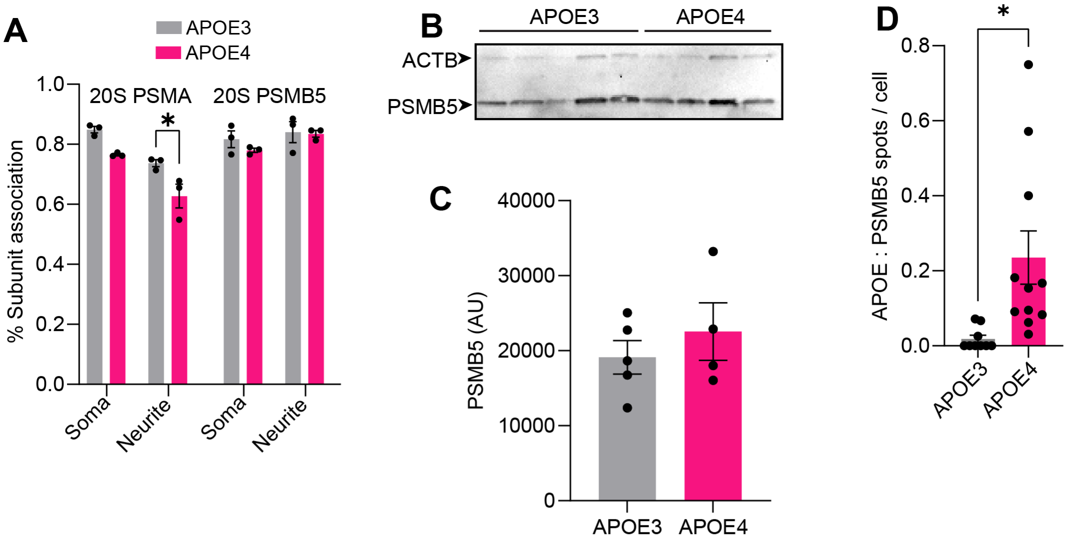


**Figure S5. Proteasome composition and protein associations in apoE3 and apoE4 neurons**

(**A**) Quantification of PSMA1 and PSMB5 association with the 20S proteasome in somatic and neuritic compartments of apoE3 and apoE4 neurons, determined by Manders' colocalization analysis. Associated with Figure 5A. n = 3 biological replicates from independent differentiations.

(**B** and **C**) Representative immunoblot (B) and associated densitometry quantification (C) showing PSMB5 protein abundance in lysates from apoE3 and apoE4 neurons used for the proteasome activity assay, with ACTB as the loading control. Associated with Figures 5D-E. n = 4 biological replicates from independent differentiations.

(**D**) Quantification of APOE:PSMB5 proximity ligation assay puncta per cell in apoE3 and apoE4 neurons. Associated with Figure 5H-I. n = 3 biological replicates from independent differentiations.

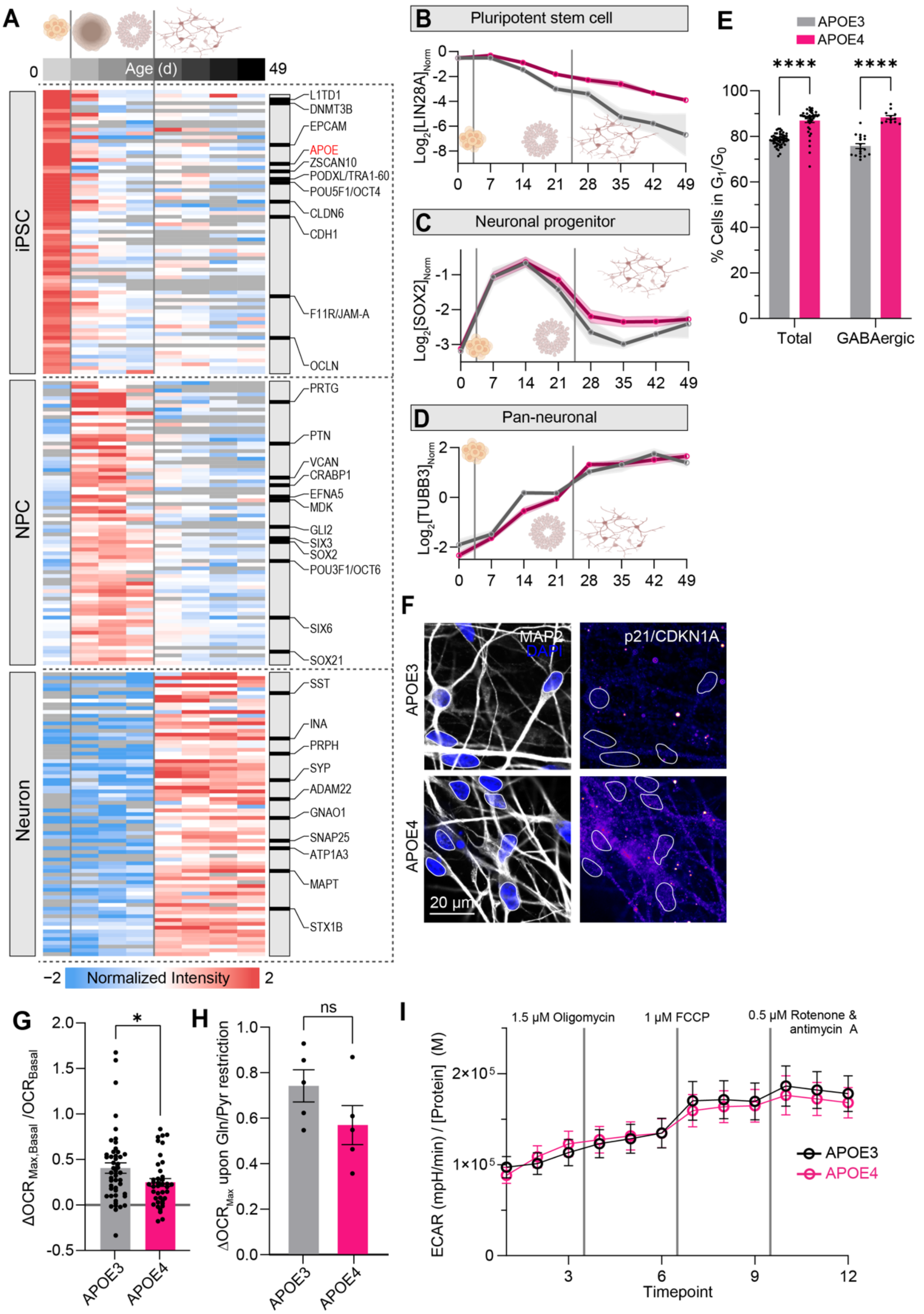


**Figure S6. Neuronal differentiation and metabolic characterization of apoE4 neurons**

(**A**) Heat-map showing temporal protein abundance changes during differentiation of iPSCs into neurons, irrespective of apoE genotype. Proteins are grouped according to pluripotent stem cell (iPSC; DIV0), neuronal progenitor cell (NPC; DIV7-DIV21), and neuronal stage (DIV21-DIV49) enrichment over other differentiation stages. APOE was enriched during the iPSC stage, and is highlighted in red. Other established stage-specific differentiation markers are highlighted by gene symbol annotation.

(**B**-**D**) Temporal protein abundance profiles of representative markers for pluripotent stem cells (LIN28A; B), neuronal progenitor cells (SOX2; C), and neurons (TUBB3; D) throughout the differentiation of apoE3 and apoE4 cultures. Dots represent proteomic measurements per stage for apoE3 (grey) or apoE4 (magenta) neurons, alongside connecting lines depicting stage-associated changes in protein abundance. Vertical lines indicate stage breaks. Shaded areas indicate the standard error of the mean (SEM). Associated with Figure 6A. $n \geq 3$ biological and technical replicates from independent differentiations.

(**E**) Flow cytometric analysis of the proportion of total (left) and GABAergic (right) DIV49 neurons in the $G_1/G_0$ cell cycle phase, determined by DAPI DNA population analysis. $n = 3$ biological and technical replicates from independent differentiations.

(**F**) Representative immunofluorescence images of MAP2-positive neurons stained for the senescence marker CDKN1A/p21 in DIV49 neurons. Associated with Figure 6I. $n = 3$ biological and technical replicates from independent differentiations.

(**G**) Quantification of spare respiratory capacity in DIV49 neurons, expressed as the change in oxygen consumption rate (ΔOCR) following FCCP treatment relative to basal respiration. Associated with Figure 6J. $n = 3$ biological and technical replicates from independent differentiations.

(**H**) Quantification of glycolytic reserve in DIV49 neurons, expressed as the change in extracellular acidification rate (ΔECAR) during sequential treatment with oligomycin and FCCP. $n = 3$ biological and technical replicates from independent differentiations.

(**I**) Glycolytic stress test of iPSC-derived neurons showing extracellular acidification rate (ECAR) changes during sequential treatment with oligomycin, FCCP, and rotenone/antimycin A. $n = 3$ biological and technical replicates from independent differentiations.
